# Engineered cytokine/antibody fusion proteins improve delivery of IL-2 to pro-inflammatory cells and promote antitumor activity

**DOI:** 10.1101/2023.05.03.539272

**Authors:** Elissa K. Leonard, Jakub Tomala, Joseph R. Gould, Michael I. Leff, Jian-Xin Lin, Peng Li, Mitchell J. Porter, Eric R. Johansen, Ladaisha Thompson, Shanelle D. Cao, Tereza Henclova, Maros Huliciak, Ondřej Vaněk, Marek Kovar, Warren J. Leonard, Jamie B. Spangler

## Abstract

Progress in cytokine engineering is driving therapeutic translation by overcoming the inherent limitations of these proteins as drugs. The interleukin-2 (IL-2) cytokine harbors great promise as an immune stimulant for cancer treatment. However, the cytokine’s concurrent activation of both pro-inflammatory immune effector cells and anti-inflammatory regulatory T cells, its toxicity at high doses, and its short serum half-life have limited clinical application. One promising approach to improve the selectivity, safety, and longevity of IL-2 is complexation with anti-IL-2 antibodies that bias the cytokine towards the activation of immune effector cells (i.e., effector T cells and natural killer cells). Although this strategy shows therapeutic potential in preclinical cancer models, clinical translation of a cytokine/antibody complex is complicated by challenges in formulating a multi-protein drug and concerns about complex stability. Here, we introduce a versatile approach to designing intramolecularly assembled single-agent fusion proteins (immunocytokines, ICs) comprising IL-2 and a biasing anti-IL-2 antibody that directs the cytokine’s activities towards immune effector cells. We establish the optimal IC construction and further engineer the cytokine/antibody affinity to improve immune biasing function. We demonstrate that our IC preferentially activates and expands immune effector cells, leading to superior antitumor activity compared to natural IL-2 without inducing toxicities associated with IL-2 administration. Collectively, this work presents a roadmap for the design and translation of immunomodulatory cytokine/antibody fusion proteins.

**One Sentence Summary:** We developed an IL-2/antibody fusion protein that expands immune effector cells and shows superior tumor suppression and toxicity profile versus IL-2.

## INTRODUCTION

Interleukin-2 (IL-2) is a multifunctional cytokine that is produced primarily by T cells and is responsible for coordinating numerous essential activities in a variety of immune cells. IL-2 is vital for inducing proliferation of both pro-inflammatory immune effector cells (Effs, i.e., CD4^+^ and CD8^+^ effector T cells and natural killer [NK] cells) and anti-inflammatory regulatory T (T_reg_) cells (*1, 2*). These activities make IL-2 an alluring candidate for therapeutic immunomodulation in diseases ranging from cancer to autoimmune disorders (*3, 4*). Unfortunately, stimulation of both pro- and anti-inflammatory cells has limited the cytokine’s efficacy in the treatment of cancer, and for the 5-10% of patients whose cancer does respond to therapy, high doses of IL-2 are required for sustained tumor regression (*5, 6*). Unfortunately, high dose IL-2 is frequently accompanied by toxicities (most prominently vascular leak syndrome) that can sometimes be fatal (*7*), limiting the cytokine’s clinical application. Moreover, IL-2 also has an extremely short serum half-life (<5 min), which has further complicated its therapeutic use (*8*).

Fusing the IL-2 cytokine to a tumor-targeting antibody to form an “immunocytokine” is a drug development approach that has been adopted to mitigate IL-2 toxicity (*9, 10*). These molecules promote a more spatially targeted immune response while also increasing serum half-life via neonatal Fc receptor (FcRN)-mediated recycling (*11*), thereby attenuating the systemic toxicity that is observed for IL-2 alone. Other trials have successfully improved therapeutic efficacy by combining IL-2 with another therapeutic agent, such as traditional chemotherapeutics (*12*) or immune checkpoint inhibitors (*13–15*).

In addition to combining IL-2 with other treatments, strategies have been developed to disrupt or bias the interaction between IL-2 and its cognate receptor, in order to achieve a greater therapeutic effect at lower, less toxic doses. These biasing strategies have shown the potential to selectively stimulate and expand particular immune cell subsets for targeted disease therapy. IL-2 signaling occurs through the IL-2 receptor- (IL-2Rβ) and the common gamma chain (γ_C_) on both pro- and anti-inflammatory immune cells (*2, 4, 13, 16*). IL-2-mediated heterodimerization of IL-2Rβ and γ_C_ leads to dimerization of the intracellular domains of these receptor chains, which activates the Janus kinase-signal transducer and activator of transcription (JAK-STAT) pathway. JAK-STAT signaling ultimately leads to phosphorylation of signal transducer and activator of transcription 5 (STAT5), which induces diverse gene expression programs that dictate cellular behavior (*17*). Though the IL-2 receptor signaling subunits are identical for all cells, immunosuppressive T_regs_ have much higher basal expression levels of the non-signaling IL-2Rα subunit compared to Effs, and this subunit is primarily responsible for binding and retaining IL-2 to facilitate interaction with IL-2Rβ and γ_C_ (*16*). IL-2Rα, IL-2Rβ, and γ_C_ together form a heterotrimeric receptor, which has a 100-fold higher affinity than the IL-2Rβ/γ_C_ heterodimeric receptor. Thus, cells that express IL- 2Rα (i.e., T_regs_) have far greater sensitivity, responsiveness, and ability to consume IL-2 compared to those that do not (i.e., naïve Effs) (*1, 18*).

By disrupting the interaction between IL-2 and IL-2Rα, the competitive advantage for IL-2Rα- expressing cells can be eliminated, and the pro-inflammatory Effs that promote immune activation can be more potently and specifically stimulated. This approach has been employed to enhance IL-2’s activity as an anti-cancer agent. One such strategy involves the design of mutant IL-2 variants (termed muteins) that have reduced or fully abrogated interaction with IL-2Rα (*19–21*). Alternatively, monoclonal antibodies that target the IL-2 cytokine and block the IL-2Rα-binding site have been identified, and complexes that combine IL-2 with these antibodies have been shown to preferentially stimulate Effs compared to IL-2 alone (*22*). Selective expansion of Effs was accompanied by corresponding improvements in antitumor efficacy for both mouse IL-2, using monoclonal antibody S4B6 (*22–26*), and for human IL-2, using monoclonal antibodies MAB602 (*23, 24, 27*), NARA1 (*27, 28*), and TCB2 (*29*).

This study has blended the immunocytokine and cytokine-targeted antibody approaches by designing a single-agent fusion protein comprised of IL-2 and MAB602 (referred to henceforth as 602), which acts as an intramolecularly assembled immunocytokine (IC). Tethering IL-2 to the antibody stabilizes the cytokine/antibody complex, reducing off-target activation, and we show that this tethering alone resulted in a dramatic improvement in biased stimulation of Effs. Moreover, fusion of IL-2 to the antibody improves the pharmacokinetics of the therapy by preventing cytokine clearance through covalent linkage to the antibody. Furthermore, as a single agent therapeutic, our IC eliminates questions of IL-2/antibody complex formulation and stoichiometric optimization, mitigating regulatory obstacles to clinical translation. We applied molecular evolution approaches to isolate a variant of 602 that enhanced the biased expansion of Effs over T_regs_, and we showed that our evolved variant further accentuated immune effector cell bias both in vitro and in vivo, leading to robust inhibition of tumor growth in mouse models of cancer. Altogether, this work constitutes a major step forward in advancing IL-2-biasing antibodies as viable candidates for clinical translation.

## RESULTS

### Modifying linker length optimizes 602 immunocytokine production

To combine the potency of cytokines with the pharmaceutically favorable properties of antibodies, the IL-2 cytokine was fused to the IL-2Rα-competitive anti-IL-2 antibody, 602 (Fig. 1A) (*23, 24*), to create an intramolecularly assembled IC. The C-terminus of IL-2 was tethered to the N-terminus of the 602 antibody light chain by a (G_4_S)_n_ linker, which was 10, 15, 25, or 35 amino acids in length (denoted 602 IC LN10, 602 IC LN15, 602 IC LN25, or 602 IC LN35, respectively). All versions of 602 IC migrated at the expected molecular weights via SDS-PAGE analysis (Fig. 1B); however, size-exclusion chromatography (SEC) traces revealed distinctive elution profiles, with each IC partitioning between three peaks (Fig. 1C). The first two peaks correspond to proteins with much larger molecular weights than would be expected for a single 602 IC (∼180 kDa). These two peaks likely correspond to oligomers of multiple ICs that have exchanged one or both of their IL- 2 moieties with another IC rather than binding intramolecularly to the 602 antibody to which they are tethered (Fig. 1C). The first two peaks are dominant in 602 IC LN10 and 602 IC LN15, which is likely due to the linker being of inadequate length to accommodate intramolecular cytokine/antibody binding. Indeed, increasing the length of the linker to 25 amino acids introduced a third peak corresponding to the expected molecular weight for a single 602 IC (Fig. 1C, fig. S1A), although more than half of the 602 IC LN25 eluted in the first two peaks (Fig. 1, C and D, fig. S1, B and C). The fractions collected from all three SEC peaks of 602 IC LN25 showed identical migration on an SDS-PAGE gel under non-reducing conditions (fig. S1D), indicating that the assembly of the higher-order oligomers was reversible. Extending the linker to 35 amino acids resulted in a molecule (602 IC LN35) that eluted predominantly (∼85%) in the third peak (Fig. 1, C and D). In producing multiple preps of 602 IC LN35, we observed that a greater percentage of the protein eluted in the second peak rather than the third when the protein was more concentrated (>5 µM) prior to SEC separation (fig. S1E). Nonetheless, ≥75% of 602 IC LN35 eluted in the third peak, even at concentrations greater than 20 µM. Taken together, these results demonstrated that extending the linker length within the IC enhanced intramolecular assembly of the component IL- 2 and anti-cytokine antibody, allowing for isolation of the pure, monomeric IC.

**Fig. 1.**
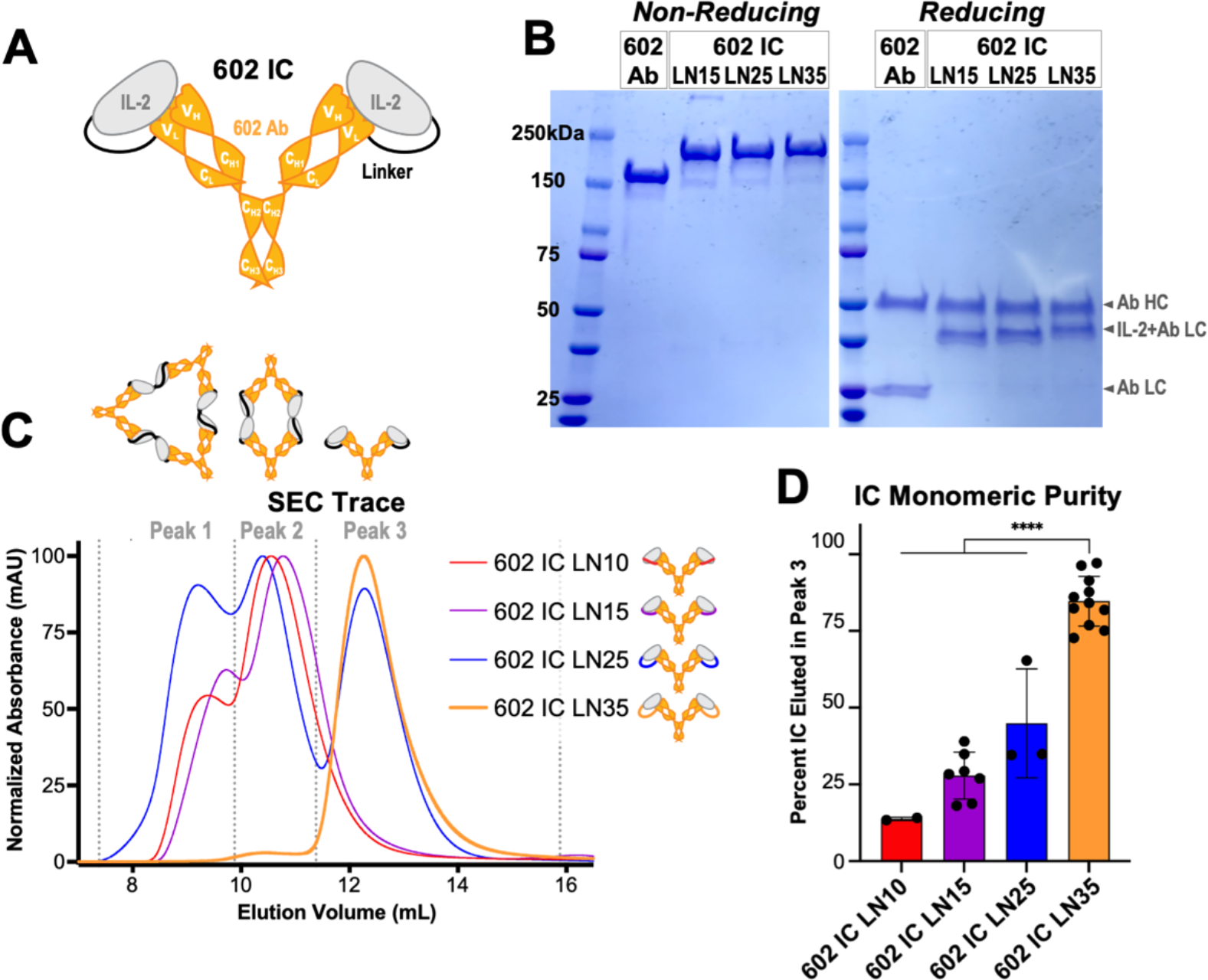
Optimization of immunocytokine design. (A) Schematic of the IL-2 cytokine/602 antibody (Ab) single-chain fusion protein (immunocytokine, IC), wherein the C-terminus of the cytokine is tethered to the N-terminus of the Ab light chain (LC) via a flexible linker. Heavy chain (HC) and LC variable and constant domains are labeled. **(B)** 602 Ab, 602 IC LN15, 602 IC LN25, and 602 IC LN35 migrate at the expected sizes by SDS-PAGE, under non-reducing (145 kDa for Ab and ∼180 kDa for 602 ICs) and reducing (49 kDa for HC, 24 kDa for Ab LC, and ∼41 for IC LC) conditions. **(C)** Representative size-exclusion chromatography (SEC) traces show the relative distribution of 4 linker length variants of 602 (602 IC LN10, 602 IC LN15, 602 IC LN25, and 602 IC LN35) amongst three peaks, corresponding to either multi-IC oligomers (peaks 1 and 2) or monomeric IC (peak 3). **(D)** The average percentage of each IC that eluted in the third peak based on the area under the curve is shown. Data represent mean ± SD from at least 2, and up to 11, purifications. Statistical significance is noted only for comparisons to 602 IC LN35. ****P<0.0001 by one-way ANOVA with Tukey’s multiple comparison test.

### Extended intramolecular linker optimizes 602 immunocytokine function

To assess the function of the species contained in each of the three SEC peaks, the peaks were separately pooled and concentrated (fig. S1, A and D). Interestingly, the contents of all three peaks exhibited identical binding to IL-2 and IL-2Rβ, as assessed by bio-layer interferometry (BLI) (fig. S1, F and G). The signaling activity of the eluted peaks was compared by measuring phosphorylation of STAT5 as a readout for IL-2-mediated signaling on cytokine-responsive human YT-1 natural killer (NK) cells (*30*). In contrast with the observed binding behavior, we found that the contents of the first two SEC peaks for 602 IC LN25 elicited weaker signaling compared to those of the third peak (fig. S1H). For this reason, all subsequent comparisons used the third peak of the LN25 and LN35 ICs, corresponding to the size of a monomeric IC. As there was no distinct third peak that provided adequate yield for characterization, the second peak of LN15 was used in subsequent comparisons.

To assess functional differences between 602 ICs with various linker lengths, binding interactions with IL-2, IL-2Rα, and IL2Rb were analyzed by BLI. All three 602 ICs showed similar equilibrium binding properties (Fig. 2, A-C, fig. S2, A-C, table S1). Tethering IL-2 to 602 in the ICs reduced exchange with immobilized IL-2 as compared to the IL-2/602 mixed cytokine/antibody complex, principally through deceleration of association rate (fig. S2A, table S1), illustrating the increased complex stability resulting from cytokine/antibody tethering. The stably assembled 602 IC LN35 showed a very similar profile to IL-2/602 complex by analytical ultracentrifugation, which revealed small amounts of dimeric and trimeric species for 602 IC LN35, and potentially dimeric species for IL-2/602 complex that may have resulted from low levels of dimeric IL-2 (fig. S3A). Thermostability measurements conducted by differential scanning fluorometry (DSF) revealed that the melting temperature (T_m_) of 602 IC LN35 was at least 0.5°C higher than that of IL-2/602 complex (fig. S3, B and C, table S2), further illustrating the increase in overall molecular stability for the IC as compared to the complex. As expected, based on the competitive properties of the 602 antibody (*23, 24*), none of the 602 ICs engaged IL-2Rα (Fig. 2B, fig. S2B). The ICs and IL- 2/602 complex showed identical equilibrium binding to IL-2Rβ (Fig. 2C), indicating that the linkers did not interfere with the interaction between IL-2 and IL-2Rβ. Binding to IL-2Rβ was potentiated for ICs and IL-2/602 complex compared to the free cytokine due to bivalency. Notably, 602 ICs with shorter linkers had slower IL-2Rβ association and dissociation rates compared with 602 IC LN35 and IL-2/602 complex (fig. S2C, table S1), possibly due to the presence of oligomeric species.

**Fig. 2.**
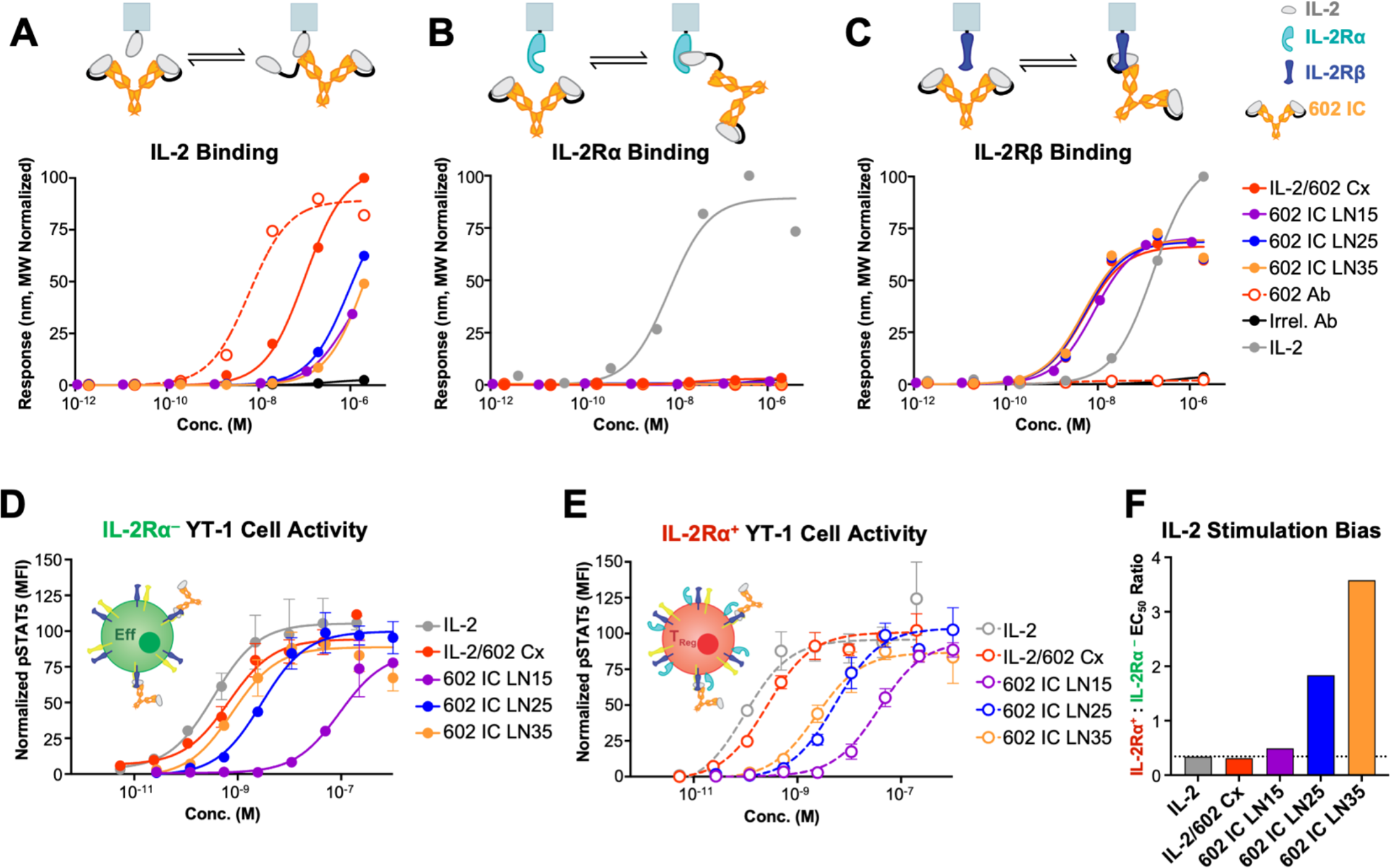
Increasing the linker length within ICs improves stability and enhances immune effector cell bias. (A) Equilibrium bio-layer interferometry (BLI) titrations of soluble IL-2/602 complex (Cx) (2:1 cytokine:antibody molar ratio), 602 antibody (Ab), 602 IC LN15, 602 IC LN25, and 602 IC LN35 against immobilized IL-2. An antibody with irrelevant specificity (Irrel. Ab) is included as a negative control. **(B)** Equilibrium BLI titrations of soluble IL-2, IL-2/602 Cx (2:1), 602 Ab, 602 IC LN15, 602 IC LN25, and 602 IC LN35 against immobilized IL-2Rα. Irrel. Ab is included as a negative control. **(C)** Equilibrium BLI titrations of soluble IL-2, IL-2/602 Cx (2:1), 602 Ab, 602 IC LN15, 602 IC LN25, and 602 IC LN35 binding to immobilized IL-2Rβ. Irrel. Ab is included as a negative control. **(D)** STAT5 phosphorylation response of IL-2Rα^-^ YT-1 human NK cells treated with IL-2, IL-2/602 Cx (1:1), 602 Ab, 602 IC LN15, 602 IC LN25, and 602 IC LN35. Data represent mean ± SD (n=3). **(E)** STAT5 phosphorylation response of IL-2Rα^+^ YT-1 human NK cells treated with IL-2, IL-2/602 Cx (1:1), 602 Ab, 602 IC 602 IC LN15, 602 IC LN25, and 602 IC LN35. Data represent mean ± SD (n=3). **(F)** Ratio of the STAT5 phosphorylation EC_50_ values for IL-2Rα^+^ to IL-2Rα^−^ cells treated with IL-2, IL-2/602 Cx (1:1), 602 Ab, 602 IC LN15, 602 IC LN25, and 602 IC LN35. Higher ratio indicates stronger bias toward IL-2Rα^-^ cell activation.

To assess biased IL-2 signaling of 602 ICs, we compared their relative signaling activity on unmodified, IL-2Rα^-^ YT-1 cells to that on induced, IL-2Rα^+^ YT-1 cells (*31*), which served as surrogates for Effs versus T_regs_, respectively. Signaling activity of 602 IC LN15 on IL-2Rα^-^ Eff-like cells was markedly diminished compared to that of IL-2/602 complex (Fig. 2D, table S3). Activity improved when the linker length was increased to 25, and IL-2 signaling was fully restored for 602 IC LN35. On IL-2Rα^+^ T_reg_-like cells, the signaling activity of 602 IC LN35 was only partially restored compared to that of IL-2/602 complex (Fig. 2E), which is likely a result of increased blockade of the IL-2/IL-2Rα interaction due to enhanced stability of the cytokine/antibody interaction when the two molecules are tethered within the ICs. Thus, signaling of 602 IC LN35 shows enhanced bias towards IL-2Rα^-^ versus IL-2Rα^+^ cells. This Eff-biased IL-2 activity was quantified as the EC_50_ ratio of STAT5 phosphorylation on IL-2Rα^+^ T_reg_-like cells versus IL-2Rα^-^ Eff-like cells (Fig. 2F). IL-2/602 complex and all three 602 ICs improved bias towards Effs compared to IL-2 alone, and 602 IC LN35 mediated the most dramatic improvement in bias. Based on the collective binding and activity studies, 35 amino acids was identified as the optimal linker length; therefore, 602 IC LN35 was used in all subsequent studies and is denoted 602 IC henceforth.

### Engineered 602 IC enhances disruption of IL-2/IL-2Rα interaction

To further bias 602 IC activity towards Effs over T_regs_, we generated an error-prone mutagenic DNA library that randomized the first and third complementarity-determining loops (CDRs) of the variable heavy and light chains of the 602 antibody, expressed in single-chain variable fragment (scFv) format. The library was transformed into competent yeast and evolved against human IL-2 using the yeast surface display directed evolution platform (*32*) through iterative rounds of magnetic-activated cell sorting (MACS) and fluorescence-activated cell sorting (FACS). Later rounds of selection were performed in the presence of excess IL-2Rα to identify clones that successfully outcompeted soluble receptor for IL-2 engagement. After 5 rounds of sorting, the evolved library showed improved binding to IL-2 (Fig. 3A), as well as enhanced competition with the IL-2Rα receptor subunit (Fig. 3B). Clones from the evolved library that showed superior competition with IL-2Rα for IL-2 binding compared to the parent 602 scFv were selected for sequence characterization (fig. S3, D and E). Amongst the sequenced clones, no mutations were observed in the CDR1 of the heavy chain, only one variant contained a mutation in the CDR1 of the light chain, and mutations in CDR3 of the heavy chain were restricted to a single residue. Most mutations were localized to the CDR3 of the light chain, and consensus was observed in mutation of the third residue (F225 [F91 in CDR3L, Kabat numbering]) from phenylalanine to less bulky amino acids.

**Fig. 3.**
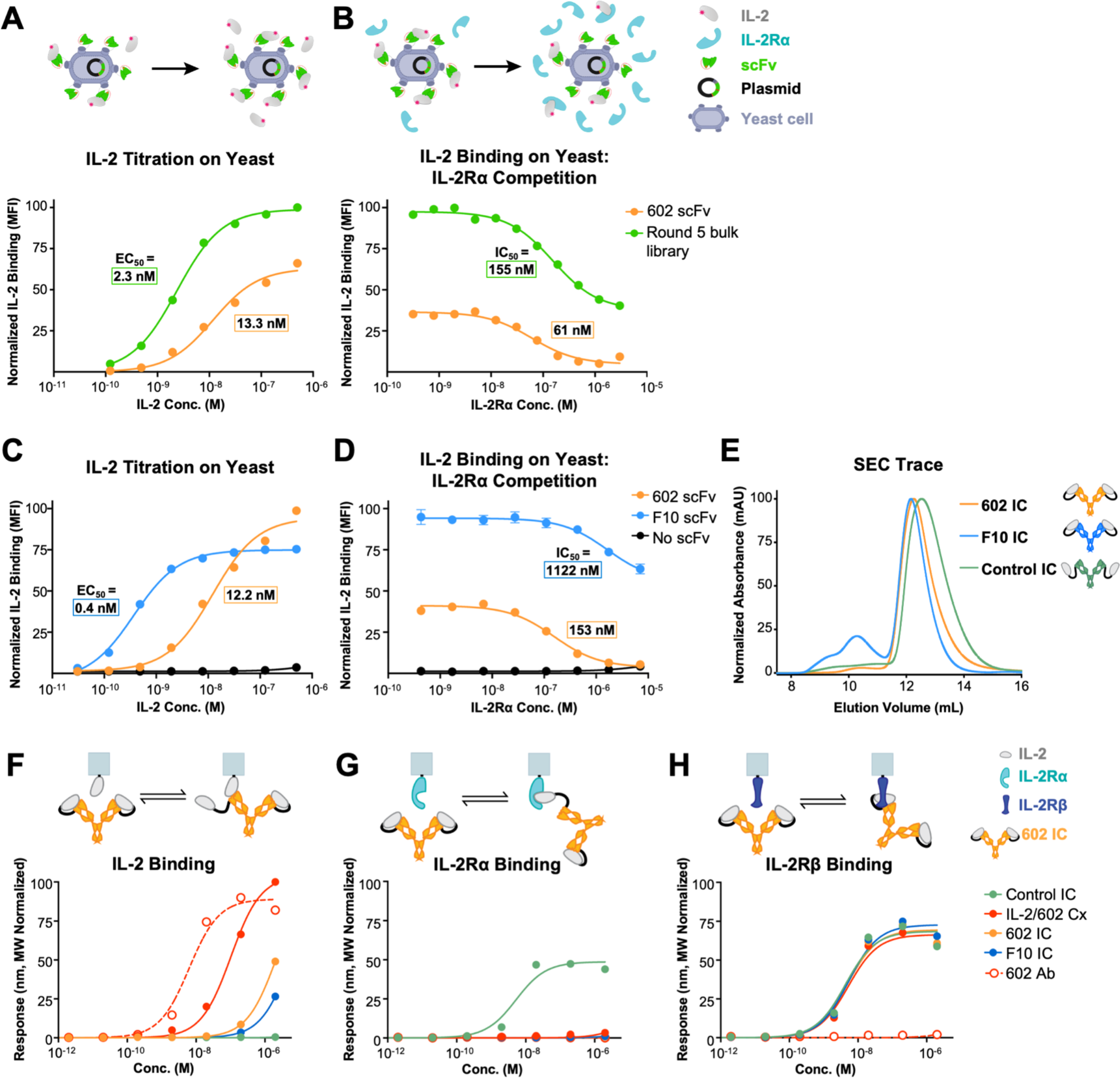
Engineered 602 variants exhibit stronger competition with IL-2Rα compared to the parent antibody. (A, B) IL-2 binding to yeast-displayed 602 scFv compared to engineered 602 scFv variants derived from an evolved error-prone mutagenic library subjected to 5 rounds of selection. IL-2 titrations (A) and IL-2 binding (10 nM) in the presence of various concentrations of IL-2Rα competitor (B) are shown. The pool of variants retains IL-2 binding at concentrations of IL-2Rα that fully blocked the interaction between IL-2 and the parent 602 antibody. **(C, D)** IL- 2 binding to yeast-displayed 602 scFv compared to the 602 scFv variant F10. IL-2 titrations (C) and IL-2 binding (5 nM) in the presence of various concentrations of IL-2Rα competitor (D) are shown. Yeast cells transformed with a plasmid encoding for c-Myc-tagged Aga2 without an scFv were used as a negative control. Data represent mean ± SD (n=3). **(E)** Representative size-exclusion chromatography (SEC) traces show that the majority of IC elutes in the third peak for F10 IC, as well as for Control IC, wherein the cytokine and antibody are not intramolecularly bound. **(F-H)** Equilibrium bio-layer interferometry (BLI) titrations of soluble IL-2/602 complex (Cx) (2:1 cytokine:antibody molar ratio), 602 antibody (Ab), 602 IC, F10 IC, and Control IC against immobilized IL-2 (F), immobilized IL-2Rα (G), and immobilized IL-2Rβ (H).

Selected 602 scFv variants were produced recombinantly in IC LN35 format (fig. S3F). Compared to the parent 602 IC, IC variants exhibited greater sensitivity to concentration. In particular, the percentage of protein eluted in the third peak for a representative variant (F10, Fig. 3, C and D) declined monotonically with increasing concentration at SEC column injection (fig. S3G), whereas the percentage of protein eluted in the third peak for 602 IC plateaued at a concentration equivalent to 5 µM (fig. S1E). As an experimental control, we prepared an IC construct in LN35 format, denoted Control IC, containing IL-2 fused to an irrelevant antibody (the anti-fluorescein antibody 4-4-20 (*33*), which shares the mouse immunoglobulin G [IgG] 2a kappa isotype with 602). A smaller fraction of Control IC was eluted in peaks 1 and 2 compared to 602 IC and variant F10 IC, presumably due to the absence of oligomeric complexes consistent with the lack of cytokine/antibody interaction (Fig. 3E, fig. S3, H-J). These observations aligned with results from analytical ultracentrifugation, which showed the presence of small amounts of dimeric and trimeric species for 602 IC and F10 IC, but only monomeric species for Control IC (fig. S3L). All 602 IC variants and Control IC migrated at the expected molecular weights via SDS-PAGE (fig. S3K).

To characterize the biophysical behavior of engineered 602 IC variants, BLI was used to assess binding to immobilized IL-2, IL-2Rα, and IL2Rb. The 602 IC variants A8 IC, C5 IC, and F10 IC showed weaker interaction with immobilized IL-2 compared to the parent IC (Fig. 3F, fig. S4A, fig. S5A, table S4), indicative of more stable intramolecular interactions for the IC variants due to stronger cytokine/antibody affinity. Consistent with yeast surface competitive binding experiments (fig. S3H), all 5 IC variants, like the parent 602 IC, did not interact with immobilized IL-2Rα (Fig. 3G, fig. S4B, fig. S5B). All five 602 IC variants bound IL-2Rβ with similar affinity compared to the parent 602 IC and Control IC (Fig. 3H, fig. S4C, fig. S5C).

Variant F10 was further analyzed and showed higher affinity for IL-2 as an scFv compared to 602 scFv (fig. S6, A and C, table S5), largely due to a substantial reduction in dissociation rate. Consistent with this, thermostability measurements conducted by DSF revealed that the T_m_ for F10 IC was at least 1.4°C higher than that for 602 IC, and the T_m_ 602 IC was at least 0.6°C higher than that for IL-2/602 complex (fig. S3, M and N, table S2). Notably, IL-2/602 complex (at a stoichiometrically equivalent 2:1 cytokine:antibody ratio) bound to immobilized IL-2 with only slightly reduced affinity compared to the unbound 602 antibody (Fig. 3F, fig. S4A, fig. S5A), while IL-2/F10 complex and F10 IC showed minimal binding to immobilized IL-2, further demonstrating the increased stability of IL-2 binding for the F10 variant compared to the parent 602 antibody (fig. S6, B and D). Moreover, F10 IC exhibited weaker binding to immobilized IL- 2 compared to 602 IC, further highlighting its improved intramolecular stability. As expected, Control IC did not interact with immobilized IL-2 (Fig. 3F, fig. S2A, fig. S4A, fig. S5A). Kinetic data revealed that both 602 IC and the F10 IC variant blocked IL-2/IL-2Rα interaction more effectively than IL-2/602 complex (fig. S5B, table S1). Control IC bound IL-2Rα with increased affinity compared to IL-2 due to bivalency (Fig. 2B, Fig. 3G, table S1). Notably, kinetic analyses showed that 602 IC, F10 IC, and IL-2/602 complex had slower IL-2Rβ association rates compared to Control IC, most likely a consequence of steric effects resulting from intramolecular cytokine/antibody assembly. Nonetheless, the IL-2/IL-2Rβ interaction remained intact for all immunocytokines, allowing for ICs to orchestrate selective disruption of the IL-2/IL-2Rα interaction in a stable unimolecular format. Collectively, these binding and thermostability studies confirmed our evolution of 602 IC variants with enhanced stability of intramolecular cytokine/antibody assembly.

### Engineered 602 IC shows enhanced IL-2 bias toward effector cells

We hypothesized that preferential engagement of IL-2Rβ over IL-2Rα by our biased ICs would decrease the natural bias of the IL-2 cytokine toward IL-2Rα^+^ (T_reg_-like) cells, and thereby favor signaling on IL-2Rα^-^ (Eff-like) cells. Stimulation of a mixed population of IL-2Rα^-^ and IL-2Rα^+^ YT-1 cells revealed that 3 of the 5 engineered 602 IC variants showed stronger IL-2Rα^-^ cell bias compared to the parent 602 IC (fig. S4, D-F). Amongst the variants, F10 IC (which contained the mutations T101S, F225S, and G227D [T99S in CDR3H and F91S and G93D in CDR3L, Kabat numbering]) was most skewed toward IL-2Rα^-^ cell stimulation. In fact, F10 IC flipped the natural bias of IL-2 to favor signaling on IL-2Rα^-^ Eff-like cells over IL-2Rα^+^ T_reg_-like cells (Fig. 4, A-C, table S6), and this molecule was therefore selected for further characterization. In contrast, like IL-2, Control IC and IL-2/602 complex were biased toward IL-2Rα^+^ over IL-2Rα^-^ cells. Similar cell activation biases were manifested on freshly-isolated human peripheral blood mononuclear cells (PBMCs), and a significant advantage was observed for F10 IC compared to 602 IC (Fig. 4, D-H). Specifically, signaling of 602 IC and F10 IC was significantly more impaired on T_regs_ compared to conventional CD4^+^ T cells (CD4^+^ T_convs_) and CD8^+^ T cells, whereas Control IC and IL-2/602 complex showed similar bias to unconjugated IL-2 or further biased the cytokine towards T_reg_ cell activation. The activation ratios for both CD4^+^ T_convs_ and CD8^+^ T cells relative to T_regs_ were higher for F10 IC versus the parent 602 IC (Fig. 4, F and H), highlighting the benefit of our antibody engineering efforts in more acutely biasing IL-2 behavior, particularly in the context of a mixed immune cell environment. Taken together, our signaling data demonstrate that enhancing the IL-2Rα competitive activities and IL-2 affinity of 602 IC led to superior elimination of the competitive advantage endowed to T_regs_ by high basal expression of IL-2Rα, resulting in enhanced IL-2-mediated stimulation of immune effector cells.

**Fig. 4.**
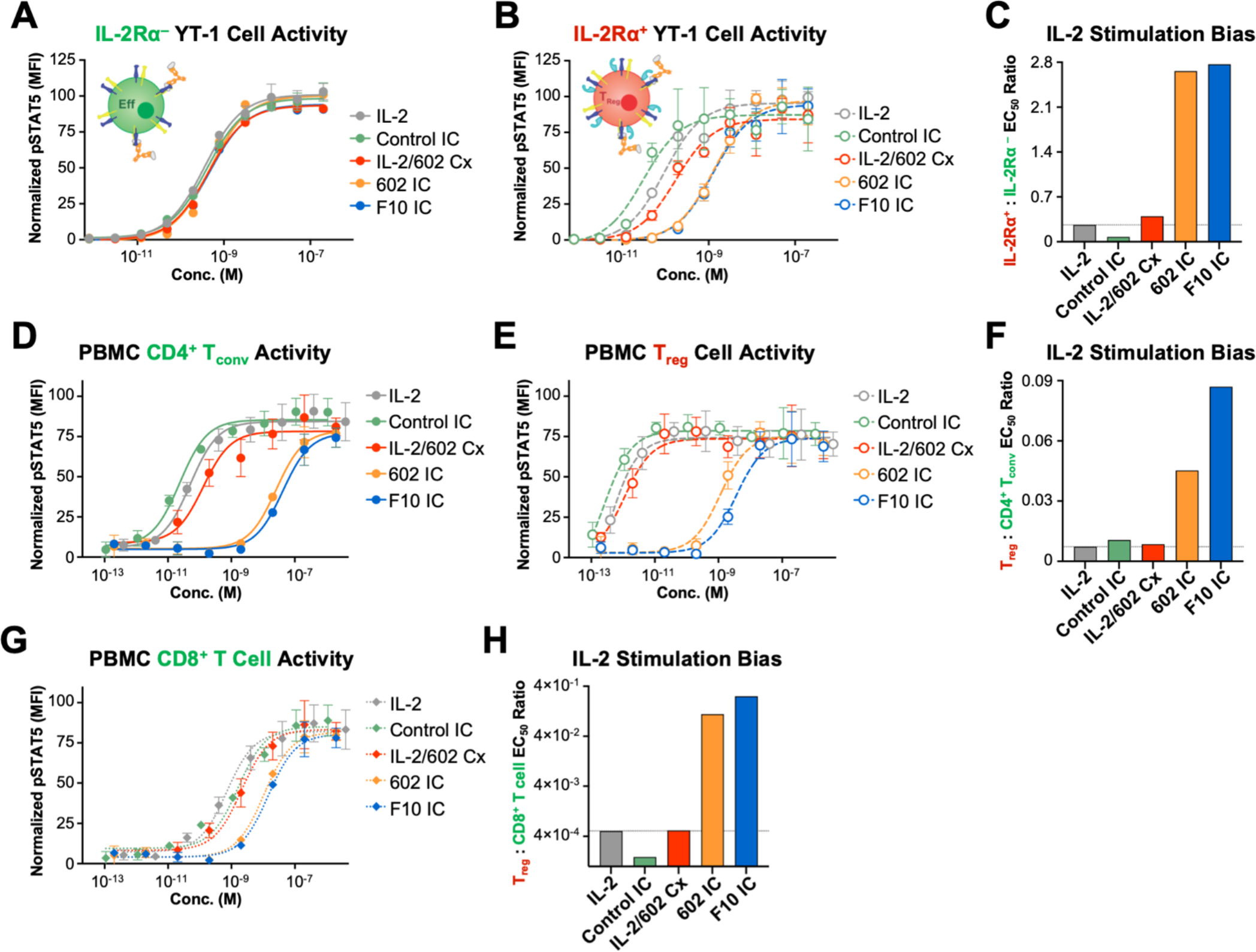
Enhanced IL-2Rα competition of the constituent antibody improves the immune effector cell bias of 602 IC. (A) STAT5 phosphorylation response of IL-2Rα^-^ YT-1 human NK cells treated with IL-2, IL-2/602 Cx (1:1 molar ratio), 602 IC, F10 IC, and Control IC. Data represent mean ± SD (n=3). **(B)** STAT5 phosphorylation response of IL-2Rα^+^ YT-1 human NK cells treated with IL-2, IL-2/602 Cx (1:1), 602 IC, F10 IC, and Control IC. Data represent mean ± SD (n=3). **(C)** Ratio of the STAT5 phosphorylation EC_50_ values for assays shown in (A) and (B). **(D)** STAT5 phosphorylation response of CD4^+^ T_convs_ (CD3^+^CD4^+^CD8^-^FoxP3^-^) cells within mixed human PBMCs treated with IL-2, IL-2/602 Cx (1:1 molar ratio), 602 IC, F10 IC, and Control IC. Data represent mean ± SD (n=3). **(E)** STAT5 phosphorylation response of T_reg_ (CD3^+^CD4^+^CD8^-^FoxP3^+^) cells within mixed human PBMCs treated with IL-2, IL-2/602 Cx (1:1 molar ratio), 602 IC, F10 IC, and Control IC. Data represent mean ± SD (n=3). **(F)** Ratio of the STAT5 phosphorylation EC_50_ values for CD4^+^ T_convs_ to T_regs_ from assays shown in (D) and (E). **(G)** STAT5 phosphorylation response of CD8^+^ T cells (CD3^+^CD8^+^CD4^-^) cells in mixed human PBMCs treated with IL-2, IL-2/602 Cx (1:1 molar ratio), 602 IC, F10 IC, and Control IC. Data represent mean ± SD (n=3). **(H)** Ratio of the STAT5 phosphorylation EC_50_ value for CD8^+^ T cells to T_regs_ from assays shown in (G) and (H).

### Structural basis for 602-mediated disruption of IL-2/IL-2Rα interaction

To further characterize the mechanistic behavior of the 602 antibody and engineered variants thereof, we determined the molecular structure of the complex between IL-2 and 602 in an scFv format. Crystals of the 602 complex diffracted to 1.65 Å and the structure was phased via molecular replacement (Fig. 5A, top). Due to the high resolution achieved for the IL-2/602 complex structure, we could confidently model the positions of individual side chains. Engagement of IL-2 by 602 occludes ∼740 Å^2^ of area on the cytokine surface. On the cytokine side, the binding interface is primarily comprised of residues in the AB loop (residues 38-45), the B helix (residues 62-69), and the CD loop and N-terminal end of the D helix (residues 107-116). On the 602 scFv side, the CDR3 domains of both the heavy and light chains contribute substantially to the interaction (Fig. 5B); notable features include hydrogen bonds by neighboring residues R100 and T101 and a stretch of interacting residues in the CDR3L region (S225, W226, D227) (Fig. 5C, top). CDR1L also contributed to the interface between 602 and the IL-2 cytokine, including salt bridges formed by residues D162 and R166. To elucidate the rationale for enhanced binding of IL-2 to F10 versus 602, we determined the 1.7 Å resolution crystal structure of the IL- 2/F10 scFv complex (Fig. 5A, bottom). Interestingly, while 602 and F10 occlude an almost identical interface on the cytokine, the two exhibit a subtle offset in binding angle (Fig. 5, B and C). In addition, the IL-2-binding paratope on the F10 antibody is nearly identical to that of the parent 602 antibody. Whereas the T101S mutation in CDR3H and the F225S mutation in CDR3L introduced for F10 did not appear to alter cytokine/antibody interactions, the mutation of glycine at position 227 in CDR3L of 602 to aspartate in F10 introduced a new salt bridge with residue R38 of IL-2, consistent with the observed improvement in IL-2 binding (Fig. 5C).

**Fig. 5.**
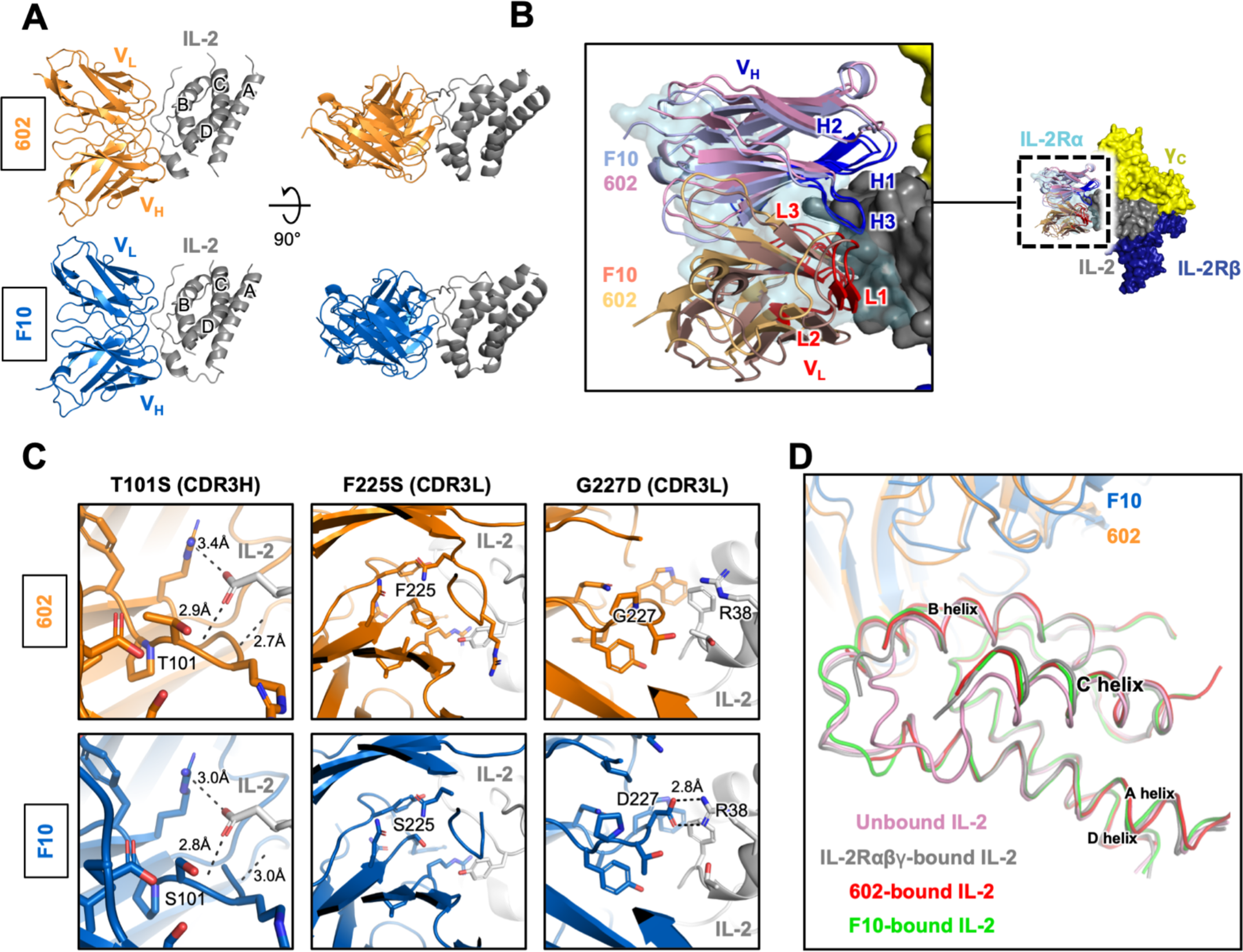
Structural basis for biased activity of F10 IC. (A) Structure of the IL-2/F10 scFv and IL-2/602 scFv complexes. **(B)** Overlay of the IL-2/F10 scFv complex structure with the structure of the quaternary IL-2 cytokine/receptor complex (PDB 2B5I). F10 scFv was found to occlude ∼735 Å^2^ of the IL-2 surface, primarily in the α-binding interface, but does not interfere with IL- 2Rβ or γ_C_ engagement. Heavy chain (HC) and light chain (LC) CDRs are delineated. **(C)** Detailed view of differential F10 scFv interactions with IL-2 compared to the parent 602 scFv. Mutations T101S, F225S, and G227D (T99S in CDR3H and F91S and G93D in CDR3L, Kabat numbering) are shown, depicting all side chains within 5 Å of the mutated residues (sticks) and inter-chain polar contacts within the same distance (black dashed lines). **(D)** Overlay of IL-2 in the unbound (pink, PDB 3INK), IL-2Rαβγ-bound (gray, PDB 2B5I), 602 scFv-bound (red), and F10 scFv- bound (green) states. Note that displacement of helix C is similar in the receptor-bound and scFv- bound states.

When comparing the structure of IL-2/antibody complexes to the structure of IL-2 bound to its high-affinity receptor complex (PDB 2B5I), we noted that 602 and F10 both had considerable overlap with the IL-2Rα subunit but did not overlap with the IL-2Rβ or γ_C_ subunits (Fig. 5B, fig. S7, A and B). This focused obstruction of IL-2/IL-2Rα engagement was reminiscent of that observed for other reported Eff-biasing antibodies, specifically the anti-mouse IL-2 antibody S4B6 (*22–26, 34, 35*) and the anti-human IL-2 antibody NARA1 (*27, 28*). The epitopes on IL-2 engaged by 602 and F10 are similar, although F10 occupies less interface on the γ_C_-adjacent lower portion of IL-2 compared to 602, when viewed from a top-down perspective. Despite this slight difference, the epitopes engaged by 602 and F10 show identical overlap with the epitope engaged by the IL- 2Rα subunit (fig. S7B). The predicted S4B6 binding interface on hIL-2 is distinct from the 602/F10 binding interface on hIL-2, with S4B6 binding closer to the IL-2Rβ subunit and 602/F10 binding closer to the γ_C_ subunit. Nonetheless, the overlap between the S4B6 and IL-2Rα interfaces and the 602/F10 and IL-2Rα interfaces are strikingly similar. In contrast, the NARA-1 interface is centered between γ_C_ and IL-2Rβ, and its overlap with the IL-2Rα subunit interface is more focused on the lower portion of IL-2, when viewed from a top-down perspective (fig. S7B).

Furthermore, we noted that 602- and F10-bound IL-2 were essentially superimposable with receptor-bound IL-2, recapitulating the 15° helical shift that the C helix of IL-2 undergoes upon receptor binding (Fig. 5D) (*16, 25, 36*). As was previously observed for the S4B6 antibody, the 602 and F10 antibodies allosterically prime IL-2 for engagement of the IL-2Rβ and γ_C_ subunits by enforcing this shift in the C helix, which is similarly induced by IL-2Rα binding. In addition, whereas structural alignments of mouse IL-2-bound S4B6 suggest that it may sterically clash with IL-2Rβ (*25*), 602 and the F10 variant do not show any steric clashes with IL-2Rβ due to the shift in binding topology away from the IL-2Rβ subunit relative to S4B6 (fig. S7C). Overall, the structural properties of the IL-2/602 complex rationalize the antibody’s focused inhibition of the IL-2Rα interaction, and comparison with the IL-2/F10 complex reveals the mechanism for enhanced cytokine/antibody affinity and improved stability of the F10 versus the 602 IC.

### ICs elicit similar gene expression profiles compared to IL-2 cytokine

To broadly examine whether antibody fusion of IL-2 in the IC format altered functional signaling compared to the unfused IL-2 cytokine, RNA-Seq analysis was performed on CD8^+^ T cells freshly isolated from human PBMCs from two healthy donors that were treated with either PBS or a saturating dose of IL-2 (1 µM), F10 IC (0.5 µM), or Control IC (0.5 µM) (fig. S8A). Overall, the gene expression profiles for the two ICs were very similar to that of IL-2 (Fig. 6, A and C). Some differences emerged after 24 hr of treatment, including differential expression of hallmark genes associated with STAT5 signaling, which showed higher induction in cells stimulated with F10 IC or Control IC compared to IL-2 (Fig. 6B). Though a saturating concentration of all proteins was used to stimulate the cells, the differences in STAT5 hallmark genes, as well as some genes involved in T cell activation and apoptosis (fig. S8, B and C), were likely due to differences in the potency of the stimulation over time. The ICs may be less easily internalized relative to IL-2, or the bivalency of the ICs may induce stronger or more durable responses. Despite these slight deviations, the gene expression response to stimulation by both ICs was remarkably similar to that seen with IL-2, indicating that fusion to an antibody does not substantially alter the quality of IL- 2 signaling.

**Fig. 6.**
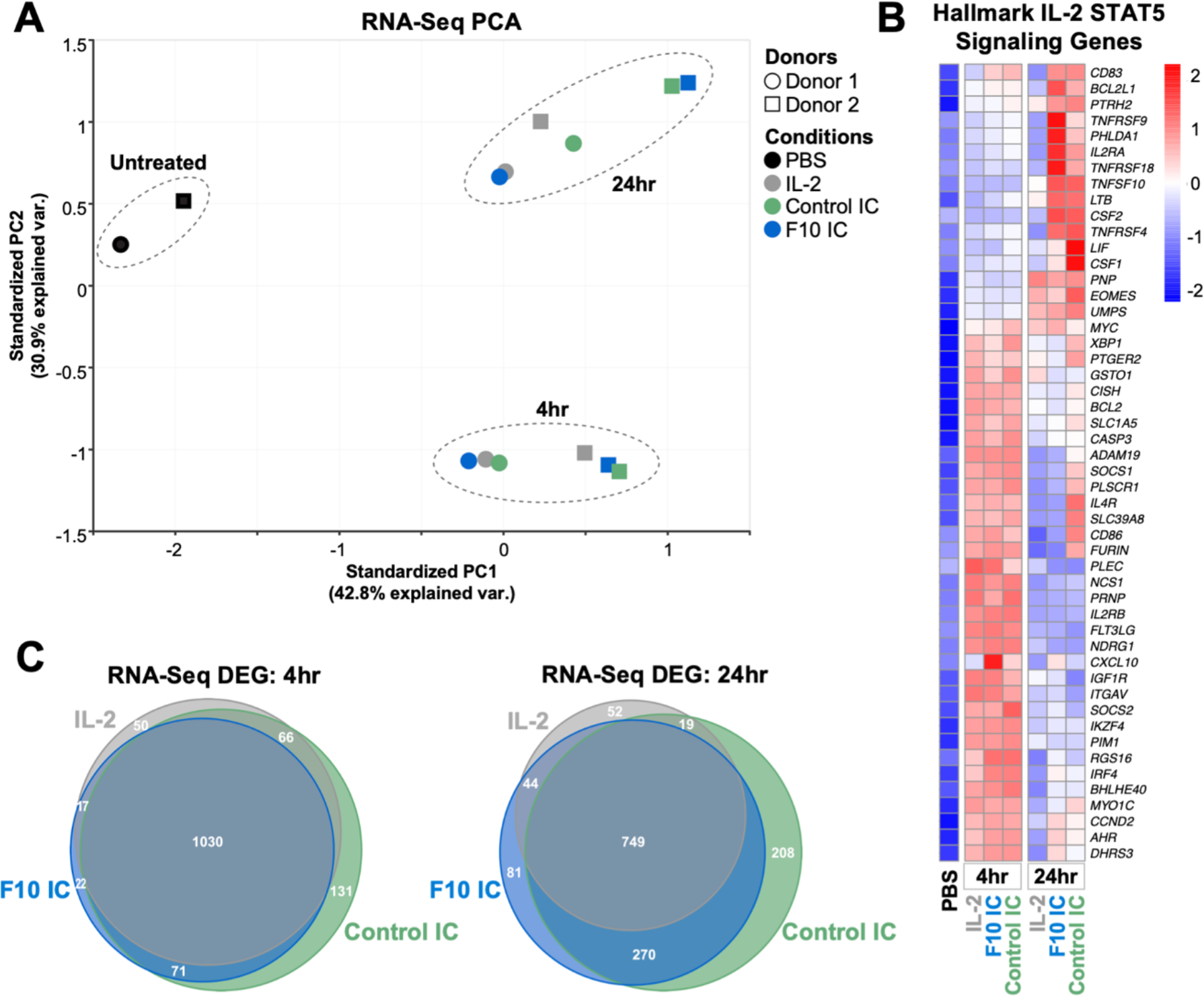
Unconjugated IL-2 and ICs induce similar gene expression profiles in human CD8^+^ T cells. (A) RNA-Seq analysis was performed on freshly isolated human CD8^+^ T cells from 2 independent donors stimulated with either PBS or a saturating concentration of IL-2 (1 µM), F10 IC (0.5 µM), or Control IC (0.5 µM) for either 4 hr or 24 hr. Principal component analysis (PCA) was performed on all evaluated genes. **(B)** Venn diagram depicting differentially expressed genes (DEG) at 4 (left) and 24 hr (right) for each treatment compared to untreated cells. Area of overlap corresponds to the number of shared DEG between treatment cohorts. **(C)** Heatmap representation of RNA-Seq analysis for hallmark STAT5 signaling genes. The color scale in the heatmap represents Z-score values ranging from blue (low expression) to red (high expression).

### Engineered 602 IC expands immune effector cell subsets in vivo

To determine whether our engineered IC’s enhanced tropism toward immune effector cell activation in vitro corresponded to increased expansion of Effs in live animals, we intraperitoneally administered Control IC, IL-2/602 complex (2:1 molar ratio), 602 IC, or F10 IC and evaluated the resulting abundance of various immune cell subsets in harvested spleens. IL-2/602 complex elicited the greatest overall expansion of the immune cell subsets compared to PBS treatment (fig. S9A). The 602 and F10 ICs elicited less expansion than IL-2/602 complex, but induced greater expansion of CD8^+^ T cells, NK cells, memory phenotype CD8^+^ T cells (CD8^+^ T_MPs_) and T_regs_ compared to PBS treatment. Only IL-2/602 complex and F10 IC expanded CD4^+^ T_convs_ above treatment with PBS, and treatment with Control IC suppressed expansion, with the greatest difference in expansion observed between Control IC and F10 IC. Relative expansion ratios for effector cell subsets versus T_regs_ were used to assess the overall pro-inflammatory immune biasing introduced by the different treatments (Fig. 7, A-C, fig. S9, C and D). F10 IC preferentially expanded CD4^+^ T_convs_, CD8^+^ T cells, CD4^+^ T effector memory cells (CD4^+^ T_EMs_), and CD8^+^ T_MPs_ relative to T_regs_, and effector bias was superior to that induced by both Control IC and IL-2/602 complex. F10 IC also preferentially expanded NK cells relative to T_regs_, and effector bias was superior to that induced by IL-2/602 complex and trended higher than that induced by Control IC. 602 IC elicited similar biased expansion of effector subsets compared to T_regs_, although the bias trended lower on all effector subsets.

**Fig. 7.**
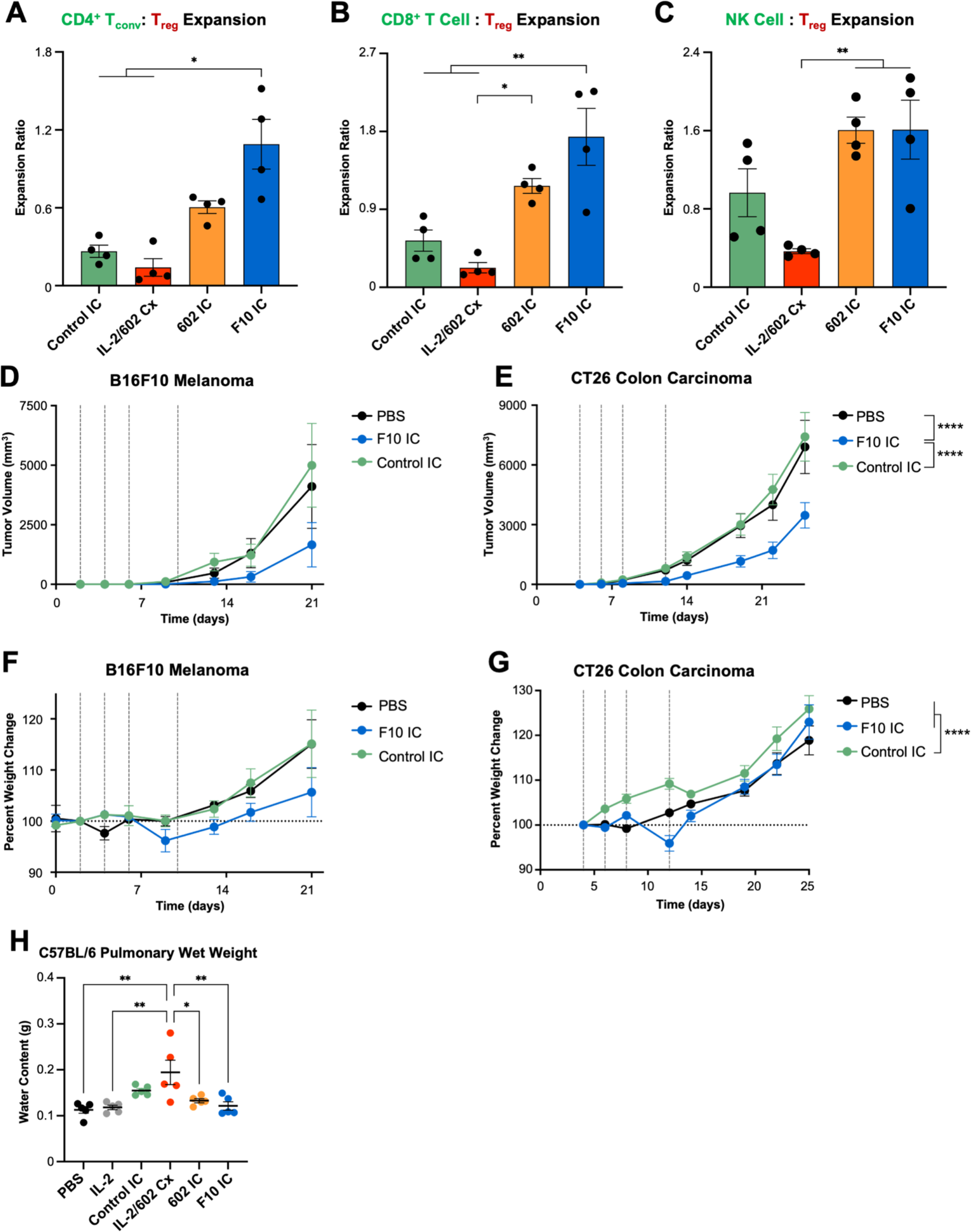
Engineered F10 IC promotes biased expansion of immune effector cells and improves **the therapeutic efficacy of IL-2.** C57BL/6 mice (n=4) were injected intraperitoneally daily for 4 days with the molar equivalent of 0.075 mg/kg IL-2/dose of Control IC, IL-2/602 complex (Cx) (2:1 cytokine:antibody molar ratio), 602 IC, or F10 IC. Mice were sacrificed on day 5, and spleens were harvested. The total cell counts of conventional T cells (CD4^+^ T_convs_, CD3^+^CD4^+^FoxP3^-^), CD8^+^ T cells (CD3^+^CD4^-^CD8^+^), natural killer (NK) cells (CD3^-^NK1.1^+^CD49b^+^), and T regulatory cells (T_regs_, CD3^+^CD4^+^CD25^+^Foxp3^+^) in each spleen were determined by flow cytometry. The ratios of CD4^+^ T_convs_ to T_regs_ **(A)**, CD8^+^ T cells to T_regs_ **(B)**, and NK cells to T_regs_ **(C)** were calculated. **(D, F)** C57BL/6 mice (n=7-9) were injected subcutaneously with 1×10^5^ B16F10 tumor cells and treated intraperitoneally on days 2, 4, 6, and 10 with either PBS or the molar equivalent of 0.125 mg/kg IL-2 of Control IC or F10 IC. Tumor size (D) and percent changes in body weight relative to weight at the time of tumor implantation (F) are shown. **(E, G)** BALB/c mice (n=8) were injected subcutaneously with 2×10^5^ CT26 tumor cells and treated intraperitoneally on days 4, 6, 8, and 12 with either PBS or the molar equivalent of 0.125 mg/kg IL-2 of Control IC or F10 IC. Tumor size (E) and percent changes in body weight relative to weight at the time of tumor implantation (G) are shown. **(H)** C57BL/6 mice (n=5) were injected daily for 4 days with PBS, 0.075 mg/kg IL-2, or the molar equivalent of 0.075 mg/kg IL-2 of Control IC, IL-2/602 complex (Cx) (2:1 cytokine:antibody molar ratio), 602 IC, or F10 IC. Mice were sacrificed on day 5, and pulmonary edema was assessed by measurement of lung water content. Data are shown as mean ± SEM. *P<0.05, **P<0.01, ***P<0.001, ****P<0.0001 by two-way ANOVA test.

Surprisingly, IL-2/602 complex showed less bias towards immune effector cells than the Control IC, due to significantly greater expansion of T_regs_ by IL-2/602 complex (Fig. 7, A-C, fig. S9, A, C and D). We speculated that this could be due to recruitment of effector function through the antibody Fc domain. As IL-2 remains on the surface of T_regs_ when interacting with only the IL- 2Rα subunit, and is only internalized upon binding to both IL-2Rβ and γ_C_ (*37*), T_regs_ are likely to have more Control IC on their surface compared to other immune cells, which express lower levels of IL-2Rα. This increased presence of Control IC on the surface could deplete T_regs_, thus attenuating their expansion relative to IL-2/602 complex treatment, in which case the antibody may freely dissociate. Consistent with this hypothesis, when Fc effector function was knocked out (referred to as ΔFc, LALA-PG Fc mutation (*38*)) in the Control IC, T_reg_ expansion was significantly greater (fig. S9, A and B). The only other significant difference observed between constructs containing native Fc versus ΔFc was in the Control IC-mediated expansion of CD8^+^ T_MPs_, which was also higher when effector function was knocked out. CD8^+^ T_MPs_ have substantially greater expression of IL-2Rβ that is not associated with higher levels of γ_C_ (*29*); thus, a lag in internalization on this cell subset may lead to Fc-mediated depletion. As no significant differences were observed for the knockout of effector function for F10 IC, ICs with the native Fc were used for therapeutic studies. Collectively, these results demonstrate that F10 IC preferentially expands effector immune cell subsets over T_regs_ in vivo, creating a potent immunostimulatory environment that could be leveraged to activate the immune system against diseases such as cancer.

### Engineered 602 IC inhibits tumor growth without inducing toxicity

To assess the therapeutic potential of our novel IL-2 biasing strategy, we tested the performance of F10 IC in multiple mouse syngeneic tumor models. We first compared the antitumor activity of our F10 IC to that of IL-2 and IL-2/602 complex in a B16F10 syngeneic mouse melanoma model. Mice were inoculated subcutaneously with B16F10 cells and treated intraperitoneally on days 4, 8, and 11 following tumor implantation. IL-2/602 complex significantly inhibited tumor growth relative to PBS and IL-2, and F10 IC elicited even stronger suppression (fig. S9E). We next compared the performance of F10 IC to Control IC, to better evaluate whether F10 IC’s antitumor activity was driven by its enhanced bias or by advantages imparted by the IC format. Mice were inoculated subcutaneously with B16F10 cells and treated intraperitoneally with PBS, F10 IC, or Control IC on days 2, 4, 6, and 10 following tumor implantation (Fig. 7D). F10 trended towards tumor growth inhibition relative to Control IC (P=0.089). We further compared the ICs in a more immunogenic mouse tumor model, specifically the CT26 syngeneic mouse model of colorectal carcinoma. Mice were inoculated subcutaneously with CT26 cells and treated intraperitoneally with PBS, F10 IC, or Control IC on days 4, 6, 8, and 12 following tumor implantation (Fig. 7E). F10 IC significantly suppressed tumor growth compared to both Control IC and PBS. In both tumor models, the body weights for mice treated with F10 IC and Control IC were not significantly lower than those for mice treated with PBS (Fig. 7, F and G), indicating minimal toxicity. We also evaluated more acute adverse effects of IC treatment in the form of pulmonary edema and liver injury. Mice were treated daily with PBS, Control IC, IL-2/602 complex, 602 IC, or F10 IC for 4 days, and sacrificed on the fifth day, at which point the pulmonary wet weight and serum concentrations of liver enzymes aspartate aminotransferase (AST) and alanine transaminase (ALT) were measured. Treatment with IL-2/602 complex led to increased fluid in the lungs of C57BL/6 but not BALB/c mice, and none of the ICs led to significant changes in pulmonary wet weight (Fig. 7H, fig. S9F). No significant differences in liver enzyme concentrations were observed between any of the treatments (fig. S9, G and H). Overall, the results of mouse tumor studies demonstrate that F10 IC drives antitumor activity without inducing severe systemic toxicities frequently associated with IL-2 immunotherapy. Furthermore, F10 IC was found to be more effective in suppressing tumor growth than the unbiased Control IC and IL-2/antibody complex.

## DISCUSSION

In this work, we engineered F10 IC, a cytokine/antibody fusion protein in which the component antibody intramolecularly binds IL-2 and blocks its interaction with IL-2Rα to promote expansion of pro-inflammatory immune cells. The therapeutic efficacy of IL-2 has long been hampered by the cytokine’s indiscriminate expansion of both pro- and anti-inflammatory cells as well as its short half-life. Through optimized genetic fusion of IL-2 to the antibody, F10 IC exhibits robust bias towards expansion of immune effector cells over T_regs_, boosting antitumor activity without inducing toxicity, while also dramatically improving cytokine half-life.

Numerous other IL-2-based molecules that bias activity towards effector cells by disrupting IL- 2/IL-2Rα interactions have been described (*21, 28, 29, 39–44*). Many of these molecules include mutations to the native human IL-2 sequence to eliminate IL-2Rα interactions (*21, 39–42, 44*), and most have additional mutations to enhance interactions with IL-2Rβ and γ_C_ (*21, 40–42, 44*). While some of these engineered cytokines have genetically fused the mutated IL-2 to other proteins (*40–42, 44*) or conjugated them to polyethylene glycol (PEG) (*45, 46*), others are still limited by short half-life. Moreover, direct mutations to IL-2 may reduce its stability and/or increase the chances of developing neutralizing anti-drug antibodies (*43*). Further, directly engineering IL-2 to interact more strongly with IL-2Rβ and γ_C_ can lead to systemic increases in IL-2 activity, which can result in inappropriate activation of T cells in non-target, non-tumor tissues. Despite these concerns, several of these molecules are actively being studied in clinical trials (*21, 28, 40, 44*), suggesting that each iterative improvement to biased IL-2 therapies brings them closer to therapeutic application. Alternative strategies involving native IL-2 and IL-2-targeted antibodies that block IL-2Rα interaction have also been developed, but most reports do not fuse IL-2 to the antibody (*22–24, 29*) and therefore allow cytokine dissociation, leading to pleiotropic activities and rapid clearance. In particular, while previous work examining the therapeutic potential of complexes of IL-2 and 602 (the F10 parent antibody) (*22–24*), our efforts demonstrated that the enhanced stability and bias resulting from cytokine/antibody fusion significantly improved tumor suppression compared to IL-2/antibody complex treatment. Recently, a molecule known as NARA1leukin grafted IL-2 into the binding domain of the NARA1 antibody (*27, 28*). This unimolecular formulation is being evaluated in active clinical trials (NCT04855929, NCT05578872), illustrating the promise for translation of cytokine/antibody fusion proteins. Our modular approach of terminal cytokine fusion allows for ready extension to other anti-IL-2 antibodies as well as other cytokine systems of interest.

Studying the impact of IC linker length on pSTAT5 signaling in Eff-like and T_reg_-like cells demonstrated that non-optimal fusion of IL-2 to the antibody dramatically diminished the potency and biasing of the IC. The reduced signaling activity observed in 602 IC LN15, and to a lesser extent in LN25, seemed to contradict the apparent functionality of all ICs based on binding studies. It is possible that shorter linker ICs attenuated signaling by disrupting the interaction between IL- 2 and γ_C_. The shorter linkers may have sterically or allosterically interfered with the γ_C_ binding site on the cytokine, which is immediately adjacent to the linker at the C-terminus of IL-2. Thus, these important engineering observations demonstrate that maximizing IC activity requires optimizing cytokine/antibody fusion format, and fusions that force unfavorable conformations can differentially ablate cytokine functions.

Linker optimization was also found to be critical for maximizing the yield of monomeric IC. During elution from protein G, the IC is highly concentrated, and low pH conditions destabilize non-covalent binding interactions. These conditions are prime for the exchange of IL-2 between proximal ICs, which leads to oligomerization. Indeed, 602 ICs with short linkers showed increased oligomerization, suggesting that these molecules did not allow for a stable intramolecular interaction between the cytokine and antibody within the IC. However, extending the linker appeared to eliminate intramolecular instability, favoring monomeric assembly of ICs.

Our 602 engineering campaign employed a competitive selection approach to further enhance antibody bias towards effector cell expansion. Structural data revealed that 602 sterically blocked IL-2/IL-2Rα binding without interfering with IL-2/IL-2Rβ or IL-2/γ_C_ interactions. Moreover, this antibody reproduced a conformational change in the C helix of IL-2 that the cytokine undergoes upon binding to IL-2Rα, which allosterically potentiates IL-2Rβ binding (*16, 25, 36*). F10 was found to recapitulate these structural properties of 602, but also introduced an additional salt bridge at the cytokine/antibody interface that led to higher affinity interaction. A slight positional shift was also observed between the 602 and F10 backbones. Collectively, the structural modifications induced by the F10 mutations promoted greater functional bias towards immune effector cell activation both in vitro and in vivo. Importantly, in contrast with previous efforts in generating IL- 2 muteins (*21, 41, 42*), the structural advantages for F10 IC were realized without modifying the IL-2 sequence, thus improving bias towards effector cells without increasing potential immunogenicity of an engineered cytokine.

Comparison to other reported anti-IL-2 antibodies that bias the cytokine towards expansion of effector cells revealed a unique binding topology for 602 and F10, despite their functional overlap with these other antibodies. Interestingly, despite the divergent binding orientations of the 602/F10 and S4B6 antibodies, the overlap between the 602/F10 and IL-2Rα interfaces on hIL-2 was remarkably similar to the overlap between the predicted S4B6 and resolved IL-2Rα interfaces on hIL-2. However, NARA1 bound IL-2 with a very different topology relative to 602/F10 and the overlap between the NARA1 and IL-2Rα interfaces on hIL-2 showed a divergent profile, with the occluded region focusing more on the lower half of the cytokine, when viewed from a top-down perspective. These structural observations suggest that distinct antibody-based IL-2Rα occlusion strategies can lead to convergent functional outcomes.

There were no observed differences between the intracellular programs induced in CD8^+^ T cells treated with IL-2 versus ICs based on RNA-Seq analysis. However, in vivo studies revealed differences in the potency of T_reg_ expansion between Control IC and IL-2/602 complex. While it is possible that the intracellular signaling pathways in T_regs_ are more sensitive than those of CD8^+^T cells to differences between stimulation with IL-2 versus ICs (*47, 48*), our comparison between native Fc and Fc with effector function knocked out suggest that differences in T_reg_ expansion following Control IC treatment can be at least partially explained by Fc-mediated depletion of cells that bind, but do not immediately internalize the IC. High expression of the non-internalizing IL- 2Rα subunit (*37*) on T_regs_ and high expression of IL-2Rβ on CD8^+^ T_MPs_ (*29*) could account for the impact of Fc knockout on the Control IC in these cell lines (Supplemental Figure 9, A and B). However, since Fc knockout did not fully restore Control IC-mediated expansion in T_regs_, additional factors such as receptor binding kinetics, internalization, and/or endosomal processing may also contribute (*48, 49*).

Elucidating the pharmacokinetic kinetic behavior and cellular processing dynamics of engineered immunocytokines will be essential in translating these molecules to the clinic, specifically for optimization of dosing amounts and schedules. As effector cells transiently express IL-2Rα once activated (*1, 2*), the dose scheduling of our IC must be carefully considered in order to best capitalize on this feedback in the context of immune responses associated with different types of cancer. Moreover, characterizing receptor expression dynamics will inform dosing regimens for combination treatments that incorporate other therapeutic modalities. The promising findings from this study in terms of the safety and efficacy of F10 IC in mouse models of cancer motivate further preclinical development of this molecule to advance clinical translation. We developed F10 IC by fusing human IL-2 to a mouse antibody; thus, both the variable and constant domains of our antibody will need to be humanized prior to any clinical testing to mitigate potential immunogenicity concerns. Nonetheless, the use of a mouse antibody in our syngeneic tumor models likely provided useful predictions for neutralization, toxicity, and other immune-mediated effects for what would be seen for a humanized antibody administered to human patients. We also note that our in vivo testing of F10 IC was limited to treatment of syngeneic mouse tumors. As with other hIL-2-based therapies, therapeutic efficacy relies upon the presence of tumor-specific Effs in the tumor microenvironment, as well as the ratio of Effs to T_regs_. Thus, characterization in additional settings, such as spontaneous disease models and humanized mouse models, will be required to further evaluate the therapeutic performance of F10 IC. Subsequent investigations in non-human primates will also be needed to validate the safety, pharmacokinetic behavior, biodistribution, and efficacy of our molecule prior to advancement to clinical trials.

Reports involving other IL-2 muteins, cytokine/antibody complexes, and fusion proteins that act by blocking the IL-2/IL-2Rα interaction and enhancing IL-2 interaction with IL-2Rβ and γ_C_ demonstrate that combining F10 IC with additional cancer therapeutics, including immune checkpoint inhibitor antibodies (*29, 50*), cancer vaccines (*28*), or adoptive cell transfer (*27*) will only further enhance tumor suppression. Given the numerous active clinical programs involving engineered IL-2 proteins (*10*), the stability, selectivity, safety, and antitumor efficacy of F10 IC make it appealing for clinical translation. Moreover, the modularity of its single-molecule construction readily invites adaptation to additional formats that will enable greater homing to and retention in the tumor microenvironment.

## MATERIALS AND METHODS

### Study Design

The objective of this study was to develop an IL-2/antibody fusion protein (IC) that preferentially activates Effs over Tregs, demonstrate that this IC blocks the IL-2/IL-2Rα interaction, enhances the IL-2/IL-2R interaction, biases IL-2 signaling in favor of Eff subsets in vitro, expands Eff subsets in vivo, and enhances antitumor efficacy in vivo without inducing toxicity. The single-agent IC was comprised of 2 IL-2 molecules genetically fused to the N-terminus of the light chains of an engineered anti-IL-2 antibody. This molecule was optimized via directed evolution, employing the yeast surface display platform, to establish the lead candidate, denoted F10 IC. The interactions between the IC and IL-2Rα as well as IL-2R were assessed by BLI. Additional biophysical and biochemical properties of F10 IC were examined by crystallographic structure analysis, analytical ultracentrifugation, and thermal unfolding experiments. In vitro signaling behavior of F10 IC compared with other IL-2 treatments was characterized through STAT5 phosphorylation studies in IL-2Rα^-^ (*30*) or IL-2Rα^+^ (*31*) YT-1 human NK cells, human PBMCs, and purified human CD8^+^ T cells. RNA expression in purified human CD8^+^ T cells following treatment with ICs versus IL- 2 was analyzed to compare overall differences in cell programs activated. Biased expansion of splenic effector immune cell populations in mice treated with F10 IC or other IL-2 interventions was assessed by flow cytometry. The therapeutic efficacy of F10 IC in suppressing tumor growth was subsequently studied in 2 mouse syngeneic tumor models (B16F10 and CT26). All mice were randomly allocated into experimental groups. Toxicity of the IC was evaluated through measurement of body weight loss, pulmonary edema, and liver injury. Investigators were not blinded to the treatment. Figure legends list the sample sizes, numbers of biological replicates, numbers of independent experiments, and statistical methods for each study.

### Cell lines

Human embryonic kidney (HEK) 293F cells were cultivated in Freestyle 293 Expression Medium (Thermo Life Technologies), supplemented with 10 U/mL penicillin-streptomycin (Gibco). Unmodified YT-1 (*30*) and IL-2Rα^+^ YT-1 human natural killer (NK) cells (*31*) were cultured in RPMI complete medium (RPMI 1640 medium supplemented with 10% fetal bovine serum, 2 mM L-glutamine, minimum non-essential amino acids, sodium pyruvate, 25 mM HEPES, and penicillin-streptomycin [all reagents from Gibco]). CT26 cells were maintained in RPMI-1640 (Sigma) with 10% FBS and 100 U/mL penicillin-streptomycin (Gibco). B16F10 cells were maintained in and DMEM high glucose medium (Sigma) supplemented with 10% FBS and 100 U/mL penicillin-streptomycin (Gibco). All cell lines were maintained at 37°C in a humidified atmosphere with 5% CO_2_. HEK 293F cells were maintained on a shaker set to 125 rpm.

### Protein expression and purification

The variable heavy (V_H_) and variable light (V_L_) chain sequences of the 602 antibody were determined by PCR amplification from 602 antibody-expressing hybridoma cells. cDNA encoding the V_H_ and V_L_ of MAB602 was obtained by performing reverse-transcription-polymerase chain reaction (RT-PCR) on mRNA isolated from MAB602 hybridoma cells. V_H_ cDNA was digested with the HindIII and KpnI restriction enzymes and ligated into the pSecTaqA vector; V_L_ cDNA was digested with the BamHI and XhoI restriction enzymes and ligated into the pcDNA 3.1^Hygro^ vector. Sequencing of the V_H_ and V_L_ was performed using the Sanger method. Recombinant antibodies were formulated as mouse immunoglobulin (IgG) 2a kappa isotype to match the parent clone. The heavy chain (HC) and light chain (LC) of the 602 antibody were separately cloned into the gWiz vector (Genlantis) (*51*). For wild type 602 immunocytokines (ICs) and variants thereof, the human IL-2 cytokine (amino acids 1–133) was fused to the full-length 602 antibody at the N-terminus of the LC by a flexible (G_4_S)_N_ linker (N=2, 3, 5, or 7; Figure 1). The control IC was comprised of hIL-2 fused to the N-terminus of the LC of an irrelevant anti-fluorescein antibody, 4-4-20 (*33*), with identical isotype to that of 602. For effector function knockout (ΔFc) ICs, mutations L234A, L235A, and P329G (*38*) were introduced to the heavy chain of the antibody.

Antibodies and ICs were expressed recombinantly in human embryonic kidney (HEK) 293F cells via transient co-transfection of plasmids encoding the HC and either LC or an IL-2/LC fusion protein. HC and LC plasmids were titrated in small-scale co-transfection tests to determine optimal DNA content ratios for large-scale expression. HEK 293F cells were grown to 1.2×10^6^ cells/mL and diluted to 1.0×10^6^ cells/mL on the day of transfection. Plasmid DNA and polyethyleneimine (PEI, Polysciences) were independently diluted to 0.05 and 0.1 mg/mL, respectively, in OptiPro medium (Thermo Life Technologies) and incubated at room temperature for 15 min. Equal volumes of diluted DNA and PEI were mixed and incubated at room temperature for an additional 15 min. Subsequently, the DNA/PEI mixture (40 mL per liter cells) was added to a flask containing the diluted cells, which was then incubated at 37°C with shaking for 5 days. Secreted protein was harvested from HEK 293F cell supernatants by Protein G affinity chromatography, followed by size-exclusion chromatography on an ÄKTA^TM^ fast protein liquid chromatography (FPLC) instrument using a Superdex 200 column (Cytiva). Analysis of the relative distribution of IC eluted in each peak was performed using UNICORN analysis software v7.1 (Cytiva). Statistical analysis was conducted using GraphPad Prism data analysis software v9.0.

Human IL-2 (amino acids 1–133), IL-2Rα ectodomain (amino acids 1–217), IL-2Rβ ectodomain (amino acids 1–214), 602 scFv (VH-[G_4_S]_3_-VL), and F10 scFv (VH-[G_4_S]_3_-VL) each contained a C-terminal hexahistidine tag, and were produced from HEK 293F cells, as described for the 602 antibody and ICs. Proteins were purified via Ni-NTA (Expedeon) affinity chromatography followed by size-exclusion chromatography on a Superdex 200 column (Cytiva) using an FPLC instrument. All proteins were stored in HEPES-buffered saline (HBS, 150 mM NaCl in 10 mM HEPES pH 7.3). Purity was verified by SDS-PAGE analysis.

For preparation of biotinylated IL-2, IL-2Rα, and IL-2Rβ, a C-terminal biotin acceptor peptide (BAP) GLNDIFEAQKIEWHE sequence was included. Following Ni-NTA affinity chromatography, the cytokine and receptors were biotinylated with the soluble BirA ligase enzyme in 0.5 mM Bicine pH 8.3, 100 mM ATP, 100 mM magnesium acetate, and 500 mM biotin (Sigma). After overnight incubation at 4°C, excess biotin was removed by size-exclusion chromatography on an ÄKTA^TM^ FPLC instrument using a Superdex 200 column (Cytiva).

### Yeast surface binding studies

General yeast display methodologies were carried out with EBY-100 yeast cells, as described previously (*32, 52*). The V_H_ chain followed by the V_L_ chain of the 602 antibody, separated by a (G_4_S)_3_ linker (scFv format), were cloned into the yeast display vector pCT3CBN (containing the yeast agglutinin protein Aga2 N-terminal to the scFv and a C-terminal c-Myc detection tag). Yeast displaying scFvs were plated at 2×10^5^ per well in a 96-well plate and incubated in PBE buffer (phosphate-buffered saline [PBS] pH 7.2 with 0.1% bovine serum albumin [BSA] and 1 mM ethylenediaminetetraacetic acid [EDTA]) containing biotinylated IL-2 in the presence or absence of IL-2Rα for 2 hr at room temperature. Cells were then washed and stained with a 1:200 dilution of Alexa Fluor 647-labeled streptavidin (Thermo Fisher Scientific) in PBSA for 15 minutes at 4°C. After a final wash, cells were analyzed for target binding using a CytoFLEX flow cytometer (Beckman Coulter). Background-subtracted and normalized binding curves were fitted to a first-order binding model and equilibrium dissociation constant (K_D_) values were determined using GraphPad Prism software v9.0. Experiments were conducted at least twice with consistent results.

#### Generation of a mutagenic yeast-displayed library of 602 scFv variants

A targeted error-prone library that mutagenized the CDR1 and CDR3 of both the heavy and light chain variable domains was generated to preserve existing IL-2 interactions while also allowing for potentially beneficial, conservative alterations in binding. The 4 targeted CDRs were amplified by error-prone PCR using Taq polymerase (New England Biolabs), 1× Taq buffer (New England Biolabs), 2 mM manganese(II) chloride, 7 mM magnesium chloride, 0.2 mM of dATP and dGTP, 1mM dCTP and dTTP, 0.5 µM of each primer, and 0.2 ng/µL of the template. Following 5 amplification cycles, the PCR mix was transferred and diluted 1:5 into fresh mix lacking the template. The amplification, transfer, and dilution steps were repeated twice more, with the final transfer undergoing 20 total amplification cycles. The 5 framework sequences adjacent to the targeted CDRs were amplified using Phusion High-Fidelity DNA polymerase (Thermo Scientific). Framework fragments were assembled with the neighboring mutagenized CDR fragments by sequential, pairwise overlap extension PCR using Phusion polymerase. The final assembled fragments contained the full 602 scFv as well as homologous sequences (≥ 97nt) on both ends, which provided overlap with the cut yeast display vector, pCT3CBN.

EBY100 yeast were transformed with the cut pCT3CBN vector and mutagenized fragment (approximately 10 µg and 50 µg, respectively), which were assembled by homologous recombination, as previously described (*32, 52*). The library yielded 1.2×10^8^ transformants, which were grown in SDCAA media (pH 4.5) for 48 hr prior to passaging, and were then induced 24 hr later in SGCAA media (pH 4.5) at an OD of 1. Individual yeast plasmids were extracted via miniprep (Zymo Research) to verify proper insertion of the fragment into the vector backbone. A productive error rate—i.e., DNA mutations that resulted in changes to the amino acid sequence— of approximately 6% was observed.

### Mutagenic 602 scFv Library Selections

Yeast selection methodologies were modified from previously described protocols (*32, 52*). Each round of selection was performed with sufficient yeast to ensure 10-fold coverage of the library diversity. Yeast clones selected from each round were grown overnight at 30°C in SDCAA media for 2 days, followed by induction in SGCAA media for 2 days at 20°C.

In the first round of selection, the naïve mutagenic 602 scFv library was debulked via magnetic-activated cell sorting (MACS). A pre-clearing step was performed to eliminate variants that bound to Alexa Fluor 647-conjugated streptavidin (SA-AF647) (Thermo Fisher Scientific), and the library was then positively selected for variants that bound biotinylated IL-2 tetramerized with SA- AF647. All staining was performed in PBE solution. In the first MACS step, the yeast were incubated with 20 µg/mL SA-AF647 for 1 hr at 4°C, washed, and then incubated with 1:20 anti- Cy5/anti-Alexa Fluor 647 microbeads (Miltenyi Biotec) for 20 minutes at 4°C, washed, and then applied to an LS MACS separation column (Miltenyi Biotec), according to the manufacturer’s protocol. Yeast that flowed through the column (i.e., did not bind to SA-AF647 alone) were collected and prepared for the second MACS step. Biotinylated IL-2 was mixed with SA-AF647 in a 4:1 molar ratio, diluted in PBE, and incubated for 15 minutes to form tetramers. Yeast were incubated with 2 nM IL-2 tetramers for 2 hr at 4°C, then washed and incubated with anti-Cy5/anti-Alexa Fluor 647 microbeads as before. Yeast were applied to an LS MACS separation column, and yeast clones eluted from the column were collected, grown, and induced for the next round of selection.

A second round of selection isolated full-length 602 scFv variants (based on c-Myc tag expression) via MACS. Yeast were incubated with a 1:100 dilution of AF647-conjugated anti-c-Myc antibody (clone 9B11, Cell Signaling Technologies) in PBE for 2 hr at 4°C, incubated with anti-Cy5/anti-Alexa Fluor 647 microbeads, and applied to an LS MACS separation column. Eluted cells were collected, regrown, and induced.

The final three rounds of selection were performed by fluorescence-activated cell sorting (FACS) on a FACSymphony S6 cell sorter (Becton Dickinson), using decreasing amounts of biotinylated IL-2 in the presence of a large excess of nonbiotinylated IL-2Rα to select for the variants that competitively blocked IL-2 interaction with the IL-2Rα subunit. The first of these competitive FACS selections used 50 nM IL-2 with 1.5 µM IL-2Rα, and the second and third both used 30 nM IL-2 and 0.3 µM IL-2Rα. The third sort selected for variants with low off-rates by first incubating the library with IL-2, washing, and then incubating with IL-2Rα at room temperature for 2 hr to allow for IL-2 dissociation. In all FACS selections, variants in the top 5% with respect to IL-2 binding were collected.

### Bio-layer interferometry binding studies

Biotinylated human IL-2, IL-2Rα and IL-2Rβ were immobilized to streptavidin-coated tips for analysis on an Octet Red96 BLI instrument (Sartorius). Less than 5 signal units (nm) of IL-2 or each receptor subunit was immobilized to minimize mass transfer effects. PBSA (PBS pH 7.2 containing 0.1% BSA) was used for all dilutions and as dissociation buffer. Tips were exposed to serial dilutions of IL-2, 602 antibody, IL-2/602 complexes (formed by incubating a 1:1 molar ratio of IL-2 and 602 antibody for 30 minutes at room temperature), various ICs, or an irrelevant protein (the human monoclonal antibody trastuzumab) in a 96-well plate for 300 s. Dissociation was then measured for at least 240 s. Surface regeneration for all interactions was conducted using 15 s exposure to 0.1 M glycine pH 3.0. Analysis and kinetic curve fitting (assuming a 1:1 binding model) were conducted using Octet Data Analysis HT software version 7.1 (Sartorius). Normalized equilibrium binding curves were obtained by plotting the response value at the end of the association phase for each sample dilution and normalizing to the maximum value. Equilibrium binding curves were fitted and K_D_ values determined using GraphPad Prism data analysis software v9.0, assuming all binding interactions to be first order. Experiments were performed at least twice with similar results.

### Analytical Ultracentrifugation

Sedimentation analysis was performed using a ProteomeLab XL-I analytical ultracentrifuge (Beckman Coulter) (*53*). Protein samples were diluted in a buffer containing 10 mM HEPES, 150 mM NaCl, pH 7.5 before the analysis; the same buffer was used as a reference. IL-2/602 complex was prepared in the same buffer by incubating IL-2 and 602 Ab at a 2:1 molar ratio for 15 min at room temperature. The sedimentation velocity experiment was conducted at 36,000-48,000 rpm and 20°C using double sector cells and an An50-Ti rotor at 0.1–0.6 mg/ml protein loading concentration. In total, 100-300 absorbance scans were recorded at 230 or 280 nm at 2-6 min intervals. Buffer density, protein partial specific volumes, particle dimensions, and the values of s_20,w_ sedimentation coefficients corrected to standard conditions (corrected to 20°C and the density of water) were estimated in Sednterp (*54*). Data were analyzed with Sedfit (*55*) using the c(s) continuous sedimentation coefficient distribution model, and the figures were prepared in GUSSI (*56*).

### Crystallographic characterization of IL-2 in complex with 602 or F10 scFv

Complexation of IL-2 with the F10 or 602 scFv was carried out overnight in the presence of carboxypeptidases A and B at 4°C. The complex was re-purified using SEC and concentrated using Amicon 30 kDa MWCO products. Screening for crystallization of the F10 scFv/IL-2 complex was carried out using commercial sparse matrices. Narrowed screens for the F10 complexes were made based on Nextal PEGs Suite condition E3, and crystals were grown using the hanging drop method in a range of conditions containing between 15-30% PEG3350 w/v and between 100-350 mM ammonium fluoride. Crystals presented a variety of plate morphologies. Data collection was carried out at Stanford Synchrotron Radiation Lightsource (SSRL) beamline BL12-1. Phasing was carried out using Phaser MR for molecular replacement with search models derived from PDB 2B5I and PDB 7RK2, both modified in Sculptor. Processing was carried out using autoproc, PHENIX, and coot. Alignments and visual representations were carried out using PyMol. Interface characterizations were carried out using PDBePISACrystal screening for the 602 scFv/IL-2 complex was achieved in the same PEG3350 and ammonium fluoride conditions as F10, and data collection/processing was carried out similarly.

### Thermostability comparisons by differential scanning fluorometry

Thermal unfolding of IL-2, 602 Ab, IL-2/602 complex (molar ratio of 2:1 of IL-2 to 602 antibody), 602 IC and F10 IC was measured as change in intrinsic tryptophan fluorescence as a function of temperature via nano differential scanning fluorometry (nanoDSF). The samples (10 µL at 0.2 mg/mL in HBS) were loaded into standard treated NT.115 capillaries (NanoTemper), and absorption at 330 and 350 nm was measured on a Prometheus NT.48 instrument (NanoTemper) in the temperature range 30-95°C with a temperature slope of 1.5°C/min. The melting curves show the ratio of the absorbances at 330 nm over 350 nm versus temperature, and the melting temperatures (T_m_) were calculated in PR.ThermControl (ver. 2.1.1) software from the first derivative of the melting curves.

### YT-1 cell STAT5 phosphorylation studies

Approximately 2×10^5^ IL-2Rα^-^ (*30*) or IL-2Rα^+^ (*31*) YT-1 human NK cells were plated in each well of a 96-well plate. Cells were washed and resuspended in RPMI complete medium containing serial dilutions of IL-2, IL-2/602 complexes (formed by pre-incubating a 1:1 molar ratio of IL-2 and 602 antibody for 30 minutes at room temperature), or various ICs. Cells were incubated with treatments for 20 min at 37°C and immediately fixed by addition of paraformaldehyde (Electron Microscopy Sciences) to 3.7%, followed by 10 min incubation at room temperature. Cells were permeabilized by resuspension in ice-cold 100% methanol and overnight incubation at -80°C. Fixed and permeabilized cells were washed twice with PBSA buffer and incubated for 2 hr at room temperature with Alexa Fluor 647-conjugated anti-STAT5 pY694 antibody (BD Biosciences, clone 47/Stat5) diluted 1:50 in PBSA buffer. Cells were then washed with PBSA buffer and analyzed on a CytoFLEX flow cytometer (Beckman Coulter). Normalized mean fluorescence intensity (MFI) was calculated by subtracting the MFI of unstimulated cells and normalizing to the maximum signal intensity. Dose-response curves were fitted to a logistic model and half-maximal effective concentrations (EC_50_) were calculated using GraphPad Prism data analysis software v9.0. Experiments were conducted in triplicate and performed 3 times with similar results.

To compare the signaling for the selected 602 IC LN35 variants derived from library selections, mixed IL-2Rα^-^/IL-2Rα^+^ YT-1 cell assays were performed in which IL-2Rα^-^ YT-1 cells were lentivirally transduced such that 4% of cells were IL-2Rα^+^. For lentiviral transduction, full-length human IL-2Rα was cloned into the pCDH-CMV-MCS-EF1-Puro expression vector, and the pPACKH1 HIV Lentivector Packaging Kit (System Biosciences) was used to produce lentivirus particles, per manufacturer instructions. Approximately 3.5×10^6^ YT-1 cells were transduced by centrifugal inoculation (30 min at 800×g) in RPMI complete medium containing polybrene (8 µg/mL) and 15% harvested lentivirus-containing medium. For STAT5 phosphorylation studies, at least 420 IL-2Rα^+^ events per sample were collected via flow cytometry. Experiments were conducted in triplicate and performed at least three times with similar results.

### Human PBMC isolation and STAT5 phosphorylation studies

Leukopaks containing de-identified human whole blood were obtained from Anne Arundel Medical Blood Donor Center (Anne Arundel, Maryland, USA). PBMCs were isolated via Ficoll-Paque Plus (GE Healthcare) gradient, per manufacturer recommendations, followed by incubation with ACK lysis buffer (Gibco) for removal of red blood cells.

Approximately 2×10^6^ PBMCs were plated in each well of a 96-well plate immediately following isolation. Greater than 99% viability was confirmed by staining with LIVE/DEAD Fixable Aqua dead cell stain (Invitrogen) diluted 1:1000 in PBS for 15 minutes at 4°C. Cells were resuspended in RPMI complete medium containing serial dilutions of IL-2, IL-2/602 complexes (formed by pre-incubating a 1:1 molar ratio of IL-2 and 602 antibody for 30 minutes at room temperature), or various ICs. Cells were incubated with treatments for 20 min at 37°C, and immediately fixed and permeabilized with Transcription Factor Phospho Buffer kit (BD Pharmingen), per manufacturer recommendations. Cells were then stained for 2 hr at room temperature with the following antibodies at the manufacturer-recommended dilutions in 1× TFP Perm/Wash buffer: APC- eFluor780 anti-human CD3 (clone UCHT1, Invitrogen); PerCP-Cy5.5 anti-human CD4 (clone SK3, BD Pharmingen); BV605 anti-human CD8 (clone SK1, Biolegend); BV421 anti-human CD25 (clone M-A251, BD Horizon); Alexa Fluor 488 anti-human CD127 (clone eBioRDR5, Invitrogen); PE anti-human FoxP3 (clone 236A/E7, BD Pharmingen); and Alexa Fluor 647 anti- STAT5 pY694 (clone 47/Stat5, BD Biosciences). Cells were washed, resuspended in PBSA buffer, and analyzed on a CytoFLEX flow cytometer (Beckman Coulter). At least 1500 T_reg_ cell events (CD3^+^CD4^+^IL-2Rα^+^Foxp3^+^) were collected in each sample.

Data were analyzed using FlowJo software v10.6.1 (Tree Star). Normalized pSTAT5 MFI for T_regs_ (CD3^+^CD4^+^Foxp3^+^), CD4^+^ T_convs_ (CD3^+^CD4^+^Foxp3^-^), and CD8^+^ T cells (CD3^+^CD8^+^) were calculated by subtracting the MFI of unstimulated cells and normalizing to the maximum signal intensity. An example of the gating strategy used for these experiments is depicted in Supplemental Figure 10. Dose-response curves were fitted to a logistic model, and half-maximal effective concentrations (EC_50_) were calculated using GraphPad Prism data analysis software v9.0. Experiments were conducted in triplicate and performed at least twice using independent donors with similar results.

### RNA sequencing analysis of IL-2 and IC stimulated primary human CD8^+^ T cells

Human CD8^+^ T cells were purified from buffy coats from healthy donors at the NIH Blood Bank using EasySep^TM^ Human CD8^+^ T Cell Isolation Kit (StemCell Technologies). Approximately 2×10^4^ CD8^+^ T cells were plated in each well of a 96-well plate immediately following isolation. Cells were incubated with treatments for 20 min at 37°C, immediately fixed and stained for phospho-STAT5 as described for the YT-1 cells, and analyzed on a FACSCanto II flow cytometer (BD). Normalized mean fluorescence intensity (MFI) was calculated by subtracting the MFI of unstimulated cells and normalizing to the maximum signal intensity, and the mean average for the two donors was plotted. Dose-response curves were fitted to a logistic model, and half-maximal effective concentrations (EC_50_) were calculated using GraphPad Prism data analysis software v9.0.

For RNA-Seq experiments, 5×10^5^ purified human CD8^+^ T cells were cultured in complete RPMI 1640 medium and stimulated for 4 or 24 hr with 1 µM IL-2 or 0.5 µM (equivalent stoichiometric quantity) Control IC or F10 IC. Total RNA was isolated using Direct-zol RNA Microprep Kit (R2061, ZYMO Research), and RNA-Seq libraries were prepared using 120 ng of total RNA and Kapa mRNA HyperPrep Kit (Kapa Biosystems). Each library was indexed using NEXTflex DNA Barcodes-24 (BIOO Scientific) and amplified by PCR. Amplified libraries between 250-400 bp were recovered using 2% E-Gel pre-cast gels (ThermoFisher), purified with Zymoclean Gel DNA Recovery Kit (Zymo Research), and quantified by Qubit 3 Fluorometer (ThermoFisher). Equal amounts of the indexed libraries were then pooled and sequenced with an Illumina Novaseq platform at the NHLBI DNA Sequencing Core.

### RNA sequencing data analysis

Sequenced reads (50 bp, single end) were obtained with the Illumina CASAVA pipeline and mapped to the human genome hg38 (GRCh38, Dec. 2013) using Bowtie2 and Tophat2. Only uniquely mapped reads were retained. Raw counts that fell on exons of each gene were calculated and normalized using RPKM (Reads Per Kilobase per Million mapped reads). R Bioconductor package “edgeR” was used to identify differentially expressed genes between cells treated with IL-2 and the ICs at the 4 hr and 24 hr time points. Gene expression heat maps were generated with the R package “pheatmap”. Gene Set Enrichment Analysis (GSEA) was used to determine statistically significant overlaps between given pre-ranked genes and Molecular Signature Database (MSigDB).

### Mice

Female 8- to 12-week-old C57BL/6 and BALB/c mice were obtained from the colony kept at the Czech Centre for Phenogenomics (CCP), Prague, Czech Republic.

### In vivo immune cell subset expansion

For relative effector (CD4^+^ T_conv_, CD8^+^ T, NK) cell:T_reg_ proliferation studies, C57BL/6 mice were injected intraperitoneally (i.p.) with the molar equivalent of 0.075 mg/kg IL-2/dose of IL-2/602 complex (prepared by incubating IL-2 and 602 Ab at a 2:1 cytokine:antibody molar ratio in PBS for 15 min at room temperature), or molar equivalent of 0.075 mg/kg IL-2 /dose of 602 IC, F10 IC or Control IC (approx. 4 µg IC) in 250 µL of PBS, on days 1, 2, 3, and 4. Mice were randomly distributed into experimental groups, with approximately equal average body weight across groups. Mice were sacrificed on day 5 (24 hr after the last injection) by cervical dislocation, and spleens were harvested. Single-cell suspensions were prepared by homogenization (GentleMACS Dissociator, Miltenyi Biotec) and subjected to ACK red blood cell lysis buffer (Thermo Fisher Scientific). The isolated splenocytes were resuspended in modified PBE buffer (PBS with 2.5% fetal calf serum [FCS], 2.5 mmol EDTA) and blocked with 10% C57BL/6 mouse serum for 30 min on ice. The cells from each spleen were then split into 3 groups, which were each stained in modified PBE buffer for 30 min on ice with each of the following panels: CD4 T cell/T_reg_ panel, which included eFluor 450 anti-mouse CD3 (clone 17A2, ThermoFisher Scientific, 1:40), PerCP anti-mouse CD4 (clone RM4-5, BD Biosciences, 1:100), and APC anti-mouse CD25 (clone PC61.5, ThermoFisher Scientific, 1:300); CD8 T cell panel, which included PE-Cy7 anti-mouse CD3 (clone 145-2C11, ThermoFisher Scientific, 1:40); PerCP anti-mouse CD4 (clone RM4-5, BD Biosciences, 1:100), BD Horizon V500 anti-mouse CD8 (clone 53-7.62, BD Biosciences, 1:80), APC anti-mouse CD44 (clone IM7, ThermoFisher Scientific, 1:500), eFluor450 anti-mouse CD62L (clone MEL-14, ThermoFisher Scientific, 1:40); and NK cell panel, which included BD Horizon V500 anti-mouse CD3 (clone 500A2, BD Biosciences, 1:30), PE anti-mouse CD49b (clone DX5, ThermoFisher Scientific, 1:50), and APC anti-mouse NK1.1 (clone PK136, ThermoFisher Scientific, 1:125). Cells were then washed twice and fixed in Fixation/Permeabilization Buffer (BD Biosciences) for 1 hr on ice. The T_reg_ panel cells were stained with PE anti-mouse/rat Foxp3 (clone FJK-16s, ThermoFisher Scientific, 1:100) for 30 min on ice, followed by two final washes and resuspension in modified PBE buffer. Samples were analyzed on a BD Fortessa flow cytometer (BD Biosciences). The percentages of splenic lymphocytes that were conventional CD4^+^ T cells (CD4^+^ T_convs_; CD3^+^CD4^+^Foxp3^-^), effector memory CD4^+^ T cells (CD4^+^ T_EMs_; CD3^+^CD4^+^CD44^+^CD62L^-^), and T_regs_ (CD3^+^CD4^+^CD25^+^Foxp3^+^) was determined from the CD4 T cell/T_reg_ panel; the percentages that were CD8^+^ T cells (CD3^+^CD4^-^CD8^+^) and memory phenotype CD8^+^ T cells (CD8^+^ T_MPs_; CD3^+^CD4^-^CD8^+^CD44^+^CD122^+^) were determined from the CD8 T cell panel; the percentage that was NK cells (CD3^-^CD49b^+^NK1.1^+^) was determined from the NK cell panel (see gating strategy in Supplemental Figure 11). Data were analyzed using FlowJo ver. 10.6.1 software (Tree Star). The total number of splenocytes was acquired by measuring each single cell suspension prepared from spleen on an automated cell counter (EVE, NanoEnTek, Korea). The experiment was performed twice with similar results. Average ratios of the relative expansion of the indicated subsets are plotted.

### Tumor therapy studies in mice

BALB/c and C57BL/6 mice were s.c. inoculated with 2×10^5^ CT26 cells or 1×10^5^ B16F10 cells, respectively, in 100 μL of sterile PBS on the shaved right anterior flank. Tumor-bearing mice were randomly distributed into experimental groups, with approximately equal average body weight across groups. Treatments were administered on days 2, 4, and 7 (B16F10, Supplemental Figure 9E), days 2, 4, 6, and 10 (B16F10, Figure 7, D and F), or days 4, 6, 8, and 12 (CT26, Figure 7, E and G) after tumor inoculation. Mice were treated i.p. with: 0.665 mg/kg /dose of F10 IC or Control IC (molar equivalent of 0.125 mg/kg IL-2, approx. 13.3 µg IC) in 200 µL PBS, or with 200 µL of PBS alone (Figure 7, D-G); or with 0.125 mg/kg of IL-2/dose, the molar equivalent of 0.125 mg/kg IL-2/dose of IL-2/602 complex (prepared by incubating IL-2 and 602 Ab at a 2:1 molar ratio of cytokine:antibody in PBS for 15 min at room temperature), or molar equivalent of 0.125 mg/kg IL-2 /dose of F10 IC (approx. 13.3 µg of IC) in 200 µL of PBS (Supplemental Figure 9E). The treatment regimen was prophylactic, i.e., the first treatment was applied before the tumors were developed (diameter of approx. 2-3 mm). The tumors were measured by caliper, and size was calculated as (shortest diameter)^2^×(longest diameter)×0.5. The mice were euthanized on day 21 (B16F10) or 25 (CT26). No mice were excluded from analyses except those with unexpected death from nontumor reasons.

### Toxicity studies in mice

To assess the degree to which any treatment caused toxicity in the form of vascular leak syndrome (VLS), C57BL/6 or BALB/c mice were injected i.p. on days 1, 2, 3, and 4 with 1.5 µg/dose IL-2, the molar equivalent of 0.075 mg/kg IL-2/dose of IL-2/602 complex (prepared by incubating IL-2 and 602 Ab at a 2:1 cytokine:antibody molar ratio in PBS for 15 min at room temperature), the molar equivalent of 0.075 mg/kg IL-2/dose of 602 IC, F10 IC, or Control IC in 250 µL of PBS, or with 250 µL of PBS alone. Mice were randomly distributed into experimental groups, with approximately equal average body weight across groups. Mice were sacrificed on day 5 (24 hr after the last injection) by cervical dislocation, and lungs were harvested. The difference in the wet weight and dry weight of the lungs, before and after lyophilization (16-24 hr, Concentrator Plus, Eppendorf), respectively, was calculated. The water content of the lungs was measured as the dry weight of lungs subtracted from the wet weight of the lungs. For ALT and AST analysis, BALB/c mice were sacrificed on day 5 by carotid cut, and all blood was collected into heparin-treated blood-collection vials, serum isolated, and analyzed for AST (IFCC; Beckman Coulter, OSR6109) and ALT (IFCC; Beckman Coulter, OSR6107) using Beckman Coulter AU480 clinical chemistry analyzer (Beckman Coulter, Czech Republic).

### Statistics

Data were graphed and analyzed using GraphPad Prism v9.0 and expressed as mean ± standard error of the mean (SEM) for in vivo summary data, or as mean ± standard deviation (SD) for non-in vivo summary data, and all summary data were analyzed by one-way ANOVA with Tukey post hoc test. A *P* value of less than 0.05 was used as the threshold for statistical significance: *P<0.05, **P<0.01, ***P<0.001, ****P<0.0001. Complete statistical analyses are provided in data file S1.

### Study approval

Mice were housed and handled according to the institutional committee guidelines. Animal experiments were approved by the Animal Care and Use Committee of the Institute of Molecular Genetics and were in agreement with local legal requirements and ethical guidelines.

## List of Supplementary Materials

Fig. S1 to S11 Table S1 to S7

Data file S1

## Funding

Department of Defense grants W81XWH1810735 and W81XWH21P0031 (JBS) National Institutes of Health grants R21CA249381, R01EB029455, and R01EB029341 (JBS)

Maryland Stem Cell Research Fund grant 2019-MSCRFD-5039 (JBS) V Foundation Scholars award V2018-005 (JBS)

Emerson Collective Cancer Research Fund (JBS) Willowcroft Foundation (JBS)

Johns Hopkins University Catalyst award (JBS) Czech Science Foundation grant 20-13029S (JT, OV) Charles University grant GAUK 285423 (OV)

Institute of Biotechnology of the Czech Academy of Sciences grant RVO 86652036 (JT, TH, MH)

Institute of Microbiology of the Czech Academy of Sciences grant RVO 61388971 (JT, MK)

Project National Institute for Cancer Research Programme EXCELES, ID LX22NPO5102

- Funded by the European Union - Next Generation EU (JT)

Czech Centre for Phenogenomics at the Institute of Molecular Genetics grants RVO 68378050 and MEYS CR LM2018126 (JT, TH, MH)

Division of Intramural Research, National Heart, Lung, and Blood Institute (J-XL, PL, WJL)

Division of Intramural Research, National Cancer Institute (J-XL, PL, WJL) National Institutes of Health Training Grant K12 GM123914 01A1 (EKL) American Cancer Society postdoctoral research fellowship (JRG)

## Author contributions

Conceptualization: EKL, JT, JBS Methodology: EKL, JT, JRG, OV, WJL, JBS

Investigation: EKL, JT, JRG, MIL, J-XL, PL, MJP, ERJ, LT, SDC, TH, MH, OV

Visualization: EKL Resources: MK

Funding acquisition: JT, OV, MK, WJL, JBS Supervision: WJL, JBS

Writing – original draft: EKL, JRG, JBS Writing – review & editing: All authors

## Data and materials availability

Coordinates and structure factors for the IL-2/602 scFv and IL- 2/F10 scFv complexes are deposited in the Worldwide Protein Data Bank (wwPDB). All data associated with this study are present in the paper or the Supplementary Materials.

