## Supplementary Materials for "Engineered cytokine/antibody fusion proteins improve delivery of IL-2 to pro-inflammatory cells and promote antitumor activity"

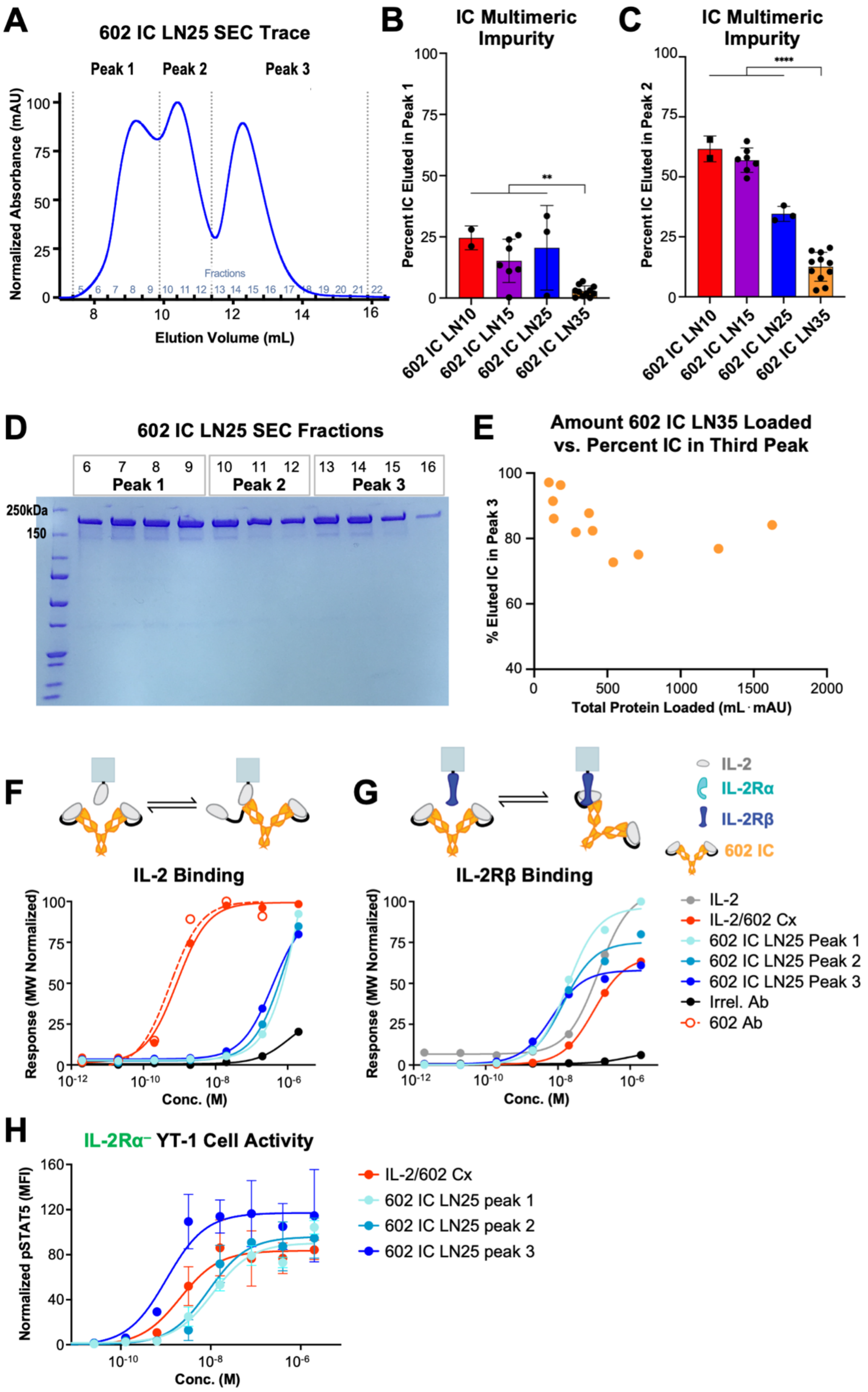


**Fig. S1. Optimization of immunocytokine linker length. (A)** Representative size-exclusion chromatography (SEC) trace shows the distribution of 602 IC LN25 in each of the 3 peaks, representing differential oligomeric states of IC. The average percentage of each IC that eluted in the first **(B)** and second **(C)** peaks based on the area under the SEC curve is shown. Data represent mean ± SD from at least 2, and up to 11, purifications. Statistical significance was determined by one-way ANOVA with Tukey's multiple comparison test and is noted only for comparisons to 602 IC LN35. **P<0.01, ****P<0.0001. **(D)** SEC elution fractions 6 to 16 of 602 IC LN25 (as annotated in (A)) show identical migration by SDS-PAGE under non-reducing conditions. **(E)** The percentage of IC that eluted in the third peak compared to the total amount of 602 IC LN35 that was loaded per SEC run. Total amount of protein was measured as the total area under all SEC three peaks (500 mL·mAU is approximately equal to a loading concentration of 7.5 µM. **(F)** Equilibrium bio-layer interferometry (BLI) titrations of soluble IL-2/602 complex (Cx) (1:1 molar ratio), 602 antibody (Ab), and 602 IC LN25 peaks 1, 2, and 3 against immobilized IL-2. An antibody with irrelevant specificity is included as a negative control. **(G)** Equilibrium BLI titrations of soluble IL-2, IL-2/602 Cx (1:1 cytokine:antibody molar ratio), 602 antibody (Ab), and 602 IC LN25 peaks 1, 2, and 3 against immobilized IL-2Rβ. An antibody with irrelevant specificity (Irrel. Ab) is included as a negative control. **(H)** STAT5 phosphorylation response of IL-2Rα$^-$ YT-1 human NK cells treated with IL-2/602 Cx (1:1 molar ratio) and 602 IC LN25 peaks 1, 2, and 3. Data represent mean ± SD (n=2).

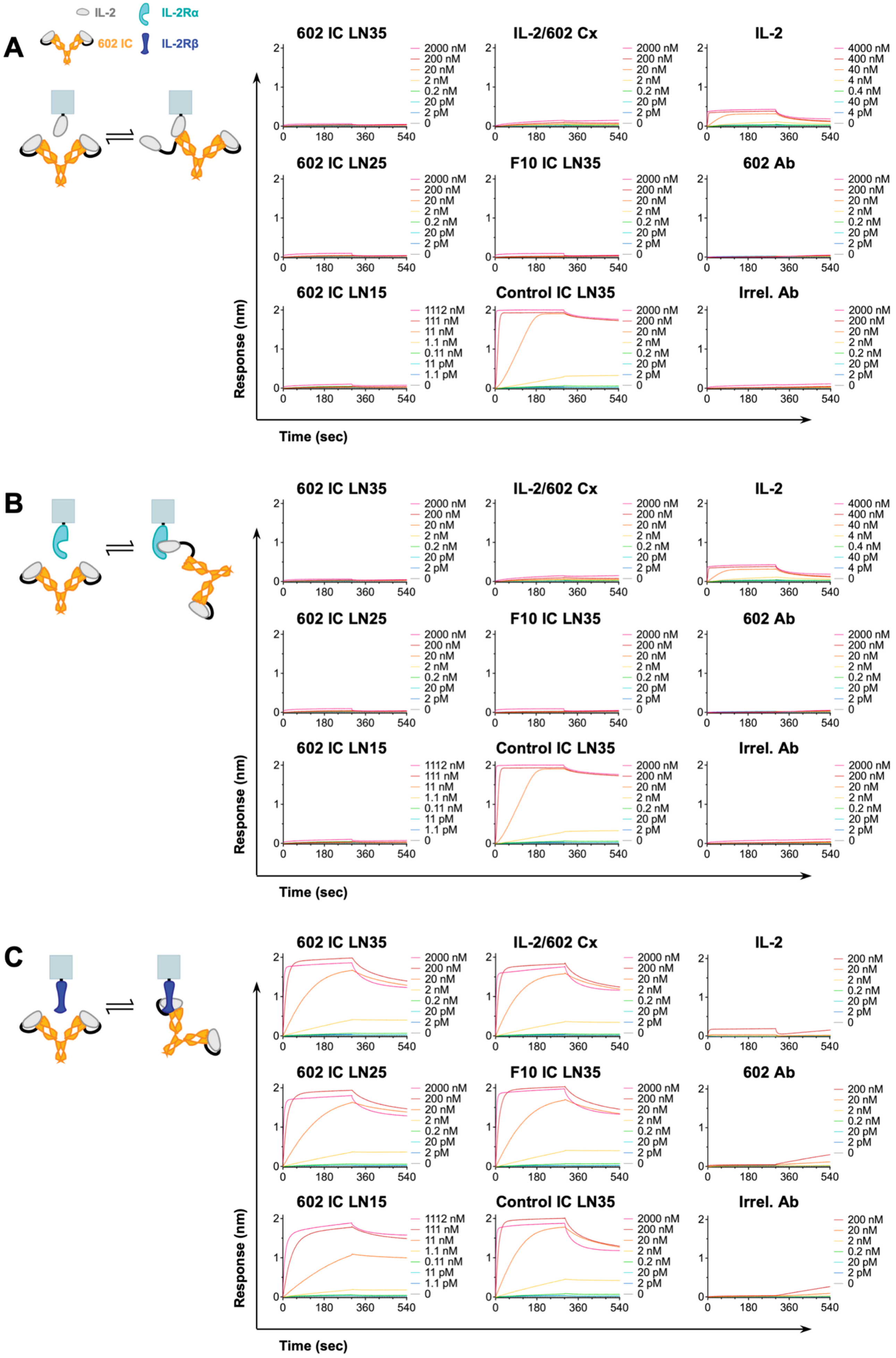

A
IL-2
IL-2Rα
602 IC
IL-2Rβ
602 IC LN35
IL-2/602 Cx
IL-2
602 IC LN25
F10 IC LN35
602 Ab
602 IC LN15
Control IC LN35
Irrel. Ab
2000 nM
200 nM
20 nM
2 nM
0.2 nM
20 pM
2 pM
0
4000 nM
400 nM
40 nM
4 nM
0.4 nM
40 pM
4 pM
1112 nM
111 nM
11 nM
1.1 nM
0.11 nM
11 pM
1.1 pM
Response (nm)
Time (sec)
0
180
360
540
B
602 IC LN35
IL-2/602 Cx
IL-2
602 IC LN25
F10 IC LN35
602 Ab
602 IC LN15
Control IC LN35
Irrel. Ab
Response (nm)
Time (sec)
C
602 IC LN35
IL-2/602 Cx
IL-2
602 IC LN25
F10 IC LN35
602 Ab
602 IC LN15
Control IC LN35
Irrel. Ab
200 nM
20 nM
2 nM
0.2 nM
20 pM
2 pM
0
Response (nm)
Time (sec)

**Fig. S2. Interaction of IC formulations with IL-2 cytokine and receptor subunits.** Bio-layer interferometry (BLI) kinetic traces depicting soluble IL-2, IL-2/602 complex (Cx) (2:1 cytokine:antibody molar ratio), 602 antibody (Ab), 602 IC LN15, LN25, and LN35, F10 IC, Control IC, or an antibody of irrelevant specificity (Irrel. Ab) binding to immobilized IL-2 **(A)**, IL-2Rα **(B)**, and IL-2Rβ **(C)** are shown. Equilibrium titrations corresponding to these traces are presented in **Fig. 2** and **Fig. 3**.

**Table S1.** IC variant and IL-2, IL-2Rα and IL-2Rβ binding properties.

| Immobilized | Soluble | Kinetic Fit Values | | | Equilibrium |
|---|---|---|---|---|---|
| | | $K_D$ (nM) | $k_{on}$ ($\times 10^4$ 1/Ms) | $k_{off}$ ($\times 10^{-4}$ 1/s) | $K_D$ (nM) |
| IL-2 | IL-2 | ND | ND | ND | >2000 |
| | Control IC LN35 | ND | ND | ND | >2000 |
| | IL-2/602 Cx | 10.6 ± 0.1 | 3.41 ± 0.01 | 3.62 ± 0.03 | 103 ± 35 |
| | 602 IC LN15 | 88.3 ± 2.5 | 0.500 ± 0.005 | 4.41 ± 0.12 | ~ 905 |
| | 602 IC LN25 | 76.9 ± 2.4 | 0.448 ± 0.003 | 3.44 ± 0.10 | 941 ± 134 |
| | 602 IC LN35 | 143.2 ± 2.9 | 0.265 ± 0.002 | 3.79 ± 0.07 | >2000 |
| | F10 IC LN35 | 166.9 ± 8.6 | 0.240 ± 0.004 | 4.00 ± 0.19 | >2000 |
| | 602 Ab | 0.13 ± 0.02 | 38.2 ± 0.3 | 0.51 ± 0.09 | 6.3 ± 5.2 |
| | Irrel. Ab | ND | ND | ND | >2000 |
| IL-2Rα | IL-2 | 3.20 ± 0.08 | 201.2 ± 3.9 | 34.4 ± 0.3 | 7.3 ± 14.5 |
| | Control IC LN35 | ND | ND | ND | 5.2 ± 8.2 |
| | IL-2/602 Cx | 0.55 ± 0.09 | 42.1 ± 1.0 | 2.3 ± 0.4 | >2000 |
| | 602 IC LN15 | ND | ND | ND | >2000 |
| | 602 IC LN25 | ND | ND | ND | >2000 |
| | 602 IC LN35 | ND | ND | ND | >2000 |
| | F10 IC LN35 | ND | ND | ND | >2000 |
| | 602 Ab | ND | ND | ND | >2000 |
| | Irrel. Ab | ND | ND | ND | >2000 |
| IL-2Rβ | IL-2 | ND | ND | ND | 112 ± 303 |
| | Control IC LN35 | 0.18 ± 0.01 | 1554 ± 35 | 3.76 ± 0.15 | 4.3 ± 6.3 |
| | IL-2/602 Cx | 3.89 ± 0.04 | 41.0 ± 0.3 | 15.5 ± 0.1 | 5.2 ± 5.6 |
| | 602 IC LN15 | 2.20 ± 0.02 | 24.2 ± 0.1 | 5.11 ± 0.05 | 8.4 ± 1.9 |
| | 602 IC LN25 | 3.66 ± 0.03 | 27.6 ± 0.1 | 9.98 ± 0.07 | 5.3 ± 6.4 |
| | 602 IC LN35 | 4.22 ± 0.04 | 34.2 ± 0.2 | 14.2 ± 0.1 | 5.0 ± 5.9 |
| | F10 IC LN35 | 4.16 ± 0.04 | 31.4 ± 0.2 | 12.9 ± 0.1 | 5.3 ± 5.2 |
| | 602 Ab | ND | ND | ND | >2000 |
| | Irrel. Ab | ND | ND | ND | >2000 |

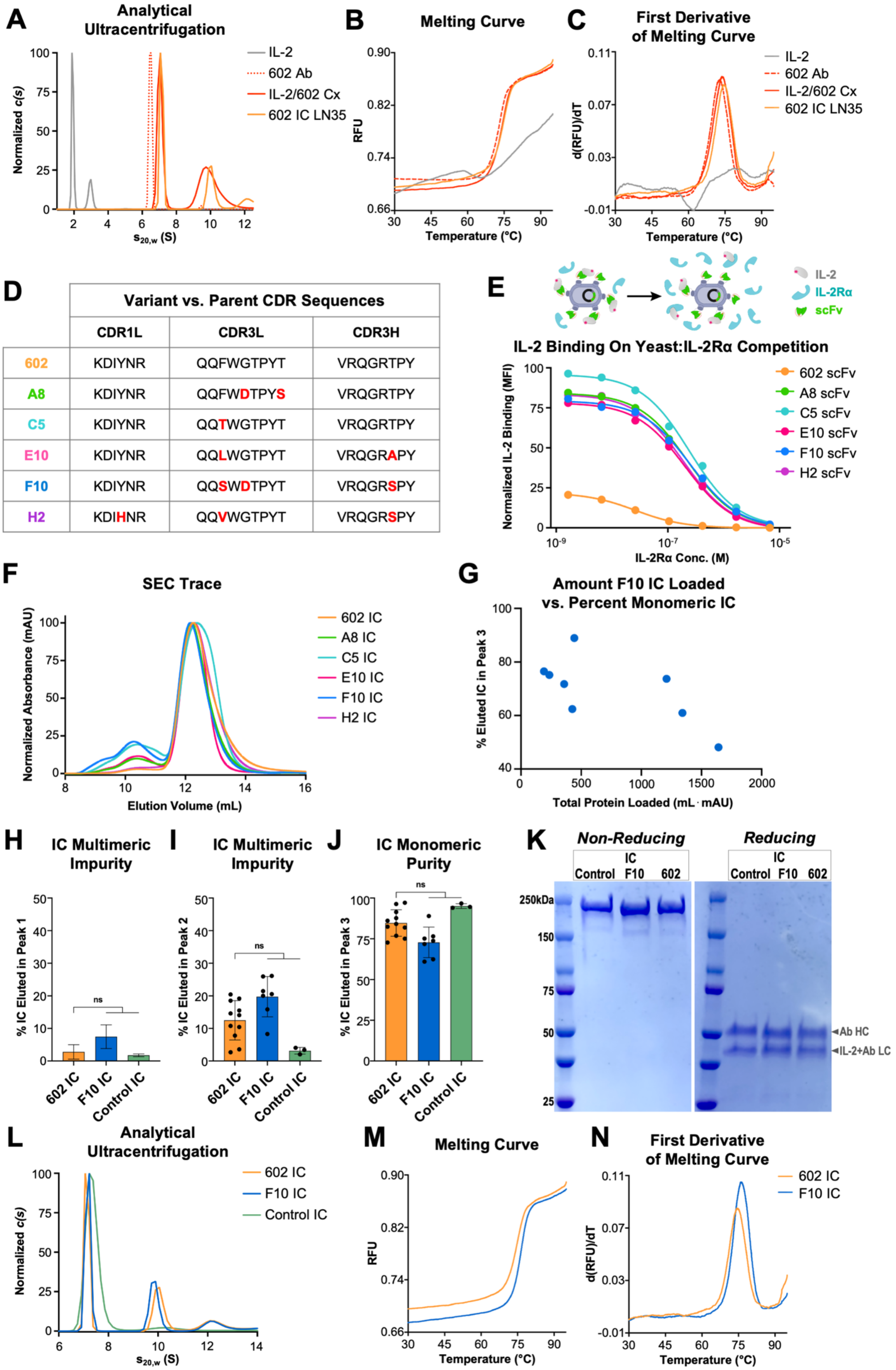
A
Analytical Ultracentrifugation
Normalized c(s)
s20,w (S)
IL-2
602 Ab
IL-2/602 Cx
602 IC LN35
B
Melting Curve
RFU
Temperature (°C)
C
First Derivative of Melting Curve
d(RFU)/dT
Temperature (°C)
IL-2
602 Ab
IL-2/602 Cx
602 IC LN35
D
Variant vs. Parent CDR Sequences
CDR1L CDR3L CDR3H
602 KDIYNR QQFWGTPYT VRQGRTPY
A8 KDIYNR QQFWDTPYS VRQGRTPY
C5 KDIYNR QQTWGTPYT VRQGRTPY
E10 KDIYNR QQLWGTPYT VRQGRAPY
F10 KDIYNR QQSWDTPYT VRQGRSPY
H2 KDIHNR QQVWGTPYT VRQGRSPY
E
IL-2
IL-2Rα
scFv
IL-2 Binding On Yeast:IL-2Rα Competition
Normalized IL-2 Binding (MFI)
IL-2Rα Conc. (M)
602 scFv
A8 scFv
C5 scFv
E10 scFv
F10 scFv
H2 scFv
F
SEC Trace
Normalized Absorbance (mAU)
Elution Volume (mL)
602 IC
A8 IC
C5 IC
E10 IC
F10 IC
H2 IC
G
Amount F10 IC Loaded vs. Percent Monomeric IC
% Eluted IC in Peak 3
Total Protein Loaded (mL · mAU)
H
IC Multimeric Impurity
% IC Eluted in Peak 1
I
IC Multimeric Impurity
% IC Eluted in Peak 2
J
IC Monomeric Purity
% IC Eluted in Peak 3
ns
602 IC
F10 IC
Control IC
K
Non-Reducing
Reducing
IC
Control F10 602
250kDa
150
75
50
25
Ab HC
IL-2+Ab LC
L
Analytical Ultracentrifugation
Normalized c(s)
s20,w (S)
602 IC
F10 IC
Control IC
M
Melting Curve
RFU
Temperature (°C)
N
First Derivative of Melting Curve
d(RFU)/dT
Temperature (°C)
602 IC
F10 IC

**Fig. S3. Characterization of representative 602 variants from evolved mutagenic library. (A)** Analytical ultracentrifugation analysis of unconjugated IL-2, 602 antibody (Ab), IL-2/602 complex (Cx) (2:1 cytokine:antibody molar ratio), and 602 IC LN35. 602 IC LN35 and IL-2/602 Cx show main species sedimenting at 6.5–7.5 S, corresponding to the expected size. Minor peaks are also observed for both, representing dimeric and trimeric species. Differential scanning fluorometry (DSF) profiles **(B)** and their first derivatives **(C)** for the thermal unfolding of IL-2, 602 Ab, and IL-2/602 Cx (2:1 cytokine:antibody molar ratio), and 602 IC LN35 in HBS. **(D)** Mutations to the CDRs for each of the five mutagenic 602 scFv library-derived variants (A8, C5, E10, F10, and H2) are indicated in red. **(E)** IL-2 binding (5 nM) to yeast-displayed 602 scFv compared to the five selected 602 scFv variants in the presence of titrated concentrations of IL-2Rα. Selected yeast-displayed 602 scFv variants show stronger IL-2 binding at higher IL-2Rα concentrations compared to parent 602 antibody, demonstrating superior receptor competition. **(F)** Representative size-exclusion chromatography (SEC) traces show that the majority of variant ICs elutes in the third peak, representing the monomeric construct. **(G)** Percentage of IC that eluted in the third peak versus the total amount of F10 IC loaded per SEC run. Total amount of protein was measured as the total area under all three SEC peaks (500 mL·mAU is approximately equivalent to a loading concentration of 7.5 μM). The average percentage of each IC (602, F10 and Control) that eluted in the first **(H)**, second **(I)**, and third **(J)** peaks based on the computed area under the SEC curve. Data represent mean ± SD from at least 2, and up to 11, purifications. Statistical significance was determined by one-way ANOVA with Tukey's multiple comparison test. ns, not significant. **(K)** 602 IC, F10 IC, and Control IC migrate at the expected sizes by SDS-PAGE, under non-reducing (~180 kDa) and reducing (49 kDa for HC, ~41 for IC LC) conditions. **(L)** Analytical ultracentrifugation analysis of 602 IC, F10 IC, and Control IC. ICs show main species sedimenting at 6.5–7.5 S, corresponding to the expected size for each. Minor peaks are also observed, representing dimeric and trimeric species. DSF profile **(M)** and its first derivative **(N)** for the thermal unfolding of 602 IC, and F10 IC in HBS.

**Table S2**. $T_m$ values of IL-2, IL-2/602 Cx, 602 Ab, 602 IC and F10 IC.

| **Sample** | **$T_m$ (°C)** |
|---|---|
| | **HBS** |
| IL-2 | 61.3* |
| 602 Ab | 73.1 |
| IL-2/602 Cx | 74.2 |
| 602 IC | 74.9 |
| F10 IC | 76.5 |

**Table S3**. EC50 values for 602 IC linker length variants signaling on IL-2Rα$^{+}$ or IL-2Rα$^{-}$ YT-1 cells.

| **Treatment** | **EC$_{50}$ (nM)** | |
|---|---|---|
| | **IL-2Rα$^{+}$** | **IL-2Rα$^{-}$** |
| IL-2 | 0.11 ± 0.35 | 0.34 ± 0.10 |
| IL-2/602 Cx | 0.25 ± 0.15 | 0.81 ± 1.18 |
| 602 IC LN15 | 38 ± 16 | 78 ± 67 |
| 602 IC LN25 | 5.1 ± 4.5 | 2.8 ± 1.4 |
| 602 IC LN35 | 2.3 ± 1.0 | 0.65 ± 1.44 |

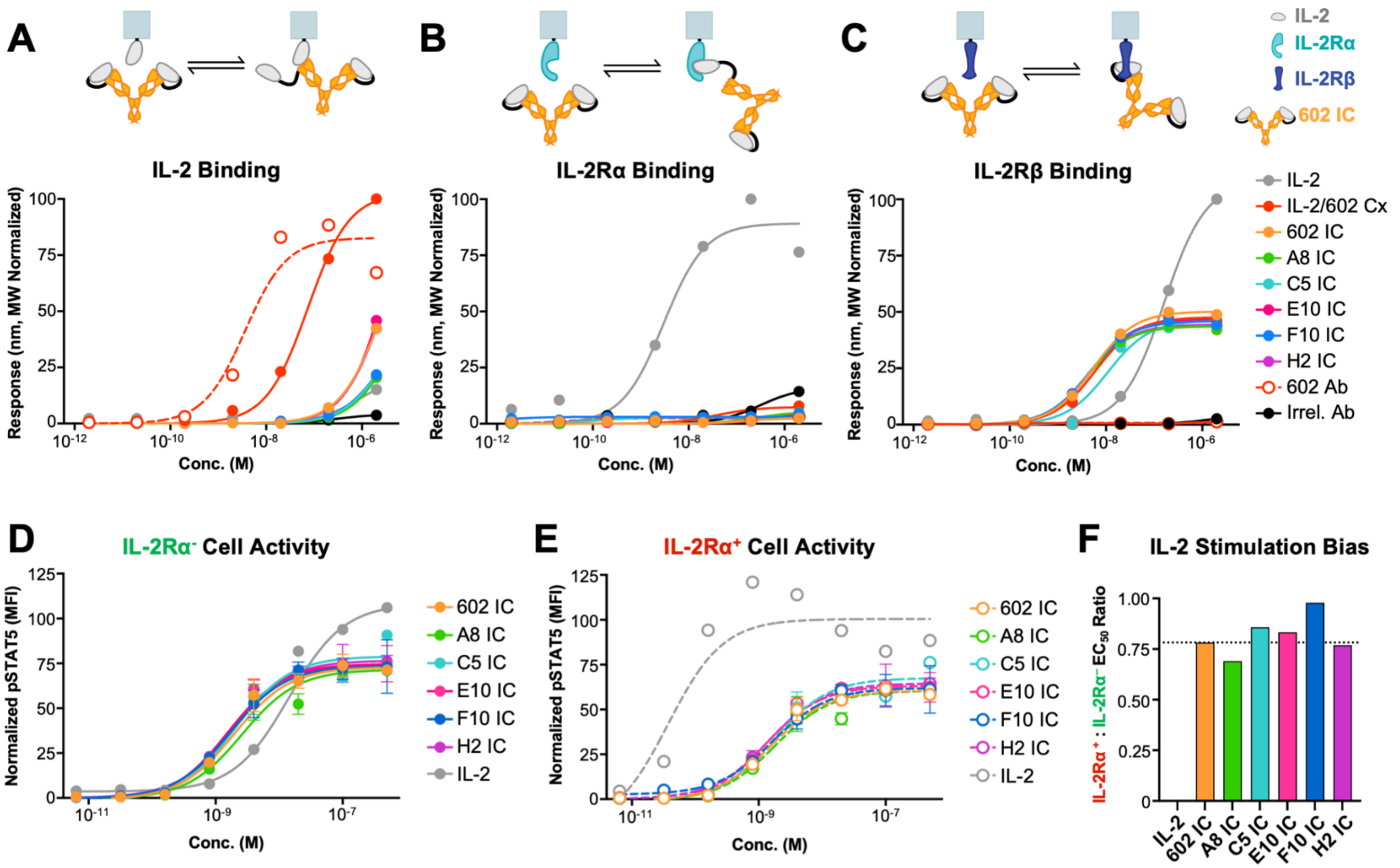


**Fig. S4. Functional activities of representative engineered 602 IC variants.** Equilibrium bio-layer interferometry (BLI) titrations of soluble IL-2, IL-2/602 complex (Cx) (2:1 cytokine:antibody molar ratio), 602 antibody (Ab), 602 IC, 5 engineered 602 IC variants (A8, C5, E10, F10, and H2), and an antibody of irrelevant specificity (Irrel. Ab) binding to immobilized IL-2 **(A)**, immobilized IL-2Rα **(B)**, and immobilized IL-2Rβ **(C)**. **(D)** STAT5 phosphorylation response of IL-2Rα$^-$ YT-1 human NK cells treated with IL-2, 602 IC, and engineered 602 IC variants in a mixed IL-2Rα$^-$/IL-2Rα$^+$ cell assay. Data represent mean ± SD (n=2). **(E)** STAT5 phosphorylation response of IL-2Rα$^+$ YT-1 human NK cells treated with IL-2, 602 IC, F10 IC, and engineered 602 IC variants in a mixed IL-2Rα$^-$/IL-2Rα$^+$ cell assay. Data represent mean ± SD for at least 420 collected flow cytometry events (avg. 930 events) (n=2). **(F)** Ratio of the IL-2Rα$^+$/IL-2Rα$^-$ STAT5 phosphorylation $EC_{50}$ values for assays shown in (D) and (E).

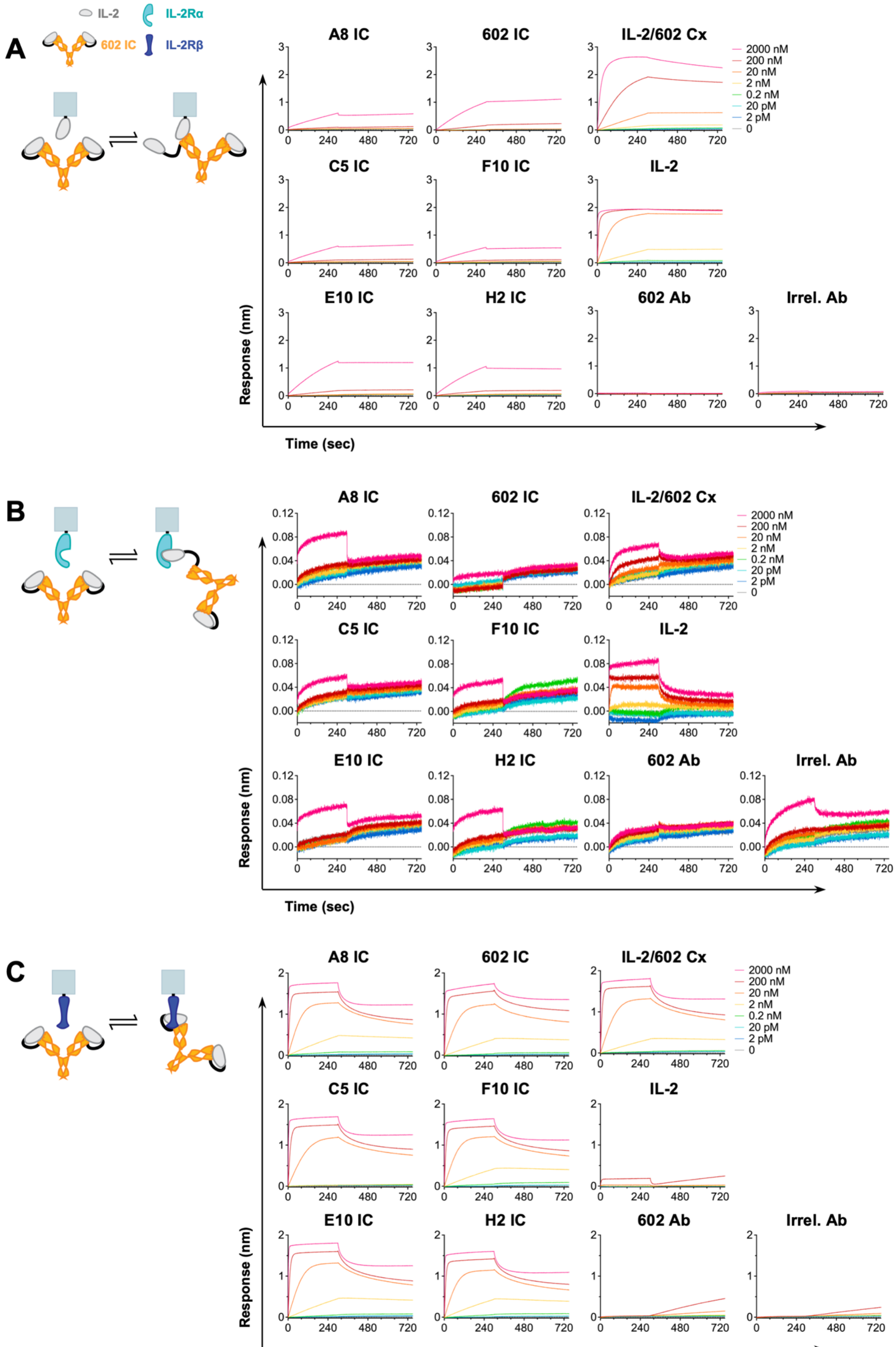
A
IL-2
IL-2Rα
602 IC
IL-2Rβ
A8 IC
602 IC
IL-2/602 Cx
C5 IC
F10 IC
IL-2
E10 IC
H2 IC
602 Ab
Irrel. Ab
2000 nM
200 nM
20 nM
2 nM
0.2 nM
20 pM
2 pM
0
Response (nm)
Time (sec)
B
A8 IC
602 IC
IL-2/602 Cx
C5 IC
F10 IC
IL-2
E10 IC
H2 IC
602 Ab
Irrel. Ab
2000 nM
200 nM
20 nM
2 nM
0.2 nM
20 pM
2 pM
0
Response (nm)
Time (sec)
C
A8 IC
602 IC
IL-2/602 Cx
C5 IC
F10 IC
IL-2
E10 IC
H2 IC
602 Ab
Irrel. Ab
2000 nM
200 nM
20 nM
2 nM
0.2 nM
20 pM
2 pM
0
Response (nm)

**Fig. S5. Interaction of engineered 602 IC variants with IL-2 cytokine and receptor subunits.** Bio-layer interferometry (BLI) kinetic traces depicting soluble IL-2, IL-2/602 complex (Cx) (1:1 cytokine:antibody molar ratio), 602 antibody (Ab), 602 IC, and five engineered 602 IC variants (A8, C5, E10, F10, and H2) binding to immobilized IL-2 **(A)**, IL-2Rα **(B)**, and IL-2Rβ **(C)**.

**Table S4.** EP602 variants and IL-2 and IL-2Rβ binding properties.

| Immobilized | Soluble | Kinetic Fit Values | | | Equilibrium |
|---|---|---|---|---|---|
| | | $K_D$ (nM) | $k_{on}$ ($\times 10^4$ 1/Ms) | $k_{off}$ ($\times 10^{-4}$ 1/s) | $K_D$ (nM) |
| IL-2 | IL-2 | ND | ND | ND | >2000 |
| | IL-2/602 Cx | 13.27 ± 0.09 | 2.52 ± 0.01 | 3.06 ± 0.02 | 74 ± 20 |
| | 602 IC | ND | ND | ND | >2000 |
| | A8 IC | ND | ND | ND | >2000 |
| | C5 IC | ND | ND | ND | >2000 |
| | E10 IC | 20.3 ± 1.8 | 0.192 ± 0.002 | 0.39 ± 0.03 | >2000 |
| | F10 IC | ND | ND | ND | >2000 |
| | H2 IC | 50.5 ± 2.5 | 0.221 ± 0.002 | 1.11 ± 0.05 | >2000 |
| | 602 Ab | 0.044 ± 0.05 | 64.2 ± 0.3 | 0.28 ± 0.03 | 3.7 ± 7.4 |
| | Irrel. Ab | ND | ND | ND | >2000 |
| IL-2Rβ | IL-2 | ND | ND | ND | 161 ± 28 |
| | IL-2/602 Cx | 0.30 ± 0.02 | 38.7 ± 1.8 | 1.17 ± 0.04 | 6.0 ± 2.6 |
| | 602 IC | 0.26 ± 0.04 | 28.8 ± 2.8 | 0.75 ± 0.07 | 5.4 ± 1.2 |
| | A8 IC | 0.52 ± 0.01 | 52.6 ± 1.0 | 2.74 ± 0.02 | 4.2 ± 1.1 |
| | C5 IC | 2.38 ± 0.04 | 48.7 ± 0.7 | 11.57 ± 0.12 | 11 ± 12 |
| | E10 IC | 0.84 ± 0.04 | 26.2 ± 1.3 | 2.20 ± 0.03 | 4.9 ± 0.7 |
| | F10 IC | 1.45 ± 0.23 | 9.7 ± 1.5 | 1.41 ± 0.04 | 4.4 ± 1.2 |
| | H2 IC | 5.1 ± 1.2 | 6.0 ± 1.4 | 3.03 ± 0.04 | 4.3 ± 0.7 |
| | 602 Ab | ND | ND | ND | >2000 |
| | Irrel. Ab | ND | ND | ND | >2000 |

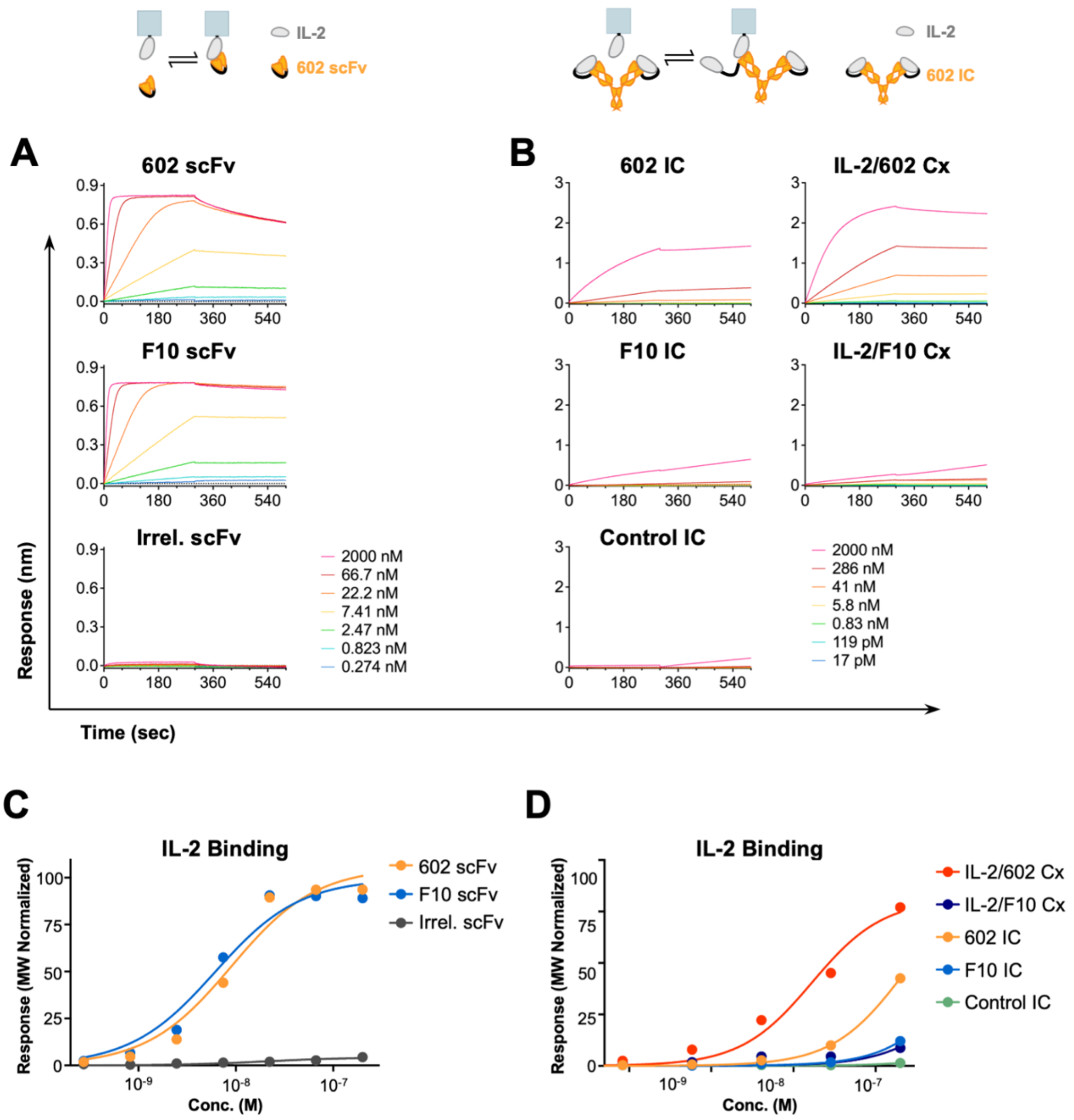


**Fig. S6. Interaction of 602 and F10 with IL-2 cytokine.** Bio-layer interferometry (BLI) kinetic traces **(A and B)** and equilibrium titrations **(C and D)** of scFvs, IL-2/antibody complexes (Cx) and ICs binding to immobilized IL-2. The irrelevant scFv was an anti-mPD-1 scFv (clone [RMP1.14]). IL-2/antibody complexes were prepared by mixing IL-2 with the indicated antibody at a 2:1 molar ratio.

**Table S5.** IL-2 binding properties of F10 and 602.

| Immobilized | Soluble | Kinetic Fit Values | | | Equilibrium |
|---|---|---|---|---|---|
| | | $K_D$ (nM) | $k_{on}$ ($\times 10^5$ 1/Ms) | $k_{off}$ ($\times 10^{-4}$ 1/s) | $K_D$ (nM) |
| IL-2 | 602 scFv | 1.94 ± 0.02 | 3.61 ± 0.04 | 6.70 ± 0.04 | 9.00 ± 8.59 |
| | F10 scFv | 0.34 ± 0.01 | 5.63 ± 0.03 | 1.88 ± 0.04 | 6.12 ± 5.66 |
| | Irrel. scFv | ND | ND | ND | >200 |
| | IL-2/602 Cx | <0.001 | 13.0 ± 0.3 | <0.001 | 168 ± 237 |
| | IL-2/F10 Cx | <0.001 | 36.5 ± 0.7 | <0.001 | >2000 |
| | 602 IC | <0.001 | 0.51 ± 0.02 | <0.001 | >2000 |
| | F10 IC | ND | ND | ND | >2000 |
| | Control IC | ND | ND | ND | >2000 |

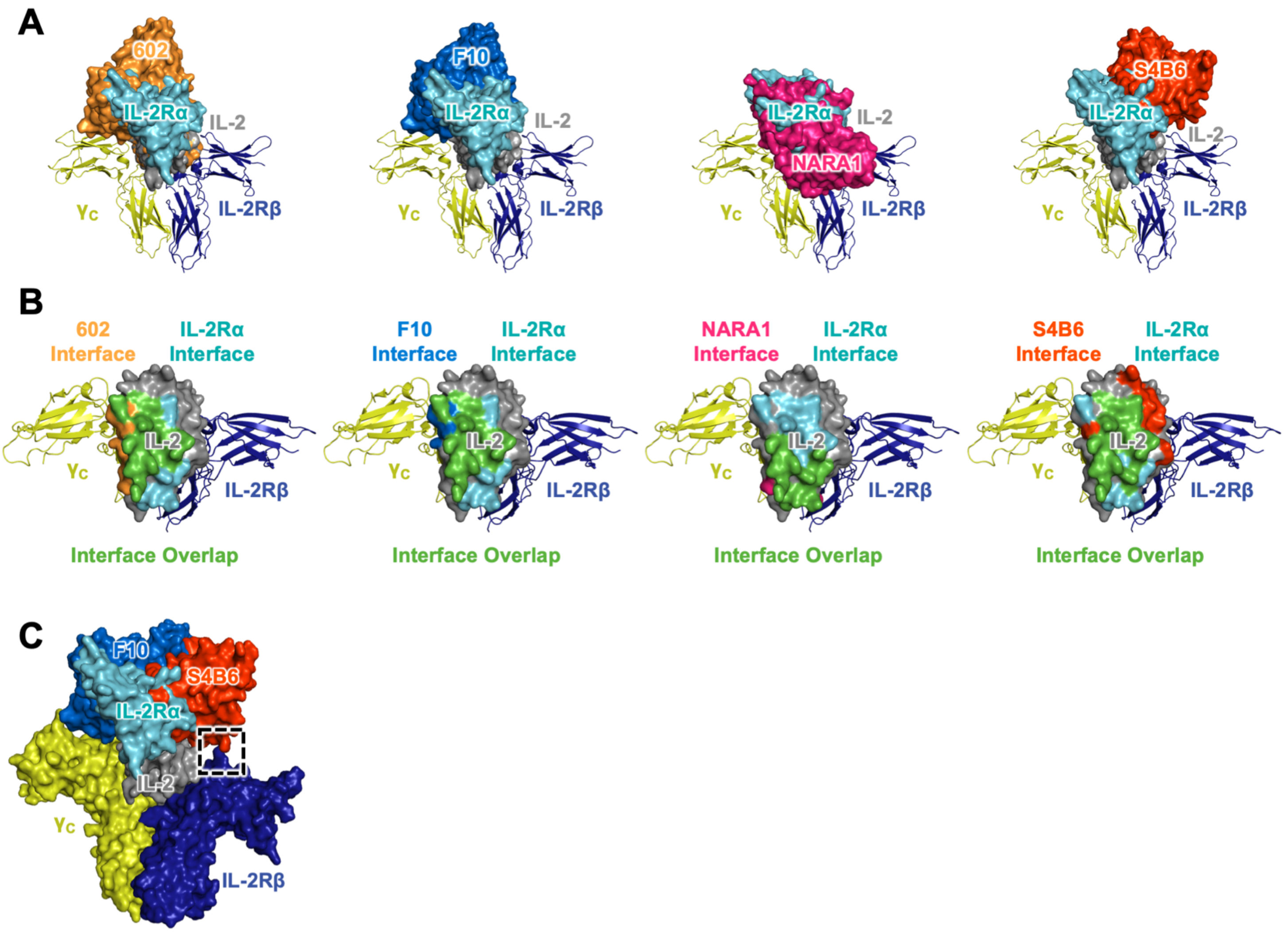


**Fig. S7. Comparison of 602, F10, NARA1, and S4B6 antibody binding topologies. (A)** Overlay of the IL-2/antibody complex structures for 602, F10, NARA1, and S4B6 (only variable domains of the antibodies are shown) and the quaternary IL-2 cytokine/receptor complex structure (PDB 2B5I). **(B)** ‘‘Top-down’’ views of the PISA-predicted binding interface between IL-2Rα and IL-2 (cyan), and between IL-2 and the 602 (yellow-orange), F10 (blue), NARA1 (pink), and S4B6 (orange) antibodies. The binding interface for S4B6 was predicted based on alignment of the mouse IL-2/S4B6 Fab structure (PDB 4YUE) to human IL-2 bound to its high affinity trimeric receptor complex (PDB 2B5I). Overlap between the IL-2Rα and antibody binding interfaces are shown in green. **(C)** Alignment of the mouse IL-2/S4B6 complex (only antibody variable domains are shown) overlayed with the resolved crystallographic structure of hIL-2 bound to its high affinity trimeric receptor complex (PDB 2B5I) and the resolved hIL-2/F10 crystal structure. A steric clash was observed between the S4B6 antibody and IL-2Rβ (dashed black box), but not between the F10 scFv and IL-2Rβ due to the latter molecule’s shift away from the IL-2Rβ subunit.

**Table S6.** EC50 values for IC variant signaling on IL-2Rα$^{+}$ or IL-2Rα$^{-}$ YT-1 cells, and various PBMC subsets.

| **Treatment** | **$EC_{50}$ (nM)** | | | | |
|---|---|---|---|---|---|
| | **IL-2Rα$^{+}$ YT-1** | **IL-2Rα$^{-}$ YT-1** | **CD3$^{+}$CD4$^{+}$ CD8$^{-}$ FoxP3$^{+}$ PBMC** | **CD3$^{+}$CD4$^{+}$ CD8$^{-}$ FoxP3$^{-}$ PBMC** | **CD3$^{+}$CD4$^{-}$ CD8$^{+}$ PBMC** |
| IL-2 | 0.09 ± 0.07 | 0.36 ± 0.06 | 3.7E-4 ± 1.54E-3 | 0.051 ± 0.044 | 0.71 ± 1.00 |
| IL-2/602 Cx | 0.03 ± 0.02 | 0.41 ± 0.09 | 2.5E-4 ± 2.9E-4 | 0.024 ± 0.036 | 1.63 ± 4.59 |
| 602 IC | 0.17 ± 0.13 | 0.44 ± 0.11 | 1.1E-3 ± 2.0E-3 | 0.13 ± 0. 46 | 2.1 ± 1.8 |
| F10 IC | 1.30 ± 0.69 | 0.49 ± 0.24 | 1.17 ± 1.11 | 26 ± 12 | 10.6 ± 4.1 |
| Control IC | 1.28 ± 0.50 | 0.46 ± 0.14 | 3.6 ± 3.7 | 42 ± 27 | 14.5 ± 6.5 |

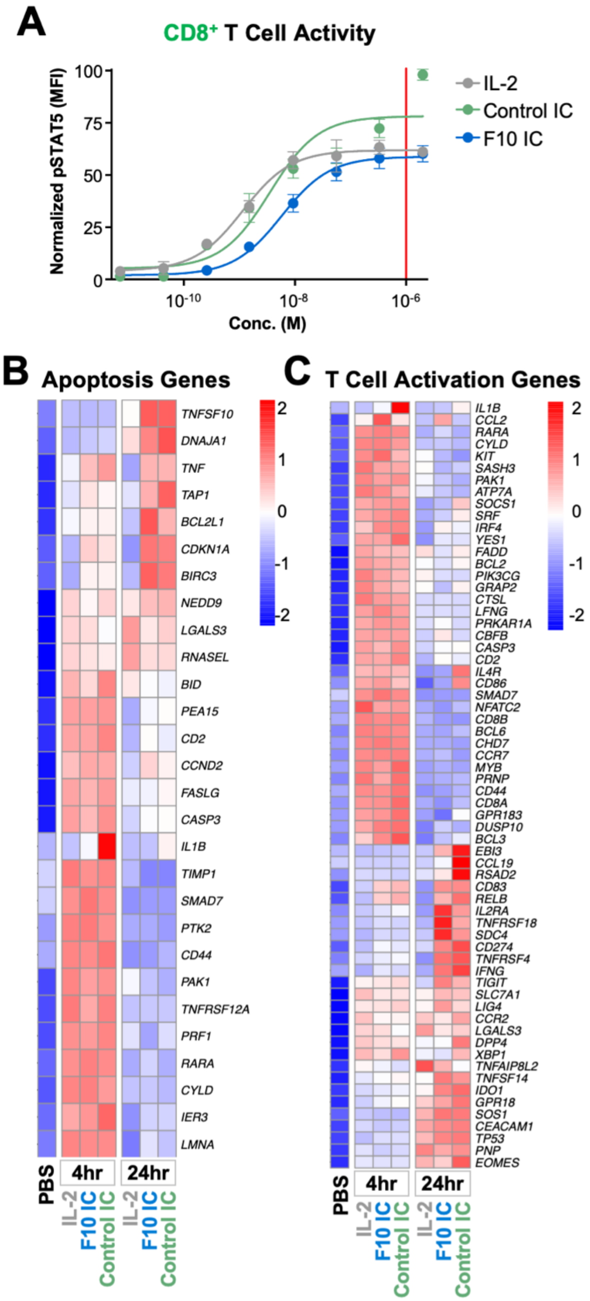


**Fig. S8. Unconjugated IL-2 and ICs induce similar gene expression profiles in human CD8$^+$ T cells. (A)** STAT5 phosphorylation response of human PBMC-derived CD8$^+$ T cells treated with IL-2, F10 IC, and Control IC. Data represent mean ± SD (n=3). Data represent mean ± SD for cells from two healthy donors (same donors that were used for RNA-Seq studies). The red line indicates the dose used for stimulation in RNA-Seq experiments (1 µM IL-2 or 0.5 µM IC). **(B and C)** RNA-Seq heatmap analysis for human CD8$^+$ T cells treated with PBS or the saturating dose of IL-2 (1 µM), F10 IC (0.5 µM), and Control IC (0.5 µM) for either 4 and 24 hr. Genes related to apoptosis (B) and T cell activation (C) are presented. The color scales in the heatmaps represent Z-score values ranging from blue (low expression) to red (high expression).

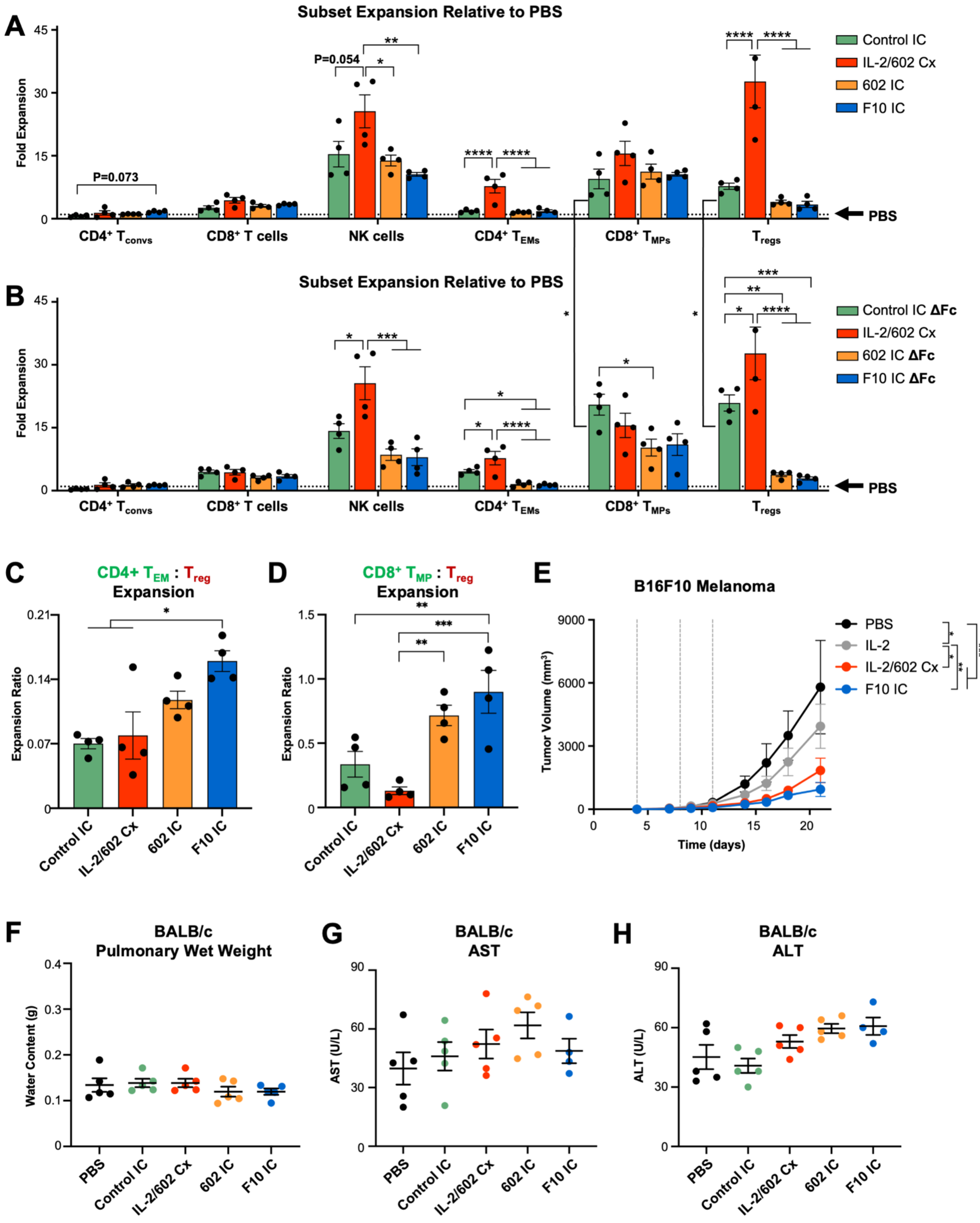

A
Subset Expansion Relative to PBS
Fold Expansion
P=0.073
P=0.054
CD4+ Tconvs
CD8+ T cells
NK cells
CD4+ TEMs
CD8+ TMPs
Tregs
PBS
Control IC
IL-2/602 Cx
602 IC
F10 IC
B
Subset Expansion Relative to PBS
Control IC ΔFc
IL-2/602 Cx
602 IC ΔFc
F10 IC ΔFc
C
CD4+ TEM : Treg
Expansion
Expansion Ratio
Control IC
IL-2/602 Cx
602 IC
F10 IC
D
CD8+ TMP : Treg
Expansion
E
B16F10 Melanoma
Tumor Volume (mm3)
Time (days)
PBS
IL-2
IL-2/602 Cx
F10 IC
F
BALB/c
Pulmonary Wet Weight
Water Content (g)
PBS
G
BALB/c
AST
AST (U/L)
H
BALB/c
ALT
ALT (U/L)

**Fig. S9. Engineered F10 IC promotes selective expansion of immune effector cells and improves the therapeutic efficacy of IL-2 without inducing toxicity. (A to D)** C57BL/6 mice (n=4) were injected intraperitoneally daily for 4 days with the molar equivalent of 0.125 mg/kg IL-2/dose of Control IC, IL-2/602 complex (Cx) (2:1 cytokine:antibody molar ratio), 602 IC, or F10 IC. Mice were sacrificed on day 5, and spleens were harvested. The total cell counts of conventional T cells ($CD4^+$ $T_{convs}$, $CD3^+CD4^+FoxP3^-$), $CD8^+$ T cells ($CD3^+CD4^-CD8^+$), natural killer (NK) cells ($CD3^-NK1.1^+CD49b^+$), $CD4^+$ effector memory T cells ($CD4^+$ $T_{EMs}$, $CD3^+CD4^+CD8^-CD44^+CD62L^-$), $CD8^+$ memory phenotype T cells ($CD8^+$ $T_{MPs}$, $CD3^+CD4^-CD8^+CD44^+CD122^+$), and T regulatory cells ($T_{regs}$, $CD3^+CD4^+CD25^+Foxp3^+$) in each spleen were determined by flow cytometry. The cell subset counts for each treatment divided by the cell subset count for PBS is shown (wild type antibody Fc regions in (A), antibody Fc regions with effector function knockout [ΔFc] in B), as are the ratios of $CD4^+$ $T_{EMs}$ to $T_{regs}$ (C) and $CD8^+$ $T_{MPs}$ to $T_{regs}$ (D). The dotted line in (A) indicates equivalent expansion to PBS treatment. **(E)** C57BL/6 mice (n=8) were injected subcutaneously s.c. with $1\times10^5$ B16F10 tumor cells and treated intraperitoneally on days 2, 4, and 7 with PBS, 0.125 mg/kg IL-2, or the molar equivalent of 0.125 mg/kg IL-2 of IL-2/602 complex (Cx) (2:1 cytokine:antibody molar ratio) or F10 IC. Tumor size is shown. **(F to H)** BALB/c mice (n=4-5/group) were injected daily for 4 days with PBS, 0.075 mg/kg IL-2/dose, or the molar equivalent of 0.075 mg/kg IL-2/dose of Control IC, IL-2/602 complex (Cx) (2:1 cytokine:antibody molar ratio), 602 IC, or F10 IC. Mice were sacrificed on day 5, and pulmonary wet weight **(**H), as well as serum concentrations of AST (F) and ALT (G) were measured. Data are shown as means ± SEM. Statistical significance was determined by one-way ANOVA with Tukey's multiple comparison test for immune cell subset expansion data and by two-way ANOVA test for tumor growth data: *P<0.05, **P<0.01, ***P<0.001, ****P<0.0001. No significant differences were observed in AST, ALT, or pulmonary wet weight by one-way ANOVA with Tukey's multiple comparison test.

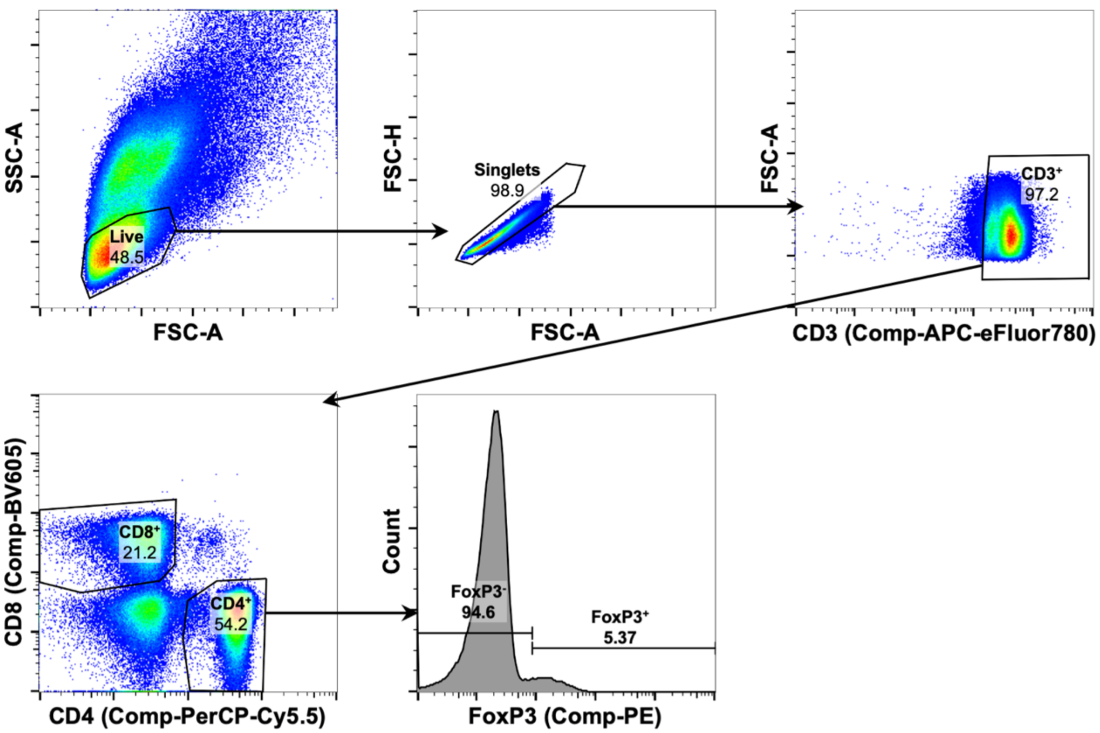


**Fig. S10. Flow cytometry gating strategy for human PBMC signaling assays.** CD4$^+$ T$_{convs}$ were gated as Live> Singlets> CD3$^+$> CD4$^+$> FoxP3$^-$, T$_{regs}$ were gated as Live> Singlets> CD3$^+$> CD4$^+$> FoxP3$^+$, and CD8$^+$ T cells were gated as Live> Singlets> CD3$^+$> CD4$^-$CD8$^+$.

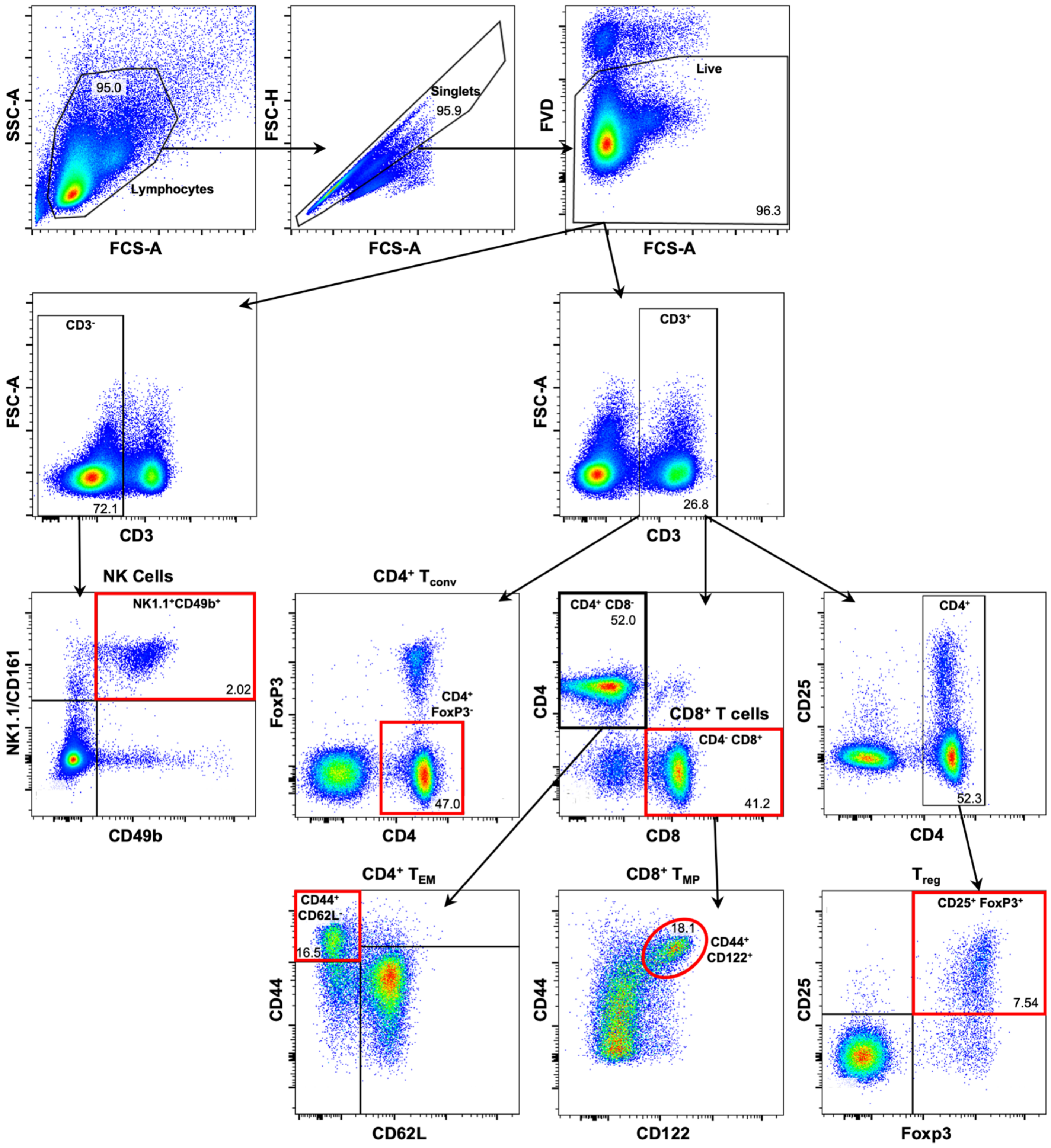


**Fig. S11. Flow cytometry gating strategy for mouse immune cell subset expansion assays.** All subsets were derived from the Lymphocytes> Singlets> Live gate. Within this gate, NK cells were gated as $CD3^{-}$> NK1.1$^{+}$ CD49b$^{+}$, CD4$^{+}$ $T_{convs}$ were gated as CD3$^{+}$> CD4$^{+}$FoxP3$^{-}$, CD4$^{+}$ $T_{EMs}$ were gated as CD3$^{+}$> CD4$^{+}$CD8$^{-}$> CD44$^{+}$CD62L$^{-}$, CD8$^{+}$ T cells were gated as CD3$^{+}$> CD4$^{-}$CD8$^{+}$, CD8$^{+}$

$T_{MPs}$ were gated as $CD3^+$> $CD4^-CD8^+$> $CD44^+CD122^+$, and $T_{regs}$ were gated as $CD3^+$> $CD4^+$> $CD25^+FoxP3^+$.
