## Supplementary material for "Engineered cytokine/antibody fusion proteins improve delivery of IL-2 to pro-inflammatory cells and promote antitumor activity": Data File S1

### Data file S1. Statistical Analy

| Statistical test | Measurement | Comparison |
|---|---|---|
| One-way ANOVA (Tukey post hoc test) | IC Monomeric Purity: % IC Eluted in Peak 3 | 602 IC LN35 vs. 602 IC LN25 |
| One-way ANOVA (Tukey post hoc test) | IC Monomeric Purity: % IC Eluted in Peak 3 | 602 IC LN35 vs. 602 IC LN15 |
| One-way ANOVA (Tukey post hoc test) | IC Monomeric Purity: % IC Eluted in Peak 3 | 602 IC LN35 vs. 602 IC LN10 |
| One-way ANOVA (Tukey post hoc test) | IC Monomeric Purity: % IC Eluted in Peak 3 | 602 IC LN35 vs. F10 IC LN35 |
| One-way ANOVA (Tukey post hoc test) | IC Monomeric Purity: % IC Eluted in Peak 3 | 602 IC LN35 vs. 4-4-20 IC LN35 |
| One-way ANOVA (Tukey post hoc test) | IC Monomeric Purity: % IC Eluted in Peak 3 | 602 IC LN25 vs. 602 IC LN15 |
| One-way ANOVA (Tukey post hoc test) | IC Monomeric Purity: % IC Eluted in Peak 3 | 602 IC LN25 vs. 602 IC LN10 |
| One-way ANOVA (Tukey post hoc test) | IC Monomeric Purity: % IC Eluted in Peak 3 | 602 IC LN25 vs. F10 IC LN35 |
| One-way ANOVA (Tukey post hoc test) | IC Monomeric Purity: % IC Eluted in Peak 3 | 602 IC LN25 vs. 4-4-20 IC LN35 |
| One-way ANOVA (Tukey post hoc test) | IC Monomeric Purity: % IC Eluted in Peak 3 | 602 IC LN15 vs. 602 IC LN10 |
| One-way ANOVA (Tukey post hoc test) | IC Monomeric Purity: % IC Eluted in Peak 3 | 602 IC LN15 vs. F10 IC LN35 |
| One-way ANOVA (Tukey post hoc test) | IC Monomeric Purity: % IC Eluted in Peak 3 | 602 IC LN15 vs. 4-4-20 IC LN35 |
| One-way ANOVA (Tukey post hoc test) | IC Monomeric Purity: % IC Eluted in Peak 3 | 602 IC LN10 vs. F10 IC LN35 |
| One-way ANOVA (Tukey post hoc test) | IC Monomeric Purity: % IC Eluted in Peak 3 | 602 IC LN10 vs. 4-4-20 IC LN35 |
| One-way ANOVA (Tukey post hoc test) | IC Monomeric Purity: % IC Eluted in Peak 3 | F10 IC LN35 vs. 4-4-20 IC LN35 |
| Two-way ANOVA (Tukey post hoc test) | IL-2Rα- YT-1 Signaling | IL-2 vs. IL-2/602 Cx |
| Two-way ANOVA (Tukey post hoc test) | IL-2Rα- YT-1 Signaling | IL-2 vs. 602 IC LN15 |
| Two-way ANOVA (Tukey post hoc test) | IL-2Rα- YT-1 Signaling | IL-2 vs. 602 IC LN25 |
| Two-way ANOVA (Tukey post hoc test) | IL-2Rα- YT-1 Signaling | IL-2 vs. 602 IC LN35 |
| Two-way ANOVA (Tukey post hoc test) | IL-2Rα- YT-1 Signaling | IL-2/602 Cx vs. 602 IC LN15 |
| Two-way ANOVA (Tukey post hoc test) | IL-2Rα- YT-1 Signaling | IL-2/602 Cx vs. 602 IC LN25 |
| Two-way ANOVA (Tukey post hoc test) | IL-2Rα- YT-1 Signaling | IL-2/602 Cx vs. 602 IC LN35 |

| | | |
|---|---|---|
| Two-way ANOVA (Tukey post hoc test) | IL-2Rα- YT-1 Signaling | 602 IC LN15 vs. 602 IC LN25 |
| Two-way ANOVA (Tukey post hoc test) | IL-2Rα- YT-1 Signaling | 602 IC LN15 vs. 602 IC LN35 |
| Two-way ANOVA (Tukey post hoc test) | IL-2Rα- YT-1 Signaling | 602 IC LN25 vs. 602 IC LN35 |
| Two-way ANOVA (Tukey post hoc test) | IL-2Rα$^{+}$ YT-1 Signaling | IL-2 vs. IL-2/602 Cx |
| Two-way ANOVA (Tukey post hoc test) | IL-2Rα$^{+}$ YT-1 Signaling | IL-2 vs. 602 IC LN15 |
| Two-way ANOVA (Tukey post hoc test) | IL-2Rα$^{+}$ YT-1 Signaling | IL-2 vs. 602 IC LN25 |
| Two-way ANOVA (Tukey post hoc test) | IL-2Rα$^{+}$ YT-1 Signaling | IL-2 vs. 602 IC LN35 |
| Two-way ANOVA (Tukey post hoc test) | IL-2Rα$^{+}$ YT-1 Signaling | IL-2/602 Cx vs. 602 IC LN15 |
| Two-way ANOVA (Tukey post hoc test) | IL-2Rα$^{+}$ YT-1 Signaling | IL-2/602 Cx vs. 602 IC LN25 |
| Two-way ANOVA (Tukey post hoc test) | IL-2Rα$^{+}$ YT-1 Signaling | IL-2/602 Cx vs. 602 IC LN35 |
| Two-way ANOVA (Tukey post hoc test) | IL-2Rα$^{+}$ YT-1 Signaling | 602 IC LN15 vs. 602 IC LN25 |
| Two-way ANOVA (Tukey post hoc test) | IL-2Rα$^{+}$ YT-1 Signaling | 602 IC LN15 vs. 602 IC LN35 |
| Two-way ANOVA (Tukey post hoc test) | IL-2Rα$^{+}$ YT-1 Signaling | 602 IC LN25 vs. 602 IC LN35 |
| One-way ANOVA (Tukey post hoc test) | CD4+ Tconv:Treg Expansion Ratio | PBS vs. Control IC |
| One-way ANOVA (Tukey post hoc test) | CD4+ Tconv:Treg Expansion Ratio | PBS vs. IL-2/602 Cx |
| One-way ANOVA (Tukey post hoc test) | CD4+ Tconv:Treg Expansion Ratio | PBS vs. 602 IC |
| One-way ANOVA (Tukey post hoc test) | CD4+ Tconv:Treg Expansion Ratio | PBS vs. F10 IC |
| One-way ANOVA (Tukey post hoc test) | CD4+ Tconv:Treg Expansion Ratio | Control IC vs. IL-2/602 Cx |
| One-way ANOVA (Tukey post hoc test) | CD4+ Tconv:Treg Expansion Ratio | Control IC vs. 602 IC |
| One-way ANOVA (Tukey post hoc test) | CD4+ Tconv:Treg Expansion Ratio | Control IC vs. F10 IC |
| One-way ANOVA (Tukey post hoc test) | CD4+ Tconv:Treg Expansion Ratio | 602 IC vs. IL-2/602 Cx |
| One-way ANOVA (Tukey post hoc test) | CD4+ Tconv:Treg Expansion Ratio | F10 IC vs. IL-2/602 Cx |
| One-way ANOVA (Tukey post hoc test) | CD4+ Tconv:Treg Expansion Ratio | 602 IC vs. F10 IC |
| One-way ANOVA (Tukey post hoc test) | CD8+ T cell:Treg Expansion Ratio | PBS vs. Control IC |

| | | |
|---|---|---|
| One-way ANOVA (Tukey post hoc test) | CD8+ T cell:Treg Expansion Ratio | PBS vs. IL-2/602 Cx |
| One-way ANOVA (Tukey post hoc test) | CD8+ T cell:Treg Expansion Ratio | PBS vs. 602 IC |
| One-way ANOVA (Tukey post hoc test) | CD8+ T cell:Treg Expansion Ratio | PBS vs. F10 IC |
| One-way ANOVA (Tukey post hoc test) | CD8+ T cell:Treg Expansion Ratio | Control IC vs. IL-2/602 Cx |
| One-way ANOVA (Tukey post hoc test) | CD8+ T cell:Treg Expansion Ratio | Control IC vs. 602 IC |
| One-way ANOVA (Tukey post hoc test) | CD8+ T cell:Treg Expansion Ratio | Control IC vs. F10 IC |
| One-way ANOVA (Tukey post hoc test) | CD8+ T cell:Treg Expansion Ratio | 602 IC vs. IL-2/602 Cx |
| One-way ANOVA (Tukey post hoc test) | CD8+ T cell:Treg Expansion Ratio | F10 IC vs. IL-2/602 Cx |
| One-way ANOVA (Tukey post hoc test) | CD8+ T cell:Treg Expansion Ratio | 602 IC vs. F10 IC |
| One-way ANOVA (Tukey post hoc test) | NK:Treg Expansion Ratio | PBS vs. Control IC |
| One-way ANOVA (Tukey post hoc test) | NK:Treg Expansion Ratio | PBS vs. IL-2/602 Cx |
| One-way ANOVA (Tukey post hoc test) | NK:Treg Expansion Ratio | PBS vs. 602 IC |
| One-way ANOVA (Tukey post hoc test) | NK:Treg Expansion Ratio | PBS vs. F10 IC |
| One-way ANOVA (Tukey post hoc test) | NK:Treg Expansion Ratio | Control IC vs. IL-2/602 Cx |
| One-way ANOVA (Tukey post hoc test) | NK:Treg Expansion Ratio | Control IC vs. 602 IC |
| One-way ANOVA (Tukey post hoc test) | NK:Treg Expansion Ratio | Control IC vs. F10 IC |
| One-way ANOVA (Tukey post hoc test) | NK:Treg Expansion Ratio | 602 IC vs. IL-2/602 Cx |
| One-way ANOVA (Tukey post hoc test) | NK:Treg Expansion Ratio | F10 IC vs. IL-2/602 Cx |
| One-way ANOVA (Tukey post hoc test) | NK:Treg Expansion Ratio | 602 IC vs. F10 IC |
| Two-way ANOVA (Tukey post hoc test) | B16F10 Melanoma Tumor Volume | PBS vs. Control IC |
| Two-way ANOVA (Tukey post hoc test) | B16F10 Melanoma Tumor Volume | PBS vs. F10 IC |
| Two-way ANOVA (Tukey post hoc test) | B16F10 Melanoma Tumor Volume | Control IC vs. F10 IC |
| Two-way ANOVA (Tukey post hoc test) | CT26 Colon Carcinoma Tumor Volume | PBS vs. Control IC |
| Two-way ANOVA (Tukey post hoc test) | CT26 Colon Carcinoma Tumor Volume | PBS vs. F10 IC |

| | | |
|---|---|---|
| Two-way ANOVA (Tukey post hoc test) | CT26 Colon Carcinoma Tumor Volume | Control IC vs. F10 IC |
| Two-way ANOVA (Tukey post hoc test) | B16F10 Melanoma Tumor Volume | PBS vs. Control IC |
| Two-way ANOVA (Tukey post hoc test) | B16F10 Melanoma Tumor Volume | PBS vs. F10 IC |
| Two-way ANOVA (Tukey post hoc test) | B16F10 Melanoma Tumor Volume | Control IC vs. F10 IC |
| Two-way ANOVA (Tukey post hoc test) | CT26 Colon Carcinoma Percent Weight Change | PBS vs. Control IC |
| Two-way ANOVA (Tukey post hoc test) | CT26 Colon Carcinoma Percent Weight Change | PBS vs. F10 IC |
| Two-way ANOVA (Tukey post hoc test) | CT26 Colon Carcinoma Percent Weight Change | Control IC vs. F10 IC |
| One-way ANOVA (Tukey post hoc test) | C57BL/6 Pulmonary Wet Weight | PBS vs. 602 IC |
| One-way ANOVA (Tukey post hoc test) | C57BL/6 Pulmonary Wet Weight | PBS vs. Control IC |
| One-way ANOVA (Tukey post hoc test) | C57BL/6 Pulmonary Wet Weight | PBS vs. F10 IC |
| One-way ANOVA (Tukey post hoc test) | C57BL/6 Pulmonary Wet Weight | PBS vs. IL-2/602 Cx |
| One-way ANOVA (Tukey post hoc test) | C57BL/6 Pulmonary Wet Weight | PBS vs. IL-2 |
| One-way ANOVA (Tukey post hoc test) | C57BL/6 Pulmonary Wet Weight | 602 IC vs. Control IC |
| One-way ANOVA (Tukey post hoc test) | C57BL/6 Pulmonary Wet Weight | 602 IC vs. F10 IC |
| One-way ANOVA (Tukey post hoc test) | C57BL/6 Pulmonary Wet Weight | 602 IC vs. IL-2/602 Cx |
| One-way ANOVA (Tukey post hoc test) | C57BL/6 Pulmonary Wet Weight | 602 IC vs. IL-2 |
| One-way ANOVA (Tukey post hoc test) | C57BL/6 Pulmonary Wet Weight | Control IC vs. F10 IC |
| One-way ANOVA (Tukey post hoc test) | C57BL/6 Pulmonary Wet Weight | Control IC vs. IL-2/602 Cx |
| One-way ANOVA (Tukey post hoc test) | C57BL/6 Pulmonary Wet Weight | Control IC vs. IL-2 |
| One-way ANOVA (Tukey post hoc test) | C57BL/6 Pulmonary Wet Weight | F10 IC vs. IL-2/602 Cx |
| One-way ANOVA (Tukey post hoc test) | C57BL/6 Pulmonary Wet Weight | F10 IC vs. IL-2 |
| One-way ANOVA (Tukey post hoc test) | C57BL/6 Pulmonary Wet Weight | IL-2/602 Cx vs. IL-2 |
| One-way ANOVA (Tukey post hoc test) | IC Multimeric Purity: % IC Eluted in Peak 1 | 602 IC LN35 vs. 602 IC LN25 |
| One-way ANOVA (Tukey post hoc test) | IC Multimeric Purity: % IC Eluted in Peak 1 | 602 IC LN35 vs. 602 IC LN15 |

| | | |
|---|---|---|
| One-way ANOVA (Tukey post hoc test) | IC Multimeric Purity: % IC Eluted in Peak 1 | 602 IC LN35 vs. 602 IC LN10 |
| One-way ANOVA (Tukey post hoc test) | IC Multimeric Purity: % IC Eluted in Peak 1 | 602 IC LN35 vs. F10 IC LN35 |
| One-way ANOVA (Tukey post hoc test) | IC Multimeric Purity: % IC Eluted in Peak 1 | 602 IC LN35 vs. 4-4-20 IC LN35 |
| One-way ANOVA (Tukey post hoc test) | IC Multimeric Purity: % IC Eluted in Peak 1 | 602 IC LN25 vs. 602 IC LN15 |
| One-way ANOVA (Tukey post hoc test) | IC Multimeric Purity: % IC Eluted in Peak 1 | 602 IC LN25 vs. 602 IC LN10 |
| One-way ANOVA (Tukey post hoc test) | IC Multimeric Purity: % IC Eluted in Peak 1 | 602 IC LN25 vs. F10 IC LN35 |
| One-way ANOVA (Tukey post hoc test) | IC Multimeric Purity: % IC Eluted in Peak 1 | 602 IC LN25 vs. 4-4-20 IC LN35 |
| One-way ANOVA (Tukey post hoc test) | IC Multimeric Purity: % IC Eluted in Peak 1 | 602 IC LN15 vs. 602 IC LN10 |
| One-way ANOVA (Tukey post hoc test) | IC Multimeric Purity: % IC Eluted in Peak 1 | 602 IC LN15 vs. F10 IC LN35 |
| One-way ANOVA (Tukey post hoc test) | IC Multimeric Purity: % IC Eluted in Peak 1 | 602 IC LN15 vs. 4-4-20 IC LN35 |
| One-way ANOVA (Tukey post hoc test) | IC Multimeric Purity: % IC Eluted in Peak 1 | 602 IC LN10 vs. F10 IC LN35 |
| One-way ANOVA (Tukey post hoc test) | IC Multimeric Purity: % IC Eluted in Peak 1 | 602 IC LN10 vs. 4-4-20 IC LN35 |
| One-way ANOVA (Tukey post hoc test) | IC Multimeric Purity: % IC Eluted in Peak 1 | F10 IC LN35 vs. 4-4-20 IC LN35 |
| One-way ANOVA (Tukey post hoc test) | IC Multimeric Purity: % IC Eluted in Peak 2 | 602 IC LN35 vs. 602 IC LN25 |
| One-way ANOVA (Tukey post hoc test) | IC Multimeric Purity: % IC Eluted in Peak 2 | 602 IC LN35 vs. 602 IC LN15 |
| One-way ANOVA (Tukey post hoc test) | IC Multimeric Purity: % IC Eluted in Peak 2 | 602 IC LN35 vs. 602 IC LN10 |
| One-way ANOVA (Tukey post hoc test) | IC Multimeric Purity: % IC Eluted in Peak 2 | 602 IC LN35 vs. F10 IC LN35 |
| One-way ANOVA (Tukey post hoc test) | IC Multimeric Purity: % IC Eluted in Peak 2 | 602 IC LN35 vs. 4-4-20 IC LN35 |
| One-way ANOVA (Tukey post hoc test) | IC Multimeric Purity: % IC Eluted in Peak 2 | 602 IC LN25 vs. 602 IC LN15 |
| One-way ANOVA (Tukey post hoc test) | IC Multimeric Purity: % IC Eluted in Peak 2 | 602 IC LN25 vs. 602 IC LN10 |
| One-way ANOVA (Tukey post hoc test) | IC Multimeric Purity: % IC Eluted in Peak 2 | 602 IC LN25 vs. F10 IC LN35 |
| One-way ANOVA (Tukey post hoc test) | IC Multimeric Purity: % IC Eluted in Peak 2 | 602 IC LN25 vs. 4-4-20 IC LN35 |
| One-way ANOVA (Tukey post hoc test) | IC Multimeric Purity: % IC Eluted in Peak 2 | 602 IC LN15 vs. 602 IC LN10 |
| One-way ANOVA (Tukey post hoc test) | IC Multimeric Purity: % IC Eluted in Peak 2 | 602 IC LN15 vs. F10 IC LN35 |

| | | |
|---|---|---|
| One-way ANOVA (Tukey post hoc test) | IC Multimeric Purity: % IC Eluted in Peak 2 | 602 IC LN15 vs. 4-4-20 IC LN35 |
| One-way ANOVA (Tukey post hoc test) | IC Multimeric Purity: % IC Eluted in Peak 2 | 602 IC LN10 vs. F10 IC LN35 |
| One-way ANOVA (Tukey post hoc test) | IC Multimeric Purity: % IC Eluted in Peak 2 | 602 IC LN10 vs. 4-4-20 IC LN35 |
| One-way ANOVA (Tukey post hoc test) | IC Multimeric Purity: % IC Eluted in Peak 2 | F10 IC LN35 vs. 4-4-20 IC LN35 |
| Two-way ANOVA (Tukey post hoc test) | IL-2 Binding on Yeast: IL-2Rα Competition | 602 scFv vs. A8 scFv |
| Two-way ANOVA (Tukey post hoc test) | IL-2 Binding on Yeast: IL-2Rα Competition | 602 scFv vs. C5 scFv |
| Two-way ANOVA (Tukey post hoc test) | IL-2 Binding on Yeast: IL-2Rα Competition | 602 scFv vs. E10 scFv |
| Two-way ANOVA (Tukey post hoc test) | IL-2 Binding on Yeast: IL-2Rα Competition | 602 scFv vs. F10 scFv |
| Two-way ANOVA (Tukey post hoc test) | IL-2 Binding on Yeast: IL-2Rα Competition | 602 scFv vs. H2 scFv |
| Two-way ANOVA (Tukey post hoc test) | IL-2 Binding on Yeast: IL-2Rα Competition | A8 scFv vs. C5 scFv |
| Two-way ANOVA (Tukey post hoc test) | IL-2 Binding on Yeast: IL-2Rα Competition | A8 scFv vs. E10 scFv |
| Two-way ANOVA (Tukey post hoc test) | IL-2 Binding on Yeast: IL-2Rα Competition | A8 scFv vs. F10 scFv |
| Two-way ANOVA (Tukey post hoc test) | IL-2 Binding on Yeast: IL-2Rα Competition | A8 scFv vs. H2 scFv |
| Two-way ANOVA (Tukey post hoc test) | IL-2 Binding on Yeast: IL-2Rα Competition | C5 scFv vs. E10 scFv |
| Two-way ANOVA (Tukey post hoc test) | IL-2 Binding on Yeast: IL-2Rα Competition | C5 scFv vs. F10 scFv |
| Two-way ANOVA (Tukey post hoc test) | IL-2 Binding on Yeast: IL-2Rα Competition | C5 scFv vs. H2 scFv |
| Two-way ANOVA (Tukey post hoc test) | IL-2 Binding on Yeast: IL-2Rα Competition | E10 scFv vs. F10 scFv |
| Two-way ANOVA (Tukey post hoc test) | IL-2 Binding on Yeast: IL-2Rα Competition | E10 scFv vs. H2 scFv |
| Two-way ANOVA (Tukey post hoc test) | IL-2 Binding on Yeast: IL-2Rα Competition | F10 scFv vs. H2 scFv |
| One-way ANOVA (Tukey post hoc test) | CD4+ Tconv Fold Expansion Over PBS | IL-2/602 Cx vs. Control IC |
| One-way ANOVA (Tukey post hoc test) | CD4+ Tconv Fold Expansion Over PBS | IL-2/602 Cx vs. 602 IC |
| One-way ANOVA (Tukey post hoc test) | CD4+ Tconv Fold Expansion Over PBS | IL-2/602 Cx vs. F10 IC |
| One-way ANOVA (Tukey post hoc test) | CD4+ Tconv Fold Expansion Over PBS | Control IC vs. 602 IC |
| One-way ANOVA (Tukey post hoc test) | CD4+ Tconv Fold Expansion Over PBS | Control IC vs. F10 IC |

| | | |
|---|---|---|
| One-way ANOVA (Tukey post hoc test) | CD4+ Tconv Fold Expansion Over PBS | 602 IC vs. F10 IC |
| One-way ANOVA (Tukey post hoc test) | CD8+ T cell Fold Expansion Over PBS | IL-2/602 Cx vs. Control IC |
| One-way ANOVA (Tukey post hoc test) | CD8+ T cell Fold Expansion Over PBS | IL-2/602 Cx vs. 602 IC |
| One-way ANOVA (Tukey post hoc test) | CD8+ T cell Fold Expansion Over PBS | IL-2/602 Cx vs. F10 IC |
| One-way ANOVA (Tukey post hoc test) | CD8+ T cell Fold Expansion Over PBS | Control IC vs. 602 IC |
| One-way ANOVA (Tukey post hoc test) | CD8+ T cell Fold Expansion Over PBS | Control IC vs. F10 IC |
| One-way ANOVA (Tukey post hoc test) | CD8+ T cell Fold Expansion Over PBS | 602 IC vs. F10 IC |
| One-way ANOVA (Tukey post hoc test) | NK Fold Expansion Over PBS | IL-2/602 Cx vs. Control IC |
| One-way ANOVA (Tukey post hoc test) | NK Fold Expansion Over PBS | IL-2/602 Cx vs. 602 IC |
| One-way ANOVA (Tukey post hoc test) | NK Fold Expansion Over PBS | IL-2/602 Cx vs. F10 IC |
| One-way ANOVA (Tukey post hoc test) | NK Fold Expansion Over PBS | Control IC vs. 602 IC |
| One-way ANOVA (Tukey post hoc test) | NK Fold Expansion Over PBS | Control IC vs. F10 IC |
| One-way ANOVA (Tukey post hoc test) | NK Fold Expansion Over PBS | 602 IC vs. F10 IC |
| One-way ANOVA (Tukey post hoc test) | CD4+ T-EM Fold Expansion Over PBS | IL-2/602 Cx vs. Control IC |
| One-way ANOVA (Tukey post hoc test) | CD4+ T-EM Fold Expansion Over PBS | IL-2/602 Cx vs. 602 IC |
| One-way ANOVA (Tukey post hoc test) | CD4+ T-EM Fold Expansion Over PBS | IL-2/602 Cx vs. F10 IC |
| One-way ANOVA (Tukey post hoc test) | CD4+ T-EM Fold Expansion Over PBS | Control IC vs. 602 IC |
| One-way ANOVA (Tukey post hoc test) | CD4+ T-EM Fold Expansion Over PBS | Control IC vs. F10 IC |
| One-way ANOVA (Tukey post hoc test) | CD4+ T-EM Fold Expansion Over PBS | 602 IC vs. F10 IC |
| One-way ANOVA (Tukey post hoc test) | CD8+ T-MP Fold Expansion Over PBS | IL-2/602 Cx vs. Control IC |
| One-way ANOVA (Tukey post hoc test) | CD8+ T-MP Fold Expansion Over PBS | IL-2/602 Cx vs. 602 IC |
| One-way ANOVA (Tukey post hoc test) | CD8+ T-MP Fold Expansion Over PBS | IL-2/602 Cx vs. F10 IC |
| One-way ANOVA (Tukey post hoc test) | CD8+ T-MP Fold Expansion Over PBS | Control IC vs. 602 IC |
| One-way ANOVA (Tukey post hoc test) | CD8+ T-MP Fold Expansion Over PBS | Control IC vs. F10 IC |

| | | |
|---|---|---|
| One-way ANOVA (Tukey post hoc test) | CD8+ T-MP Fold Expansion Over PBS | 602 IC vs. F10 IC |
| One-way ANOVA (Tukey post hoc test) | Treg Fold Expansion Over PBS | IL-2/602 Cx vs. Control IC |
| One-way ANOVA (Tukey post hoc test) | Treg Fold Expansion Over PBS | IL-2/602 Cx vs. 602 IC |
| One-way ANOVA (Tukey post hoc test) | Treg Fold Expansion Over PBS | IL-2/602 Cx vs. F10 IC |
| One-way ANOVA (Tukey post hoc test) | Treg Fold Expansion Over PBS | Control IC vs. 602 IC |
| One-way ANOVA (Tukey post hoc test) | Treg Fold Expansion Over PBS | Control IC vs. F10 IC |
| One-way ANOVA (Tukey post hoc test) | Treg Fold Expansion Over PBS | 602 IC vs. F10 IC |
| One-way ANOVA (Tukey post hoc test) | CD4+ Tconv Fold Expansion Over PBS | Control IC vs. Control IC ΔFc |
| One-way ANOVA (Tukey post hoc test) | CD4+ Tconv Fold Expansion Over PBS | 602 IC vs. 602 IC ΔFc |
| One-way ANOVA (Tukey post hoc test) | CD4+ Tconv Fold Expansion Over PBS | F10 IC vs. F10 IC ΔFc |
| One-way ANOVA (Tukey post hoc test) | CD8+ T cell Fold Expansion Over PBS | Control IC vs. Control IC ΔFc |
| One-way ANOVA (Tukey post hoc test) | CD8+ T cell Fold Expansion Over PBS | 602 IC vs. 602 IC ΔFc |
| One-way ANOVA (Tukey post hoc test) | CD8+ T cell Fold Expansion Over PBS | F10 IC vs. F10 IC ΔFc |
| One-way ANOVA (Tukey post hoc test) | NK Fold Expansion Over PBS | Control IC vs. Control IC ΔFc |
| One-way ANOVA (Tukey post hoc test) | NK Fold Expansion Over PBS | 602 IC vs. 602 IC ΔFc |
| One-way ANOVA (Tukey post hoc test) | NK Fold Expansion Over PBS | F10 IC vs. F10 IC ΔFc |
| One-way ANOVA (Tukey post hoc test) | CD4+ T-EM Fold Expansion Over PBS | Control IC vs. Control IC ΔFc |
| One-way ANOVA (Tukey post hoc test) | CD4+ T-EM Fold Expansion Over PBS | 602 IC vs. 602 IC ΔFc |
| One-way ANOVA (Tukey post hoc test) | CD4+ T-EM Fold Expansion Over PBS | F10 IC vs. F10 IC ΔFc |
| One-way ANOVA (Tukey post hoc test) | CD8+ T-MP Fold Expansion Over PBS | Control IC vs. Control IC ΔFc |
| One-way ANOVA (Tukey post hoc test) | CD8+ T-MP Fold Expansion Over PBS | 602 IC vs. 602 IC ΔFc |
| One-way ANOVA (Tukey post hoc test) | CD8+ T-MP Fold Expansion Over PBS | F10 IC vs. F10 IC ΔFc |
| One-way ANOVA (Tukey post hoc test) | Treg Fold Expansion Over PBS | Control IC vs. Control IC ΔFc |
| One-way ANOVA (Tukey post hoc test) | Treg Fold Expansion Over PBS | 602 IC vs. 602 IC ΔFc |

| | | |
|---|---|---|
| One-way ANOVA (Tukey post hoc test) | Treg Fold Expansion Over PBS | F10 IC vs. F10 IC ΔFc |
| One-way ANOVA (Tukey post hoc test) | CD4+ Tconv Fold Expansion Over PBS | Control IC vs. 602 IC ΔFc |
| One-way ANOVA (Tukey post hoc test) | CD4+ Tconv Fold Expansion Over PBS | Control IC vs. F10 IC ΔFc |
| One-way ANOVA (Tukey post hoc test) | CD4+ Tconv Fold Expansion Over PBS | Control IC ΔFc vs. 602 IC |
| One-way ANOVA (Tukey post hoc test) | CD4+ Tconv Fold Expansion Over PBS | Control IC ΔFc vs. F10 IC |
| One-way ANOVA (Tukey post hoc test) | CD4+ Tconv Fold Expansion Over PBS | 602 IC vs. F10 IC ΔFc |
| One-way ANOVA (Tukey post hoc test) | CD4+ Tconv Fold Expansion Over PBS | F10 IC vs. 602 IC ΔFc |
| One-way ANOVA (Tukey post hoc test) | CD8+ T cell Fold Expansion Over PBS | Control IC vs. 602 IC ΔFc |
| One-way ANOVA (Tukey post hoc test) | CD8+ T cell Fold Expansion Over PBS | Control IC vs. F10 IC ΔFc |
| One-way ANOVA (Tukey post hoc test) | CD8+ T cell Fold Expansion Over PBS | Control IC ΔFc vs. 602 IC |
| One-way ANOVA (Tukey post hoc test) | CD8+ T cell Fold Expansion Over PBS | Control IC ΔFc vs. F10 IC |
| One-way ANOVA (Tukey post hoc test) | CD8+ T cell Fold Expansion Over PBS | 602 IC vs. F10 IC ΔFc |
| One-way ANOVA (Tukey post hoc test) | CD8+ T cell Fold Expansion Over PBS | F10 IC vs. 602 IC ΔFc |
| One-way ANOVA (Tukey post hoc test) | NK Fold Expansion Over PBS | Control IC vs. 602 IC ΔFc |
| One-way ANOVA (Tukey post hoc test) | NK Fold Expansion Over PBS | Control IC vs. F10 IC ΔFc |
| One-way ANOVA (Tukey post hoc test) | NK Fold Expansion Over PBS | Control IC ΔFc vs. 602 IC |
| One-way ANOVA (Tukey post hoc test) | NK Fold Expansion Over PBS | Control IC ΔFc vs. F10 IC |
| One-way ANOVA (Tukey post hoc test) | NK Fold Expansion Over PBS | 602 IC vs. F10 IC ΔFc |
| One-way ANOVA (Tukey post hoc test) | NK Fold Expansion Over PBS | F10 IC vs. 602 IC ΔFc |
| One-way ANOVA (Tukey post hoc test) | CD4+ T-EM Fold Expansion Over PBS | Control IC vs. 602 IC ΔFc |
| One-way ANOVA (Tukey post hoc test) | CD4+ T-EM Fold Expansion Over PBS | Control IC vs. F10 IC ΔFc |
| One-way ANOVA (Tukey post hoc test) | CD4+ T-EM Fold Expansion Over PBS | Control IC ΔFc vs. 602 IC |
| One-way ANOVA (Tukey post hoc test) | CD4+ T-EM Fold Expansion Over PBS | Control IC ΔFc vs. F10 IC |
| One-way ANOVA (Tukey post hoc test) | CD4+ T-EM Fold Expansion Over PBS | 602 IC vs. F10 IC ΔFc |

| | | |
|---|---|---|
| One-way ANOVA (Tukey post hoc test) | CD4+ T-EM Fold Expansion Over PBS | F10 IC vs. 602 IC ΔFc |
| One-way ANOVA (Tukey post hoc test) | CD8+ T-MP Fold Expansion Over PBS | Control IC vs. 602 IC ΔFc |
| One-way ANOVA (Tukey post hoc test) | CD8+ T-MP Fold Expansion Over PBS | Control IC vs. F10 IC ΔFc |
| One-way ANOVA (Tukey post hoc test) | CD8+ T-MP Fold Expansion Over PBS | Control IC ΔFc vs. 602 IC |
| One-way ANOVA (Tukey post hoc test) | CD8+ T-MP Fold Expansion Over PBS | Control IC ΔFc vs. F10 IC |
| One-way ANOVA (Tukey post hoc test) | CD8+ T-MP Fold Expansion Over PBS | 602 IC vs. F10 IC ΔFc |
| One-way ANOVA (Tukey post hoc test) | CD8+ T-MP Fold Expansion Over PBS | F10 IC vs. 602 IC ΔFc |
| One-way ANOVA (Tukey post hoc test) | Treg Fold Expansion Over PBS | Control IC vs. 602 IC ΔFc |
| One-way ANOVA (Tukey post hoc test) | Treg Fold Expansion Over PBS | Control IC vs. F10 IC ΔFc |
| One-way ANOVA (Tukey post hoc test) | Treg Fold Expansion Over PBS | Control IC ΔFc vs. 602 IC |
| One-way ANOVA (Tukey post hoc test) | Treg Fold Expansion Over PBS | Control IC ΔFc vs. F10 IC |
| One-way ANOVA (Tukey post hoc test) | Treg Fold Expansion Over PBS | 602 IC vs. F10 IC ΔFc |
| One-way ANOVA (Tukey post hoc test) | Treg Fold Expansion Over PBS | F10 IC vs. 602 IC ΔFc |
| One-way ANOVA (Tukey post hoc test) | CD4+ Tconv Fold Expansion Over PBS | IL-2/602 Cx vs. Control IC ΔFc |
| One-way ANOVA (Tukey post hoc test) | CD4+ Tconv Fold Expansion Over PBS | IL-2/602 Cx vs. 602 IC ΔFc |
| One-way ANOVA (Tukey post hoc test) | CD4+ Tconv Fold Expansion Over PBS | IL-2/602 Cx vs. F10 IC ΔFc |
| One-way ANOVA (Tukey post hoc test) | CD4+ Tconv Fold Expansion Over PBS | Control IC ΔFc vs. 602 IC ΔFc |
| One-way ANOVA (Tukey post hoc test) | CD4+ Tconv Fold Expansion Over PBS | Control IC ΔFc vs. F10 IC ΔFc |
| One-way ANOVA (Tukey post hoc test) | CD4+ Tconv Fold Expansion Over PBS | 602 IC ΔFc vs. F10 IC ΔFc |
| One-way ANOVA (Tukey post hoc test) | CD8+ T cell Fold Expansion Over PBS | IL-2/602 Cx vs. Control IC ΔFc |
| One-way ANOVA (Tukey post hoc test) | CD8+ T cell Fold Expansion Over PBS | IL-2/602 Cx vs. 602 IC ΔFc |
| One-way ANOVA (Tukey post hoc test) | CD8+ T cell Fold Expansion Over PBS | IL-2/602 Cx vs. F10 IC ΔFc |
| One-way ANOVA (Tukey post hoc test) | CD8+ T cell Fold Expansion Over PBS | Control IC ΔFc vs. 602 IC ΔFc |
| One-way ANOVA (Tukey post hoc test) | CD8+ T cell Fold Expansion Over PBS | Control IC ΔFc vs. F10 IC ΔFc |

| | | |
|---|---|---|
| One-way ANOVA (Tukey post hoc test) | CD8+ T cell Fold Expansion Over PBS | 602 IC ΔFc vs. F10 IC ΔFc |
| One-way ANOVA (Tukey post hoc test) | NK Fold Expansion Over PBS | IL-2/602 Cx vs. Control IC ΔFc |
| One-way ANOVA (Tukey post hoc test) | NK Fold Expansion Over PBS | IL-2/602 Cx vs. 602 IC ΔFc |
| One-way ANOVA (Tukey post hoc test) | NK Fold Expansion Over PBS | IL-2/602 Cx vs. F10 IC ΔFc |
| One-way ANOVA (Tukey post hoc test) | NK Fold Expansion Over PBS | Control IC ΔFc vs. 602 IC ΔFc |
| One-way ANOVA (Tukey post hoc test) | NK Fold Expansion Over PBS | Control IC ΔFc vs. F10 IC ΔFc |
| One-way ANOVA (Tukey post hoc test) | NK Fold Expansion Over PBS | 602 IC ΔFc vs. F10 IC ΔFc |
| One-way ANOVA (Tukey post hoc test) | CD4+ T-EM Fold Expansion Over PBS | IL-2/602 Cx vs. Control IC ΔFc |
| One-way ANOVA (Tukey post hoc test) | CD4+ T-EM Fold Expansion Over PBS | IL-2/602 Cx vs. 602 IC ΔFc |
| One-way ANOVA (Tukey post hoc test) | CD4+ T-EM Fold Expansion Over PBS | IL-2/602 Cx vs. F10 IC ΔFc |
| One-way ANOVA (Tukey post hoc test) | CD4+ T-EM Fold Expansion Over PBS | Control IC ΔFc vs. 602 IC ΔFc |
| One-way ANOVA (Tukey post hoc test) | CD4+ T-EM Fold Expansion Over PBS | Control IC ΔFc vs. F10 IC ΔFc |
| One-way ANOVA (Tukey post hoc test) | CD4+ T-EM Fold Expansion Over PBS | 602 IC ΔFc vs. F10 IC ΔFc |
| One-way ANOVA (Tukey post hoc test) | CD8+ T-MP Fold Expansion Over PBS | IL-2/602 Cx vs. Control IC ΔFc |
| One-way ANOVA (Tukey post hoc test) | CD8+ T-MP Fold Expansion Over PBS | IL-2/602 Cx vs. 602 IC ΔFc |
| One-way ANOVA (Tukey post hoc test) | CD8+ T-MP Fold Expansion Over PBS | IL-2/602 Cx vs. F10 IC ΔFc |
| One-way ANOVA (Tukey post hoc test) | CD8+ T-MP Fold Expansion Over PBS | Control IC ΔFc vs. 602 IC ΔFc |
| One-way ANOVA (Tukey post hoc test) | CD8+ T-MP Fold Expansion Over PBS | Control IC ΔFc vs. F10 IC ΔFc |
| One-way ANOVA (Tukey post hoc test) | CD8+ T-MP Fold Expansion Over PBS | 602 IC ΔFc vs. F10 IC ΔFc |
| One-way ANOVA (Tukey post hoc test) | Treg Fold Expansion Over PBS | IL-2/602 Cx vs. Control IC ΔFc |
| One-way ANOVA (Tukey post hoc test) | Treg Fold Expansion Over PBS | IL-2/602 Cx vs. 602 IC ΔFc |
| One-way ANOVA (Tukey post hoc test) | Treg Fold Expansion Over PBS | IL-2/602 Cx vs. F10 IC ΔFc |
| One-way ANOVA (Tukey post hoc test) | Treg Fold Expansion Over PBS | Control IC ΔFc vs. 602 IC ΔFc |
| One-way ANOVA (Tukey post hoc test) | Treg Fold Expansion Over PBS | Control IC ΔFc vs. F10 IC ΔFc |

| | | |
|---|---|---|
| One-way ANOVA (Tukey post hoc test) | Treg Fold Expansion Over PBS | 602 IC ΔFc vs. F10 IC ΔFc |
| One-way ANOVA (Tukey post hoc test) | CD4+ T-EM:Treg Expansion Ratio | PBS vs. Control IC |
| One-way ANOVA (Tukey post hoc test) | CD4+ T-EM:Treg Expansion Ratio | PBS vs. IL-2/602 Cx |
| One-way ANOVA (Tukey post hoc test) | CD4+ T-EM:Treg Expansion Ratio | PBS vs. 602 IC |
| One-way ANOVA (Tukey post hoc test) | CD4+ T-EM:Treg Expansion Ratio | PBS vs. F10 IC |
| One-way ANOVA (Tukey post hoc test) | CD4+ T-EM:Treg Expansion Ratio | Control IC vs. IL-2/602 Cx |
| One-way ANOVA (Tukey post hoc test) | CD4+ T-EM:Treg Expansion Ratio | Control IC vs. 602 IC |
| One-way ANOVA (Tukey post hoc test) | CD4+ T-EM:Treg Expansion Ratio | Control IC vs. F10 IC |
| One-way ANOVA (Tukey post hoc test) | CD4+ T-EM:Treg Expansion Ratio | 602 IC vs. IL-2/602 Cx |
| One-way ANOVA (Tukey post hoc test) | CD4+ T-EM:Treg Expansion Ratio | F10 IC vs. IL-2/602 Cx |
| One-way ANOVA (Tukey post hoc test) | CD4+ T-EM:Treg Expansion Ratio | 602 IC vs. F10 IC |
| One-way ANOVA (Tukey post hoc test) | CD8+ T-MP:Treg Expansion Ratio | PBS vs. Control IC |
| One-way ANOVA (Tukey post hoc test) | CD8+ T-MP:Treg Expansion Ratio | PBS vs. IL-2/602 Cx |
| One-way ANOVA (Tukey post hoc test) | CD8+ T-MP:Treg Expansion Ratio | PBS vs. 602 IC |
| One-way ANOVA (Tukey post hoc test) | CD8+ T-MP:Treg Expansion Ratio | PBS vs. F10 IC |
| One-way ANOVA (Tukey post hoc test) | CD8+ T-MP:Treg Expansion Ratio | Control IC vs. IL-2/602 Cx |
| One-way ANOVA (Tukey post hoc test) | CD8+ T-MP:Treg Expansion Ratio | Control IC vs. 602 IC |
| One-way ANOVA (Tukey post hoc test) | CD8+ T-MP:Treg Expansion Ratio | Control IC vs. F10 IC |
| One-way ANOVA (Tukey post hoc test) | CD8+ T-MP:Treg Expansion Ratio | 602 IC vs. IL-2/602 Cx |
| One-way ANOVA (Tukey post hoc test) | CD8+ T-MP:Treg Expansion Ratio | F10 IC vs. IL-2/602 Cx |
| One-way ANOVA (Tukey post hoc test) | CD8+ T-MP:Treg Expansion Ratio | 602 IC vs. F10 IC |
| Two-way ANOVA (Tukey post hoc test) | B16F10 Melanoma Tumor Volume | IL-2 vs. Control |
| Two-way ANOVA (Tukey post hoc test) | B16F10 Melanoma Tumor Volume | IL-2 vs. 602 Cx |
| Two-way ANOVA (Tukey post hoc test) | B16F10 Melanoma Tumor Volume | IL-2 vs. 602 IC |

| | | |
|---|---|---|
| Two-way ANOVA (Tukey post hoc test) | B16F10 Melanoma Tumor Volume | IL-2 vs. F10 IC |
| Two-way ANOVA (Tukey post hoc test) | B16F10 Melanoma Tumor Volume | Control vs. 602 Cx |
| Two-way ANOVA (Tukey post hoc test) | B16F10 Melanoma Tumor Volume | Control vs. 602 IC |
| Two-way ANOVA (Tukey post hoc test) | B16F10 Melanoma Tumor Volume | Control vs. F10 IC |
| Two-way ANOVA (Tukey post hoc test) | B16F10 Melanoma Tumor Volume | 602 Cx vs. 602 IC |
| Two-way ANOVA (Tukey post hoc test) | B16F10 Melanoma Tumor Volume | 602 Cx vs. F10 IC |
| Two-way ANOVA (Tukey post hoc test) | B16F10 Melanoma Tumor Volume | 602 IC vs. F10 IC |
| One-way ANOVA (Tukey post hoc test) | BALB/c AST | PBS vs. 602 IC |
| One-way ANOVA (Tukey post hoc test) | BALB/c AST | PBS vs. Control IC |
| One-way ANOVA (Tukey post hoc test) | BALB/c AST | PBS vs. F10 IC |
| One-way ANOVA (Tukey post hoc test) | BALB/c AST | PBS vs. IL-2/602 Cx |
| One-way ANOVA (Tukey post hoc test) | BALB/c AST | 602 IC vs. Control IC |
| One-way ANOVA (Tukey post hoc test) | BALB/c AST | 602 IC vs. F10 IC |
| One-way ANOVA (Tukey post hoc test) | BALB/c AST | 602 IC vs. IL-2/602 Cx |
| One-way ANOVA (Tukey post hoc test) | BALB/c AST | Control IC vs. F10 IC |
| One-way ANOVA (Tukey post hoc test) | BALB/c AST | Control IC vs. IL-2/602 Cx |
| One-way ANOVA (Tukey post hoc test) | BALB/c AST | F10 IC vs. IL-2/602 Cx |
| One-way ANOVA (Tukey post hoc test) | BALB/c ALT | PBS vs. 602 IC |
| One-way ANOVA (Tukey post hoc test) | BALB/c ALT | PBS vs. Control IC |
| One-way ANOVA (Tukey post hoc test) | BALB/c ALT | PBS vs. F10 IC |
| One-way ANOVA (Tukey post hoc test) | BALB/c ALT | PBS vs. IL-2/602 Cx |
| One-way ANOVA (Tukey post hoc test) | BALB/c ALT | 602 IC vs. Control IC |
| One-way ANOVA (Tukey post hoc test) | BALB/c ALT | 602 IC vs. F10 IC |
| One-way ANOVA (Tukey post hoc test) | BALB/c ALT | 602 IC vs. IL-2/602 Cx |

| | | |
|---|---|---|
| One-way ANOVA (Tukey post hoc test) | BALB/c ALT | Control IC vs. F10 IC |
| One-way ANOVA (Tukey post hoc test) | BALB/c ALT | Control IC vs. IL-2/602 Cx |
| One-way ANOVA (Tukey post hoc test) | BALB/c ALT | F10 IC vs. IL-2/602 Cx |
| One-way ANOVA (Tukey post hoc test) | BALB/c Pulmonary Wet Weight | PBS vs. 602 IC |
| One-way ANOVA (Tukey post hoc test) | BALB/c Pulmonary Wet Weight | PBS vs. Control IC |
| One-way ANOVA (Tukey post hoc test) | BALB/c Pulmonary Wet Weight | PBS vs. F10 IC |
| One-way ANOVA (Tukey post hoc test) | BALB/c Pulmonary Wet Weight | PBS vs. IL-2/602 Cx |
| One-way ANOVA (Tukey post hoc test) | BALB/c Pulmonary Wet Weight | 602 IC vs. Control IC |
| One-way ANOVA (Tukey post hoc test) | BALB/c Pulmonary Wet Weight | 602 IC vs. F10 IC |
| One-way ANOVA (Tukey post hoc test) | BALB/c Pulmonary Wet Weight | 602 IC vs. IL-2/602 Cx |
| One-way ANOVA (Tukey post hoc test) | BALB/c Pulmonary Wet Weight | Control IC vs. F10 IC |
| One-way ANOVA (Tukey post hoc test) | BALB/c Pulmonary Wet Weight | Control IC vs. IL-2/602 Cx |
| One-way ANOVA (Tukey post hoc test) | BALB/c Pulmonary Wet Weight | F10 IC vs. IL-2/602 Cx |

**ysis**

| Summary | Adjusted P Value | Figure |
|---|---|---|
| **** | <0.0001 | 1D |
| **** | <0.0001 | 1D |
| **** | <0.0001 | 1D |
| ns | 0.0983 | 1D |
| ns | 0.4958 | 1D |
| ns | 0.0976 | 1D |
| ** | 0.0087 | 1D |
| ** | 0.0015 | 1D |
| **** | <0.0001 | 1D |
| ns | 0.3867 | 1D |
| **** | <0.0001 | 1D |
| **** | <0.0001 | 1D |
| **** | <0.0001 | 1D |
| **** | <0.0001 | 1D |
| * | 0.0143 | 1D |
| *** | 0.001 | 2D |
| ns | >0.9999 | 2D |
| ns | >0.9999 | 2D |
| ns | >0.9999 | 2D |
| ns | >0.9999 | 2D |
| ns | >0.9999 | 2D |
| ns | >0.9999 | 2D |

| | | |
|---|---|---|
| ns | >0.9999 | 2D |
| **** | <0.0001 | 2D |
| **** | <0.0001 | 2D |
| ns | >0.9999 | 2E |
| ns | >0.9999 | 2E |
| ns | >0.9999 | 2E |
| ns | >0.9999 | 2E |
| ns | >0.9999 | 2E |
| ns | >0.9999 | 2E |
| ns | >0.9999 | 2E |
| ** | 0.0077 | 2E |
| **** | <0.0001 | 2E |
| **** | <0.0001 | 2E |
| **** | <0.0001 | 7A |
| **** | <0.0001 | 7A |
| *** | 0.0006 | 7A |
| * | 0.0129 | 7A |
| ns | 0.9881 | 7A |
| ns | 0.6924 | 7A |
| * | 0.0454 | 7A |
| ns | 0.4196 | 7A |
| * | 0.0188 | 7A |
| ns | 0.3789 | 7A |
| * | 0.0217 | 7B |

| | | |
|---|---|---|
| ** | 0.5406 | 7B |
| ns | 0.9993 | 7B |
| ns | 0.0034 | 7B |
| ns | 0.8196 | 7B |
| ns | 0.2487 | 7B |
| ** | 0.008 | 7B |
| * | 0.0405 | 7B |
| ** | 0.0011 | 7B |
| ns | 0.3426 | 7B |
| ns | 0.5355 | 7C |
| ns | 0.9914 | 7C |
| * | 0.0161 | 7C |
| * | 0.0156 | 7C |
| ns | 0.2442 | 7C |
| ns | 0.1936 | 7C |
| ns | 0.1877 | 7C |
| ** | 0.0038 | 7C |
| ** | 0.0037 | 7C |
| ns | >0.9999 | 7C |
| ns | 0.97 | 7D |
| ns | 0.2898 | 7D |
| ns | 0.0891 | 7D |
| ns | 0.9141 | 7E |
| **** | <0.0001 | 7E |

| | | |
|---|---|---|
| **** | <0.0001 | 7E |
| ns | 0.9902 | 7F |
| ns | 0.181 | 7F |
| ns | 0.0736 | 7F |
| **** | <0.0001 | 7G |
| ns | 0.9329 | 7G |
| **** | <0.0001 | 7G |
| ns | 0.8572 | 7H |
| ns | 0.1897 | 7H |
| ns | 0.9959 | 7H |
| ** | 0.0012 | 7H |
| ns | 0.9996 | 7H |
| ns | 0.7967 | 7H |
| ns | 0.9856 | 7H |
| * | 0.0183 | 7H |
| ns | 0.9563 | 7H |
| ns | 0.4133 | 7H |
| ns | 0.2451 | 7H |
| ns | 0.3133 | 7H |
| ** | 0.0039 | 7H |
| ns | >0.9999 | 7H |
| ** | 0.0024 | 7H |
| ** | 0.0047 | S1B |
| ** | 0.0085 | S1B |

| | | |
|---|---|---|
| ** | 0.003 | S1B |
| ns | 0.7046 | S1B |
| ns | 0.9999 | S1B |
| ns | 0.855 | S1B |
| ns | 0.9842 | S1B |
| ns | 0.0849 | S1B |
| * | 0.022 | S1B |
| ns | 0.5127 | S1B |
| ns | 0.2928 | S1B |
| ns | 0.0718 | S1B |
| * | 0.0382 | S1B |
| * | 0.0105 | S1B |
| ns | 0.8178 | S1B |
| **** | <0.0001 | S1C |
| **** | <0.0001 | S1C |
| **** | <0.0001 | S1C |
| ns | 0.0974 | S1C |
| ns | 0.1229 | S1C |
| **** | <0.0001 | S1C |
| *** | 0.0001 | S1C |
| ** | 0.0061 | S1C |
| **** | <0.0001 | S1C |
| ns | 0.8852 | S1C |
| **** | <0.0001 | S1C |

| | | |
|---|---|---|
| **** | <0.0001 | S1C |
| **** | <0.0001 | S1C |
| **** | <0.0001 | S1C |
| ** | 0.0018 | S1C |
| **** | <0.0001 | S3B |
| **** | <0.0001 | S3B |
| **** | <0.0001 | S3B |
| **** | <0.0001 | S3B |
| **** | <0.0001 | S3B |
| ns | 0.7123 | S3B |
| ns | 0.9325 | S3B |
| ns | 0.9987 | S3B |
| ns | 0.9986 | S3B |
| ns | 0.1958 | S3B |
| ns | 0.4686 | S3B |
| ns | 0.4667 | S3B |
| ns | 0.9935 | S3B |
| ns | 0.9937 | S3B |
| ns | >0.9999 | S3B |
| ns | 0.3409 | S8A |
| ns | 0.9781 | S8A |
| ns | 0.9726 | S8A |
| ns | 0.8181 | S8A |
| ns | 0.073 | S8A |

| | | |
|---|---|---|
| ns | 0.6174 | S8A |
| ns | 0.0807 | S8A |
| ns | 0.3041 | S8A |
| ns | 0.6535 | S8A |
| ns | 0.9875 | S8A |
| ns | 0.8137 | S8A |
| ns | 0.9955 | S8A |
| ns | 0.0542 | S8A |
| * | 0.0197 | S8A |
| ** | 0.0019 | S8A |
| ns | 0.9989 | S8A |
| ns | 0.7372 | S8A |
| ns | 0.9399 | S8A |
| **** | <0.0001 | S8A |
| **** | <0.0001 | S8A |
| **** | <0.0001 | S8A |
| ns | >0.9999 | S8A |
| ns | >0.9999 | S8A |
| ns | >0.9999 | S8A |
| ns | 0.4718 | S8A |
| ns | 0.8011 | S8A |
| ns | 0.6882 | S8A |
| ns | 0.9973 | S8A |
| ns | 0.9998 | S8A |

| | | |
|---|---|---|
| ns | >0.9999 | S8A |
| **** | <0.0001 | S8A |
| **** | <0.0001 | S8A |
| **** | <0.0001 | S8A |
| ns | 0.9354 | S8A |
| ns | 0.8806 | S8A |
| ns | >0.9999 | S8A |
| ns | 0.9877 | S8A/B |
| ns | 0.9884 | S8A/B |
| ns | 0.9162 | S8A/B |
| ns | 0.0572 | S8A/B |
| ns | >0.9999 | S8A/B |
| ns | >0.9999 | S8A/B |
| ns | 0.9998 | S8A/B |
| ns | 0.6466 | S8A/B |
| ns | 0.9806 | S8A/B |
| ns | 0.0801 | S8A/B |
| ns | >0.9999 | S8A/B |
| ns | 0.9994 | S8A/B |
| * | 0.0276 | S8A/B |
| ns | >0.9999 | S8A/B |
| ns | >0.9999 | S8A/B |
| * | 0.0201 | S8A/B |
| ns | >0.9999 | S8A/B |

| | | |
|---|---|---|
| ns | >0.9999 | S8A/B |
| ns | 0.3919 | S8A/B |
| ns | 0.4723 | S8A/B |
| ns | 0.387 | S8A/B |
| * | 0.015 | S8A/B |
| ns | 0.9963 | S8A/B |
| ns | 0.9545 | S8A/B |
| ns | 0.9682 | S8A/B |
| ns | 0.7988 | S8A/B |
| ns | 0.2319 | S8A/B |
| ns | 0.5494 | S8A/B |
| ns | 0.9942 | S8A/B |
| ns | 0.9991 | S8A/B |
| ns | 0.369 | S8A/B |
| ns | 0.2788 | S8A/B |
| ns | >0.9999 | S8A/B |
| ns | 0.9061 | S8A/B |
| ns | 0.5319 | S8A/B |
| ns | 0.995 | S8A/B |
| ns | >0.9999 | S8A/B |
| ns | 0.9993 | S8A/B |
| * | 0.0453 | S8A/B |
| ns | 0.0774 | S8A/B |
| ns | >0.9999 | S8A/B |

| | | |
|---|---|---|
| ns | >0.9999 | S8A/B |
| ns | >0.9999 | S8A/B |
| ns | 0.9987 | S8A/B |
| ns | 0.0882 | S8A/B |
| ns | 0.0587 | S8A/B |
| ns | >0.9999 | S8A/B |
| ns | >0.9999 | S8A/B |
| ns | 0.9228 | S8A/B |
| ns | 0.8214 | S8A/B |
| ** | 0.0019 | S8A/B |
| ** | 0.0013 | S8A/B |
| ns | >0.9999 | S8A/B |
| ns | >0.9999 | S8A/B |
| ns | 0.094 | S8B |
| ns | >0.9999 | S8B |
| ns | >0.9999 | S8B |
| ns | 0.1136 | S8B |
| ns | 0.1482 | S8B |
| ns | >0.9999 | S8B |
| ns | >0.9999 | S8B |
| ns | 0.381 | S8B |
| ns | 0.6714 | S8B |
| ns | 0.2968 | S8B |
| ns | 0.5675 | S8B |

| | | |
|---|---|---|
| ns | 0.9987 | S8B |
| * | 0.0252 | S8B |
| *** | 0.0005 | S8B |
| *** | 0.0003 | S8B |
| ns | 0.5783 | S8B |
| ns | 0.4654 | S8B |
| ns | >0.9999 | S8B |
| * | 0.039 | S8B |
| **** | <0.0001 | S8B |
| **** | <0.0001 | S8B |
| * | 0.0486 | S8B |
| * | 0.0321 | S8B |
| ns | >0.9999 | S8B |
| ns | 0.6955 | S8B |
| ns | 0.6251 | S8B |
| ns | 0.7645 | S8B |
| * | 0.0473 | S8B |
| ns | 0.0769 | S8B |
| ns | >0.9999 | S8B |
| * | 0.0441 | S8B |
| **** | <0.0001 | S8B |
| **** | <0.0001 | S8B |
| ** | 0.0017 | S8B |
| *** | 0.0009 | S8B |

| | | |
|---|---|---|
| ns | >0.9999 | S8B |
| **** | <0.0001 | S8C |
| **** | <0.0001 | S8C |
| **** | <0.0001 | S8C |
| *** | 0.0006 | S8C |
| ns | 0.9965 | S8C |
| ns | 0.3796 | S8C |
| * | 0.0238 | S8C |
| ns | 0.5702 | S8C |
| * | 0.0452 | S8C |
| ns | 0.4884 | S8C |
| ns | 0.9893 | S8D |
| ns | 0.8911 | S8D |
| ns | 0.0629 | S8D |
| ** | 0.0066 | S8D |
| ns | 0.592 | S8D |
| ns | 0.0999 | S8D |
| ** | 0.0089 | S8D |
| ** | 0.0066 | S8D |
| *** | 0.0006 | S8D |
| ns | 0.6845 | S8D |
| * | 0.0347 | S8E |
| * | 0.0239 | S8E |
| *** | 0.0009 | S8E |

| | | |
|---|---|---|
| ** | 0.0012 | S8E |
| **** | <0.0001 | S8E |
| **** | <0.0001 | S8E |
| **** | <0.0001 | S8E |
| ns | 0.902 | S8E |
| ns | 0.8982 | S8E |
| ns | >0.9999 | S8E |
| ns | 0.664 | S8F |
| ns | 0.998 | S8F |
| ns | 0.9916 | S8F |
| ns | 0.9539 | S8F |
| ns | 0.8834 | S8F |
| ns | 0.9559 | S8F |
| ns | 0.9853 | S8F |
| ns | >0.9999 | S8F |
| ns | 0.9979 | S8F |
| ns | >0.9999 | S8F |
| ns | 0.888 | S8G |
| ns | 0.9994 | S8G |
| ns | 0.8804 | S8G |
| ns | 0.9914 | S8G |
| ns | 0.7279 | S8G |
| ns | >0.9999 | S8G |
| ns | 0.996 | S8G |

| | | |
|---|---|---|
| ns | 0.7276 | S8G |
| ns | 0.9408 | S8G |
| ns | 0.9936 | S8G |
| X | 0.9812 | S8H |
| ns | >0.9999 | S8H |
| ns | 0.9815 | S8H |
| ns | >0.9999 | S8H |
| ns | 0.9398 | S8H |
| ns | >0.9999 | S8H |
| ns | 0.9398 | S8H |
| ns | 0.9403 | S8H |
| ns | >0.9999 | S8H |
| ns | 0.9403 | S8H |
